# CellDiffusion: Enhancing scRNA-seq data integration with accurate inference of cell-cell association

**DOI:** 10.1101/2024.01.28.577667

**Authors:** Yu-Chen Liu, Ziying Huang, Simon Liang Lu, Lei Jiang, Zedong Lin, Jou-Hsuan Lee, Vijaya B. Kolachalama, Rong Li, Dong Xu, Juexin Wang, Heng Pan, Chao Zhang

## Abstract

Integrating single-cell transcriptomic datasets across laboratories, conditions, and technologies remains challenging due to cell-type-specific batch effects and heterogeneous biological compositions. Current methods often fail to separate genuine biological variation from technical noise, misaligning distinct cell populations or leaving batch effects uncorrected. To address these challenges, we present CellDiffusion, an unsupervised graph-based model that propagates cellular features through a diffusion process to strengthen meaningful cell-cell associations in data integration. By learning a cell-cell attention network, CellDiffusion adaptively identifies corresponding populations across batches. Our approach applies cell-specific correction in data integration, generating cell embeddings that align cells across batches while preserving intrinsic cell types and developmental trajectories. We validate CellDiffusion across various datasets, including the human endometrium, one of the most dynamic and complex tissues featuring extensive cellular and molecular changes with the menstrual cycle. This framework also improves other essential analyses such as label transfer, clustering, and trajectory inference. CellDiffusion represents a powerful paradigm for accurate single-cell transcriptomic integrative analysis.

**Teaser:** A diffusion-based framework removes technical noise and preserves biological variation in single-cell datasets.

## Introduction

Single-cell RNA sequencing (scRNA-seq) has substantially advanced our understanding of cellular heterogeneity, providing unprecedented insights into development, disease progression, and tissue dynamics(*1–5*). As recent large-scale comprehensive studies increasingly rely on comparing single-cell datasets from diverse sources, such as different individuals, conditions, cohorts, or platforms, data integration has effectively become critical in scRNA-seq analysis.

In scRNA-seq data integration, cell-type-specific batch effect removal is one of the most crucial challenges(*6, 7*), which is prevalent in datasets containing various cell types that may respond differently to dissociation protocols, reagents, and degradations. Therefore, optimized integration must account for this cell-type-specific technical noise to avoid distorting the underlying biological signal. However, existing integration tools are unable to handle such a complicated scenario, which may lead to improper batch correction. For instance, fastMNN(*7*), Scanorama(*8*), and ComBat(*9*) often fail to align identical cell types from different batches, suffering from the downalignment problem. Meanwhile, other methods, such as Harmony(*10*), Seurat(*11*), and scVI(*12*), may align biologically distinct cell types together due to overcorrection, which was named as misalignment in this manuscript. The suboptimal integration will introduce severe artifacts that distort cell states, intercellular relationships, and ultimately the biological interpretation.

Furthermore, real-world datasets often contain inter-sample or inter-batch imbalances in addition to batch effects(*13*), indicating critical biological variations across samples and batches, such as unequal cell compositions and unmatched cell types. The imbalance comprises underlying biological signals in plenty of comparative studies relevant to embryonic development, cancer evolution, and autoimmune diseases(*14–16*). The co-occurrence of data imbalances with batch effects significantly increases the risk of misaligning distinct cell types across batches in data integration. Although the importance of addressing data imbalance in integration is well recognized(*13, 17, 18*), there are still no existing tools that consistently and effectively account for biological imbalances across batches. By constructing experimental design constraints, the recent study, CellANOVA(*19*), has made significant progress in recovering biological signals, that were often falsely removed due to overcorrection and misalignment. However, it relies on user-defined control pools to discriminate batch effects from biological signals, which introduces subjectivity and limits its generalizability in studies lacking well-defined controls or sufficient control samples. This gap underscores the need for integration methods that can accurately and spontaneously disentangle meaningful biological variations from complex technical artifacts across diverse datasets.

Accurately modeling cell-cell associations across batches is indispensable to optimized batch effect removal and data integration. Nearly every downstream task, including unsupervised clustering, query-to-reference mapping, and trajectory inference, also depends on the precise inference of intercellular relationships(*3, 20, 21*). Within each sample, cell-cell associations are reflected in the hierarchical organization of cell populations, spanning biological lineages, broad cell types, and fine-grained subtypes. Recent studies emphasize that robust biological analyses rely on cell embeddings that preserve this hierarchical cell ontology(*22*). However, when integrating data across batches, these hierarchical structures can be distorted by batch effects and further complicated by imbalanced cell compositions. Consequently, many existing methods fail to capture inter-batch cell-cell associations in data integration reliably.

Recent studies of graph neural diffusion methods(*23, 24*) present novel insights in modeling graph node associations. These models simulate a continuous diffusion process on graphs, treating the diffusion network as the discretization of an underlying partial differential equation, where the model architecture and graph topology correspond to the discretization of temporal and spatial operators. Graph neural diffusion methods learn node-node relationships and model graph topological structures by capturing the dynamic flow of information across nodes. Further, graph neural diffusion models employ an attention mechanism(*25, 26*) to guide the information diffusion process on graphs, improving the accuracy in association modeling. With their strong mathematical foundations and physically inspired structures, graph neural diffusion models have great potential for addressing complex cell-cell associations in scRNA-seq data integration under challenging scenarios.

Inspired by the concept of graph neural diffusion, we developed CellDiffusion, a graph-based diffusion model that can infer cell-cell associations accurately, handle complex data structures, and disentangle biological signals from technical variations to enhance scRNA-seq integration. Our model not only exhibits optimized batch effect removal, even in imbalanced and heterogeneous datasets, but also preserves the hierarchical organization of cell populations. It enhances downstream analyses such as unsupervised clustering, query-to-reference mapping, and trajectory inference, which also highly depend on the precise inference of intercellular relationships(*3, 20, 21*). Across multiple benchmarks, including dynamic human endometrium and bone marrow datasets, CellDiffusion outperformed existing tools, preserving unique cell identities and biological trajectories while effectively correcting for batch effects. Our model provides a versatile and accessible framework for robust integrative analysis of scRNA-seq data, paving the way towards the large-scale comprehensive scRNA-seq study across different institutes, populations, and sequencing platforms.

## Results

### CellDiffusion disentangles biological variations from batch effects via accurate inference of cell-cell associations

Real-world data is extremely complicated due to the simultaneous presence of batch effects, imbalanced cell-type compositions, and unmatched cell types across samples and batches (**Fig. 1A**). Overlooking inherent biological heterogeneity can lead to faulty batch correction in data integration. Many existing methods rely on “anchors” to match cells across batches, while they often struggle to identify reliable “anchors” due to massive unknown confounders. This can result in two critical integration pitfalls: downalignment, when identical cells fail to align due to incomplete batch correction, and misalignment, when distinct cells are merged due to overcorrection (**Fig. 1B**). The latter issue is more severe in Harmony(*10*), the most popular integration tool, as well as other state-of-the-art methods(*19, 27*). For instance, the human endometrium undergoes dynamic changes in cell composition and gene expression along with the menstrual cycle, which introduces a dramatic biological imbalance into the endometrial scRNA-seq dataset where samples from different menstrual cycles harbor different subtypes of stromal cells(*28*). Harmony falsely merged stromal cells in different statuses (**Fig. 1C**), obscuring lineages of stromal cells and preventing further exploration of biological signals relevant to menstrual cycles. In contrast, CellDiffusion demonstrates a unique ability to disentangle biological variation from underlying batch effects. It successfully resolved stromal cell states across cycles while avoiding both downalignment and misalignment simultaneously (**Fig. 1C**), preserving the biologically meaningful lineages of endometrial stromal cells.

**Fig. 1.**
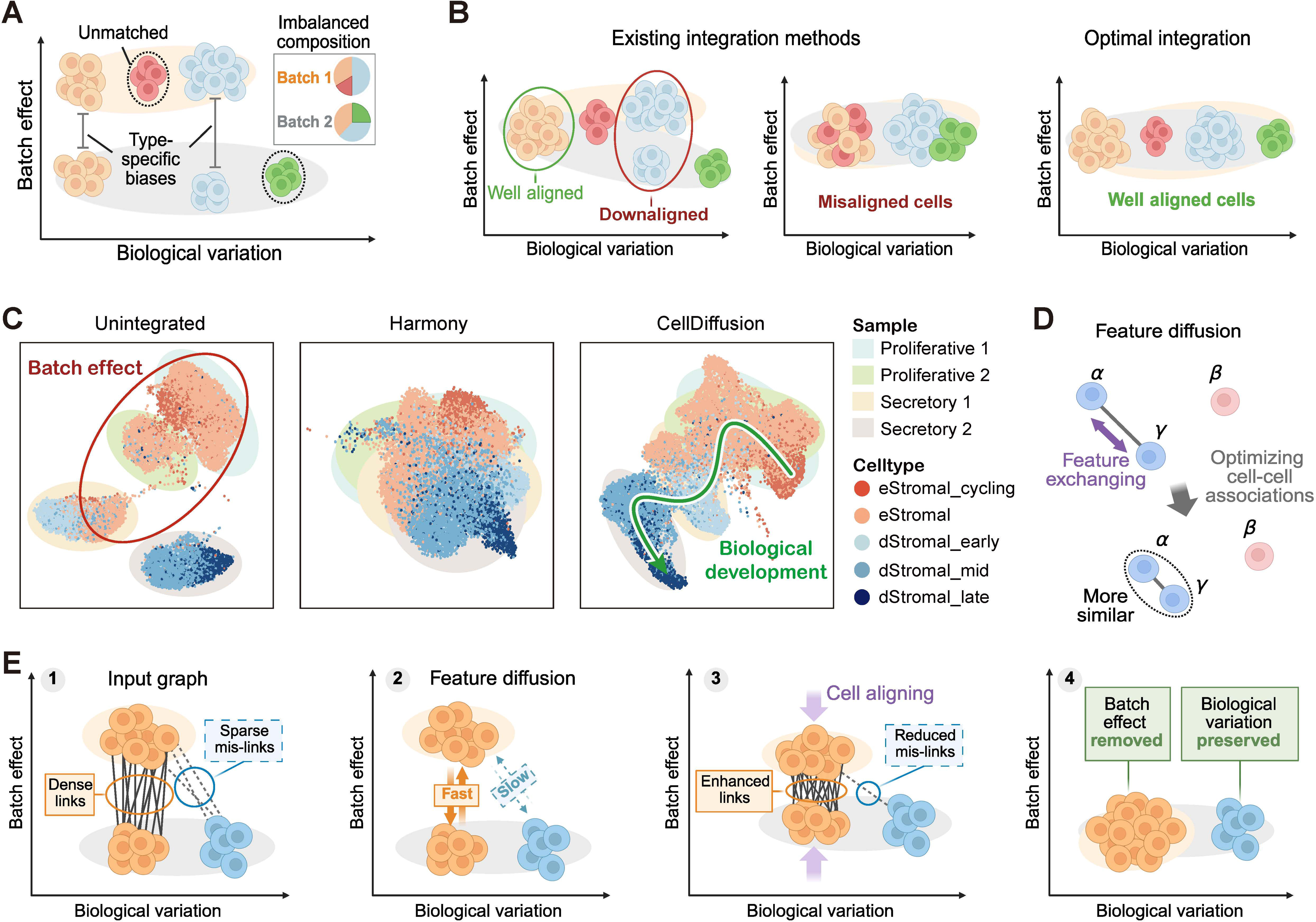
CellDiffusion distinguishes biological variations from batch effects. (**A**) Real-world scRNA-seq data from different cohorts contain interwoven batch effects and biological variations, such as cell type-specific biases, imbalanced cell compositions, and unmatched cell types across batches. (**B**) Compared with ideal data integration, existing tools often suffer from under- and over-batch correction, incorrectly splitting identical cell types and misaligning distinct cell identities. (**C**) An example using clinical samples demonstrates that Harmony misaligned distinct stromal cell types, whereas CellDiffusion preserved their biological differences during integration. (**D**) In CellDiffusion, feature diffusion through linked cells enhances associations among cells with similar biological identities. (**E**) Inter-batch diffusion aligns the same cell types across batches while preserving distinctions between different biological states.

CellDiffusion is a graph-based framework that models intercellular relationships by constructing a cell-cell network and propagating transcription-derived features across connected cells. The diffusion process gradually homogenizes connected cells, enabling biologically coherent alignment (**Fig. 1D**; **figs. S1-S2**). By performing diffusion across batches, CellDiffusion facilitates feature exchanges between similar cells from different batches and aligns them together, thereby removing batch effects and integrating data simultaneously. CellDiffusion initiates with an Optimized Mutual Nearest Neighbor (OMNN) algorithm to connect cells across batches, preferentially linking putatively matched cell types (See **Materials and Methods** for details). To avoid downalignment caused by missing connections between cells belonging to the same subgroup, OMNN tends to connect biologically corresponding cells across batches with dense edges, which may occasionally introduce incorrect connections between different cell types. CellDiffusion can tolerate such incorrect connections in the graph since they are expected to be sparser relative to correct biological linkages. CellDiffusion leverages diffusion dynamics to intrinsically attenuate weak associations. Strongly linked cells rapidly exchange features while sparse and imprecise connections between unrelated cells experience minimal diffusion (**Fig. 1E**), thereby avoiding possible misalignment. To further ensure accurate cell alignment, CellDiffusion employs an attention mechanism(*25, 26*) to guide and optimize the feature propagation through all diffusion layers in the latent space (**fig. S1A**; **fig. S2B-C**). This mechanism allows CellDiffusion to enhance meaningful inter-batch correspondences, suppress misleading associations, align the same cell types across batches, and maintain biological distinctions, in a cell-specific manner.

To retain the original biological structure of the hierarchical cell ontology (**fig. S2D**), CellDiffusion reconstructs a K-nearest neighbor (KNN) graph within each batch following diffusion (**fig. S1B**), enforcing structural preservation through a loss function that compares reconstructed KNN adjacencies with their input counterparts before diffusion. This strategy prevents mis-relocation of populations within each batch while permitting the cross-batch alignment. Accordingly, CellDiffusion can preserve the original biological information during integration, demonstrating its advantage in handling real-world complexities, particularly when batches contain unmatched populations or imbalanced cell compositions (**Fig. 1C**).

### CellDiffusion preserves menstrual stromal lineages when integrating human endometrial samples

The endometrium, a steroid hormone-responsive mucosal layer lining the uterine cavity, orchestrates cyclic tissue remodeling throughout the menstrual cycle in reproductive-age females (**Fig. 2A**). This inherent dynamism presents significant challenges for scRNA-seq analysis, as endometrial tissues frequently exhibit pronounced imbalances in cellular compositions and contain distinct cell populations across samples of different menstrual stages(*29*). Using stromal fibroblasts as an example, they undergo phase-specific functional transitions. Proliferative stromal cells (eStromal cells) mediate tissue restructuring through enhanced extracellular matrix (ECM) biosynthesis, whereas decidualized stromal cells (dStromal cells) in the secretory stage establish a specialized microenvironment via paracrine secretion of WNT antagonists (particularly DKK1) that coordinate epithelial glandular maturation(*30*) (**Fig. 2A**). Spatial transcriptomics data further reveal distinct spatial distribution of stromal subtypes along different menstrual stages (**Fig. 2B**), highlighting their biological authenticity and divergence. The functional dichotomy between eStromal and dStromal cells underscores the critical need for precise cellular stratification, which enables the accurate reconstruction of endometrial remodeling trajectories and the identification of phase-specific signals in different physiological and pathological contexts. Failing to discriminate between eStromal and dStromal cells would introduce menstrual confounders into the downstream analysis, mistakenly attributing expression changes derived from different menstrual stages and tissue compositions to endometrial disorders such as endometrial aging and recurrent implantation failure. Therefore, we employed this endometrial dataset as a biologically complex real-world scenario to evaluate CellDiffusion.

**Fig. 2.**
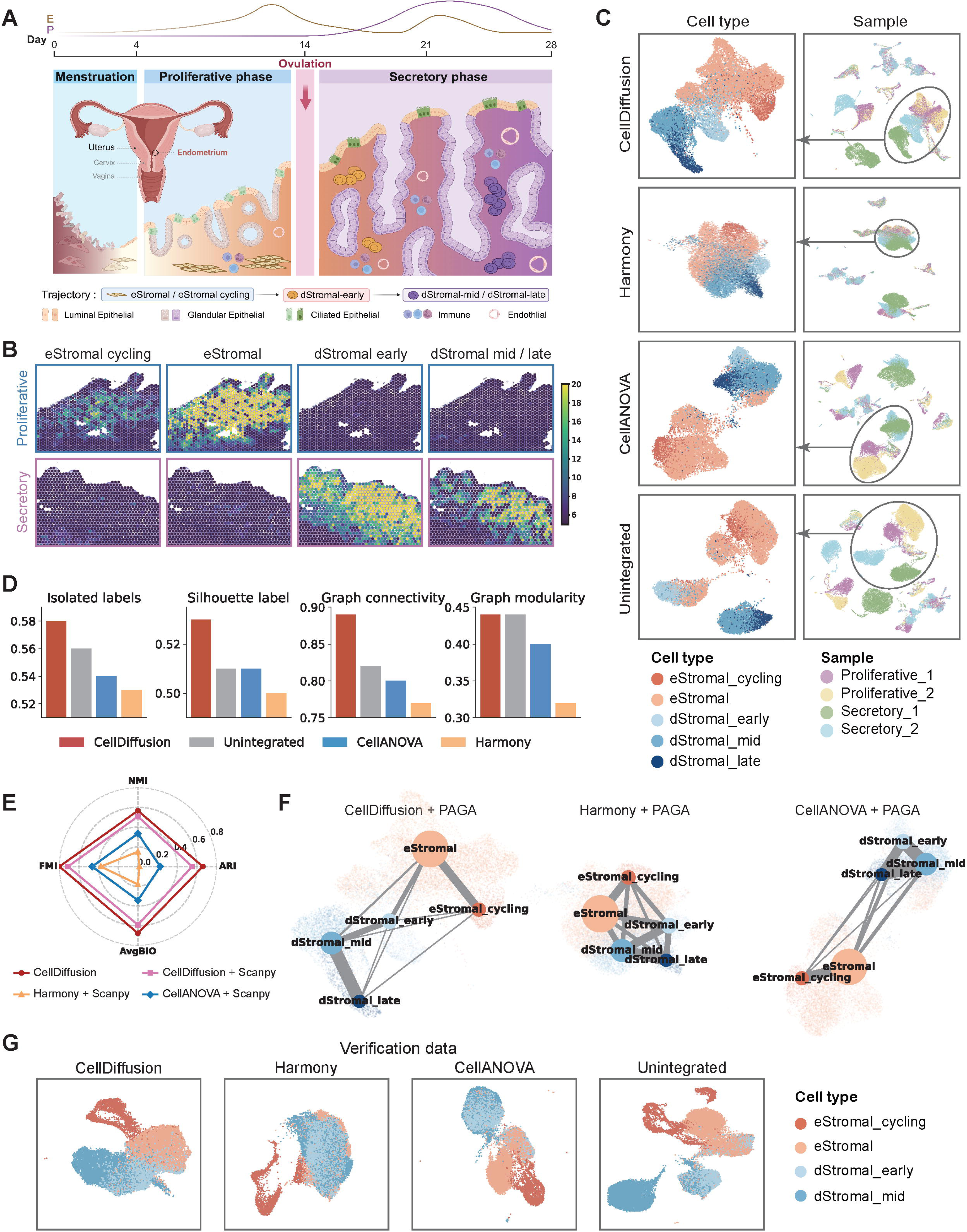
Data integration of human endometrium datasets with both batch effects and cell type imbalance. (**A**) Schematic illustration of the dynamic cell composition and morphological changes in the human endometrium across the menstrual cycle. Created in BioRender. Huang, Z. https://BioRender.com/fynd2c1 (2026). (**B**) Different stromal cell subtypes exhibit distinct spatial distribution within the functional layers and appear only in specific phases. (**C**) CellDiffusion outperforms Harmony and CellANOVA in preserving biological integrity during integration of the endometrium samples from different phases. The stromal cell population is highlighted as the example in UMAP plots. (**D**) Four different metrics to quantitatively evaluate the integration performance of different methods on the stromal cell population. (**E**) Evaluation of clustering results for each method applied to the stromal cell population. (**F**) PAGA trajectory inference is applied to the stromal subtypes based on integration results of each method. CellDiffusion better preserves the cell fate transition trajectory consistent with current biological knowledge compared to Harmony and CellANOVA. (**G**) UMAP visualizations of integration results on endometrium samples from an independent cohort. Abbreviations: E, estrogen; P, progesterone. “Unintegrated” denotes the common preprocessed cell representation before applying any batch-correction or integration method.

We randomly selected two proliferative and two secretory samples from a recently published high-quality human endometrial study (Endometrium-35k)(*28*). Without integration, strong batch effects drove sample-specific grouping despite the presence of common cell types across samples (**Fig. 2C**; **fig. S3**). After applying CellDiffusion, all samples were effectively integrated. Matching cells across samples were correctly grouped, and inherent biological variations were properly retained, even in imbalanced stromal subtypes. In contrast, Harmony(*10*) failed to resolve stromal heterogeneity, showing extensive overcorrection of stromal cells, while CellANOVA(*19*), using secretory samples as a control pool, misaligned stromal cells in secretory samples and failed to integrate eStromal cells in proliferative samples (**Fig. 2C**). Alternative control pool configurations were also tested but did not improve the performance (**fig. S4**). We further employed multiple metrics to quantify the batch correction performance. Most commonly used evaluation metrics of the batch correction(*27*) were not designed to handle samples with highly imbalanced cell compositions and unmatched cell types, as in the Endometrium-35k dataset. We thus selected four appropriate label-based metrics that ensure impartiality to evaluate imbalanced datasets, including isolated labels, silhouette label, graph connectivity, and graph modularity (**Supplementary Text 1** and **Materials and Methods**). CellDiffusion consistently outperformed Harmony and CellANOVA in all metrics (**Fig. 2D**). These observations confirmed the superior performance of CellDiffusion in data integration, removing potential batch effects without affecting underlying biological signals.

### CellDiffusion enhances downstream analysis by providing a well-calibrated integration output of human endometrial samples

Data integration critically impacts downstream analyses such as clustering and trajectory inference. To fairly benchmark the direct impact of different data integration approaches, we applied an identical downstream analysis pipeline to the integration output of each approach.

First, we conducted clustering using Scanpy(*31*) to rediscover five stromal clusters. To evaluate the performance, we computed an aggregated metric, AvgBio, defined as the mean of adjusted Rand index (ARI)(*32*), normalized mutual information (NMI)(*33*), and the Fowlkes-Mallows index (FMI)(*34*) (**Supplementary Text 2**). Of note, CellDiffusion outperformed other methods under this standardized clustering pipeline (**Fig. 2E**). Moreover, when employing its built-in attention-based clustering approach, CellDiffusion achieved the highest score of all evaluation metrics, underscoring its effectiveness for capturing biologically meaningful cell populations (**Fig. 2E**). To further confirm the biological representation of clusters derived from different integration approaches, we compared gene expression of known cell-type-specific markers in clusters of each integration method (**fig. S5**). CellDiffusion nearly recapitulated marker gene expression patterns observed in the manually curated dataset(*28*) (**fig. S5**). The misalignment of Harmony caused three dStromal subclusters being indistinguishable. The cluster-4 of CellANOVA expressed markers from both dStromal and eStromal cells, which is absent in curated annotations. We further identified cluster-specific markers for each method and performed pathway enrichment analysis. CellDiffusion consistently determined biologically relevant functions that aligned with manually annotated cell types. In contrast, both Harmony and CellANOVA yielded largely irrelevant functions due to misalignment of cells from distinct cell types (**fig. S6**).

Second, we examined the continuous trajectory of stromal subtypes, which spans both proliferative and decidualized stages. Trajectory inference using PAGA(*21*) on integrated data of different approaches revealed that CellDiffusion properly preserved the underlying menstrual continuum, whereas both Harmony and CellANOVA obscured key transitions due to distortion of biologically relevant structure during integration (**Fig. 2F**).

### CellDiffusion is robust in data integration upon different sample selections and cohorts in the human endometrium

To test the robustness of our framework, we applied CellDiffusion to the full dataset of Endometrium-35k(*28*) (Endometrium-314k, 313,527 cells from 87 samples) and our in-house curated dataset (Endometrium-EV). Endometrium-314k was used to test whether sample selection would distort the performance and whether CellDiffusion applies to a large-scale dataset. Of note, CellDiffusion produced a more coherent and biologically meaningful integration compared to other methods, in a fast and scalable manner. Besides stromal cells, CellDiffusion also demonstrated a strong capability in distinguishing epithelial cells, another important functional group of endometrial components (**fig. S7**). Endometrium-EV was designed to validate the generalization of CellDiffusion across different cohorts, containing two proliferative and two secretory samples as Endometrium-35k. To enable comparable cell compositions, marker genes of Endometrium-314k(*28*) were used to annotate Endometrium-EV via Seurat(*20*). After manual curation, ten different cell types were identified in Endometrium-EV. Similarly, CellDiffusion effectively aligned common stromal cells across samples while preserving biologically distinct populations, outperforming both Harmony and CellANOVA (**Fig. 2G**; **fig. S8A**). Harmony and CellANOVA exhibited heavy misalignment, failing to distinguish stromal subtypes, which expressed distinct markers from each other **(fig. S8B)**. Similarly, CellDiffusion achieved the highest scores compared to other methods in the Endometrium-EV dataset in terms of the evaluation of batch effect removal and downstream clustering **(fig. S8C-D)**. Thus, we concluded that CellDiffusion provides a fast, accurate, and generalized deep learning framework for correcting batch effects in a real-world, complex clinical scenario, thereby enhancing downstream scRNA-seq analysis, including clustering and trajectory inference.

### CellDiffusion retains functionally different cell types with subtle transcriptional differences in the scRNA-seq of human bone marrow mononuclear cells

Although the endometrium datasets reflect a real-world scenario with complex, continuous cell transitions and imbalanced cell populations, most existing integration methods are not designed to address this challenge. Also, a systematic evaluation standard has not been well-established for such clinical settings. To enable extensive and fair comparison with extant approaches, we employed a gold-standard dataset of bone marrow mononuclear cells (BMMC)(*35*), comprising 40 distinct cell types from four healthy human donors. This dataset rigorously tests whether an integration method can preserve cells in different immune states featuring similar gene expression profiles, which are critical for understanding the antibody diversification and immune spectrum. CellDiffusion achieved better integration performance compared with benchmarked methods, including Harmony(*10*), Seurat(*11*), and ComBat(*9*) (**Fig. 3A**; **fig. S9A**). CellANOVA could not be applied due to a lack of appropriate control pools. CellDiffusion successfully distinguished immunoglobulin kappa constant-positive (IGKC+) and -negative (IGKC-) subgroups within B cells, which were erroneously merged by all the other methods. IGKC encodes the constant domain of kappa-type light chains, contributing to antibody production in B cells. IGKC+ and IGKC-B cells displayed distinct gene expression profiles, marked by specific signature genes, indicating functional heterogeneity in healthy immune systems **(fig. S9B)**. CellDiffusion could also distinguish IGKC +/- subpopulations within plasma cells and plasmablasts. In contrast, these biologically distinct subpopulations were obscured to a different extent by other methods, due to potential overcorrection **(fig. S9A)**. The ability to preserve such subtle but biologically meaningful differences highlights the strength of CellDiffusion in accurately resolving complex cellular heterogeneity while minimizing overcorrection. This provides a solid technical foundation for future applications of CellDiffusion to clinically relevant datasets, such as the tumor microenvironment, where precise delineation of immune cell states may inform the study of disease-associated cellular programs.

**Fig. 3.**
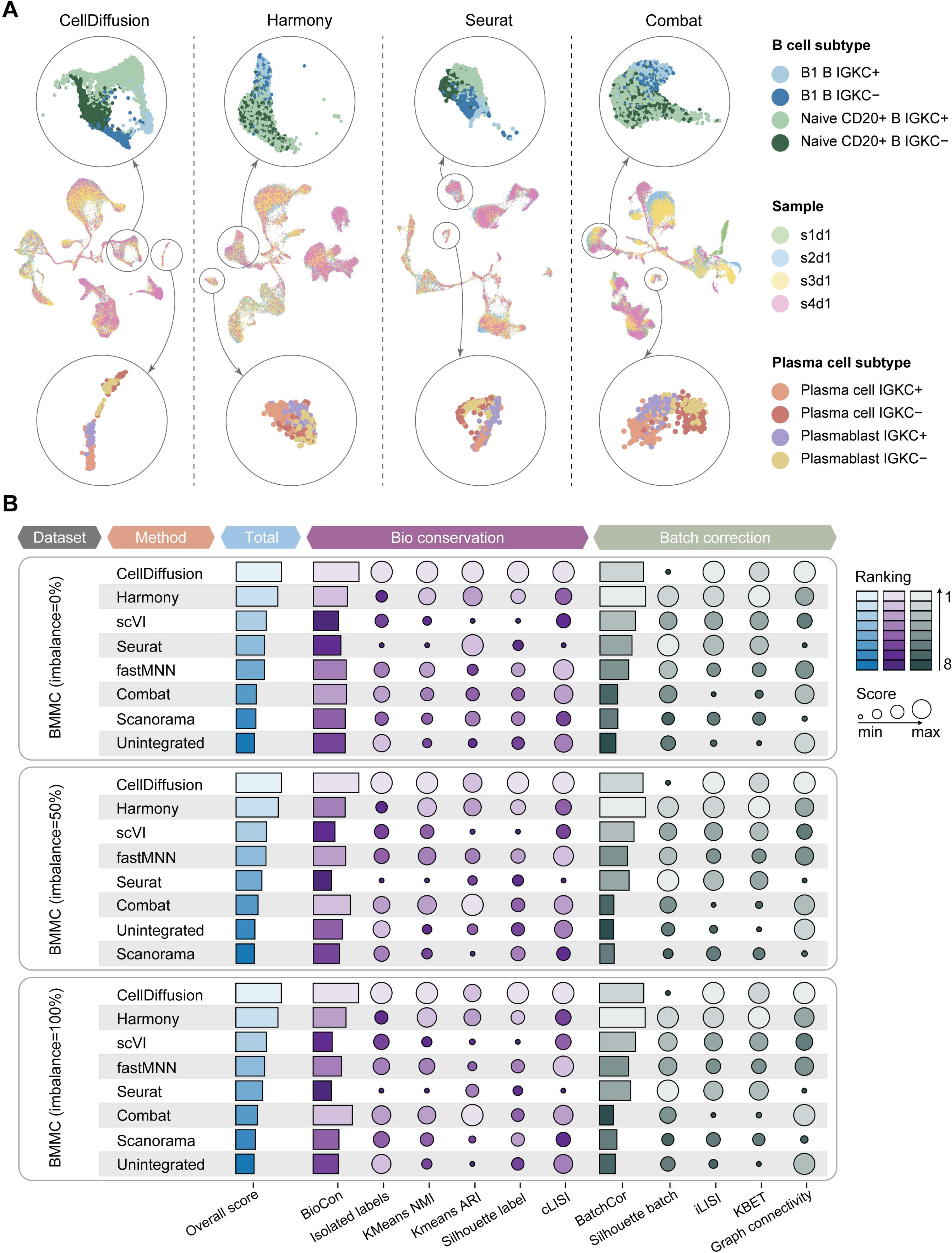
Benchmarking CellDiffusion and other integration methods on human bone marrow mononuclear cell data. (**A**) UMAP visualizations of integration results on the BMMC dataset using different methods. B cell and plasma cell populations are highlighted to better illustrate each method’s ability to distinguish subtle subtypes. (**B**) Performance of each method measured by different metrics under three conditions (‘0%’ indicates no data imbalance; ‘50%’ represents partial samples missing 50% of cells from selected cell types; and ‘100%’ indicates complete removal of selected cell types from partial samples).

Despite misaligning B cell subtypes, ComBat also failed to integrate reticulocytes properly, grouping them by sample rather than cell type, suggesting residual technical variation and under-correction (**fig. S9A**). CellDiffusion avoided both overcorrection and under-correction, achieving simultaneous integration of different cell types across samples while preserving biologically important subtypes. Collectively, we proved that CellDiffusion can implement accurate and efficient data integration across different tissue contexts.

### CellDiffusion achieves better integration performance across distinct real-world and simulated datasets compared with extant methods

To further evaluate the performance of CellDiffusion systematically, we collected two additional benchmark datasets besides BMMC, PBMC-10k (2 batches, 9 cell types)(*36*), and PBMC-23k (3 batches, 30 cell types)(*11*). Five cell type-imbalanced data scenarios with varying degrees of imbalance were introduced into each dataset by randomly selecting some cell types and removing cells within these cell types from some batches at different percentages (see **Materials and Methods** for details). This setting was used to test the integration performance across a range of imbalanced data structures representative of real-world scenarios. Six benchmark methods were selected to be evaluated with CellDiffusion. Integration performance was quantified using three aggregate metrics: BioCon (biological conservation), BatchCor (batch correction), and Total (overall integration quality)(*27*) (**Supplementary Text 1**).

In the BMMC dataset, CellDiffusion consistently achieved the highest BioCon scores across all imbalance levels in cell compositions, demonstrating superior biological fidelity (**Fig. 3B**; **fig. S10**). CellDiffusion also attained highly competitive BatchCor scores, second only to Harmony. This is consistent with previous observations that excessively high BatchCor scores may indicate overcorrection, where biological variations are erroneously removed(*13*). Harmony frequently achieved the highest BatchCor scores at the expense of suboptimized BioCon scores, indicating a loss of biological integrity due to falsely mixing distinct cell types. In terms of Total scores, CellDiffusion outperformed all benchmark methods across all tested imbalance levels, reassuring its optimal performance in data integration. The superior performance of CellDiffusion is robust in the PBMC-10k and PBMC-23k datasets (**fig. S11-S12**).

### CellDiffusion transfers accurate labels by integrating query data with references

Query-to-reference mapping is another challenge that depends on accurately inferring cell-cell associations across datasets. Precise label transfer is foundational to scRNA-seq analysis, enabling knowledge transfer from well-annotated references to newly created datasets. Existing reference-based annotation methods exhibit limited accuracy when query cells lack counterparts in references(*3, 12*) (**Fig. 4A**). Given the fact that CellDiffusion properly integrates imbalanced and unmatched cell types across batches, we hypothesized that our framework can correctly detect query cells that are rare or missing in the reference dataset (**Fig. 4B**).

**Fig. 4.**
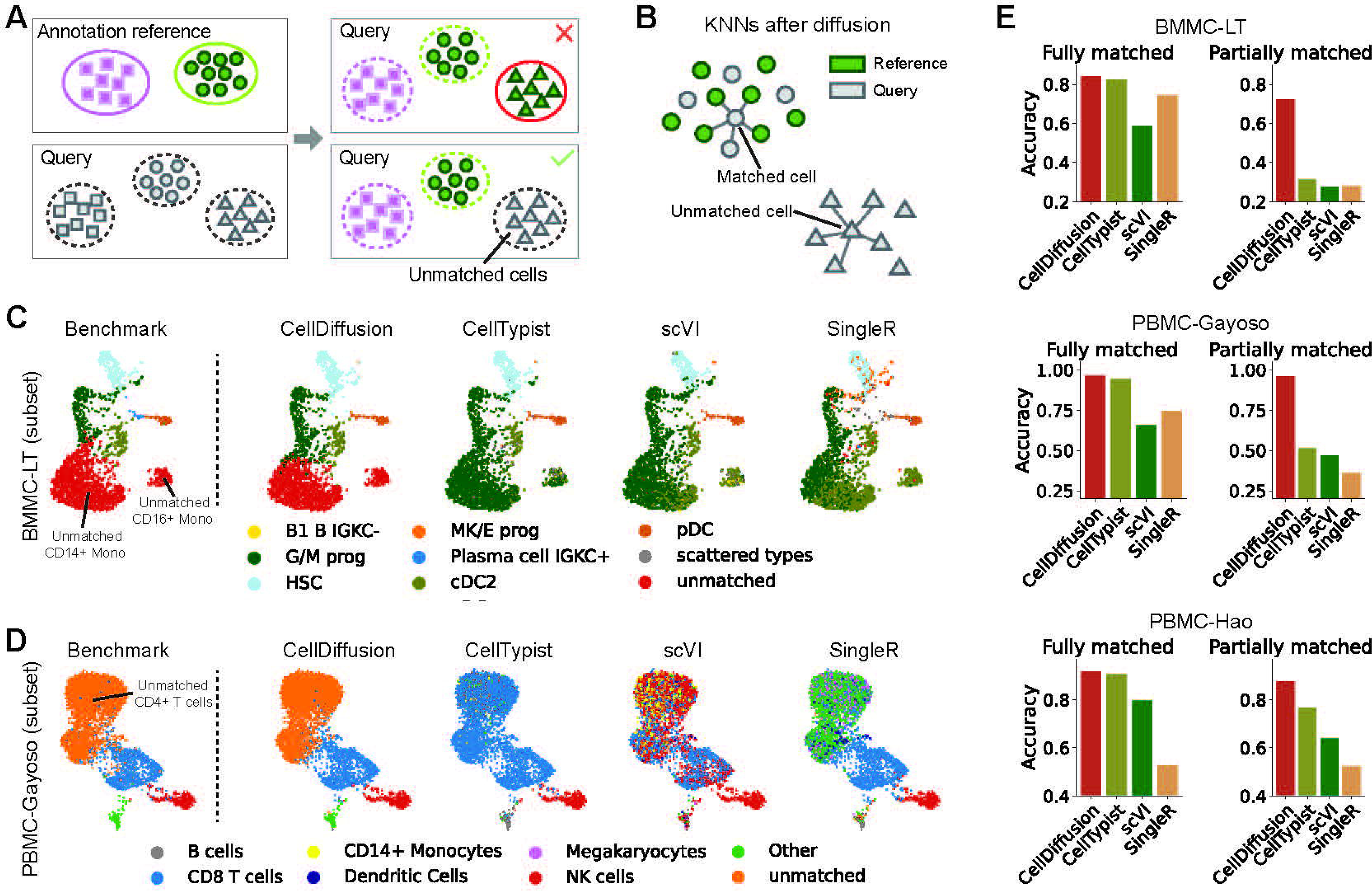
Benchmarking of reference-based cell type annotation methods. (**A**) Illustration of label transfer using an incomplete annotated reference. (**B**) Illustration of CellDiffusion distinguishing unmatched query data from the reference annotation. (**C**) UMAP visualizations highlighting a subset of the BMMC-LT dataset annotated using different cell type annotation methods. (**D**) UMAP visualizations highlighting a subset of the BMMC-Gayoso dataset annotated using different cell type annotation methods. (**E**) Quantitative evaluation of different cell type annotation methods applied to three datasets under two conditions (‘Fully matched’: the query samples have exact same cell types as the reference; ‘Partially matched’: the query samples contain unmatched cell types).

We thus benchmarked CellDiffusion against three widely used annotation methods: CellTypist(*37*), scVI(*12*), and SingleR(*3*). Their performance was tested in three gold-standard datasets with manually curated labels: BMMC-LT (29 cell types)(*35*), PBMC-Gayoso (9 cell types)(*36*), and PBMC-Hao (28 cell types)(*11*). To mimic real-world applications where the reference dataset does not always include all query cell types, several important query cell types were intentionally excluded from references. In this challenging setting, CellDiffusion consistently identified unmatched cell types, which were frequently misclassified by other methods (**Fig. 4C-D**; **figs. S13-S15**). For example, in the BMMC-LT dataset, CellDiffusion was the only method that successfully identified unmatched CD14+ and CD16+ monocytes, achieving at least 40% higher accuracy score over benchmark methods (**Fig. 4C, E**; **fig. S13**). Consistently, CellDiffusion accurately detected unmatched CD4+ T cells and FCGR3A+ monocytes in the PBMC-Gayoso dataset, with accuracy scores over 50% higher than the second-best model (**Fig. 4D-E**; **fig. S14**). CellDiffusion also demonstrated the best performance on the PBMC-Hao dataset (**fig. S15**). To ensure a comprehensive evaluation, we also tested the performance of label transfer when all query cell types are present in the reference dataset, where CellDiffusion still achieved the best accuracy (**Fig. 4E**). Notably, CellDiffusion’s annotation accuracy decreased only slightly by 1%-14% when transitioning from a fully matched to a partially matched reference, whereas competing methods exhibited dramatic drops of 16%-62%. CellDiffusion is less vulnerable to unmatched query cell types and offers a robust solution for annotating complex datasets where reference completeness is uncertain.

### CellDiffusion captures the structure of hierarchical cell ontology

CellDiffusion preserves the biological integrity during the imbalanced data integration and annotation through recovering KNN graph structures within each batch (**Fig. 1B and 2D**). In addition, intra-batch diffusion accurately captures the structure of hierarchical cell ontology. This is achieved through a combination of simultaneous local and global diffusion processes on single-cell graphs, inspired by physical principles of diffusion(*38*).

Within the KNN graph of a single batch, cells that are locally similar exhibit dense intra-group connectivity, enabling rapid local diffusion. This drives cells with a similar identity to cluster more tightly, which we refer to as the local equilibrium effect (**Fig. 5A**). The local equilibrium effect makes distant cell groups clearly separated from each other, enhancing the identification of independent cell populations. On the other hand, for cell types in the continuous biological trajectory, global diffusion propagates features across closely related populations, reinforcing inter-group relationships while unrelated cell types separate from each other. This process highlights lineage structures and reveals latent biological trajectories, termed the global equilibrium effect (**Fig. 5B**).

**Fig. 5.**
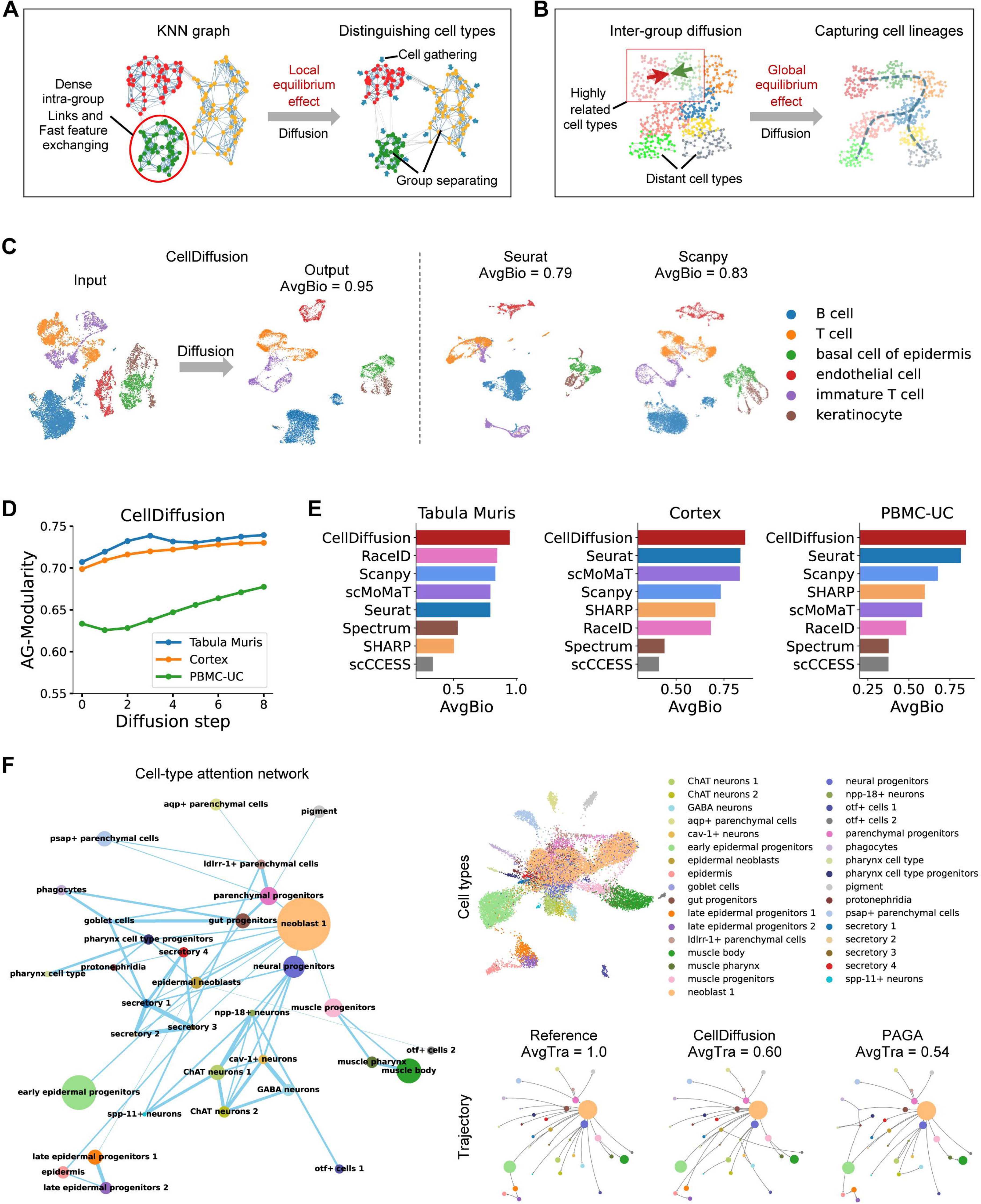
Benchmarking CellDiffusion and other methods for clustering and trajectory inference. (**A**) Illustration demonstrating how the local equilibrium effect enhances the separation of biologically distinct cell types. (**B**) Illustration demonstrating how the global equilibrium effect highlights biological lineages. (**C**) UMAP plots of the Tabula Muris dataset show that CellDiffusion embeddings have clearer boundaries of each cell type compared with Seurat and Scanpy. (**D**) The cell type-based AG-Modularity changes in the diffusion process for three datasets, including the Tabula Muris, Cortex, and PBMC-UC. (**E**) Quantitative evaluation of clustering results from eight methods across three datasets. (**F**) Attention-based trajectory inference applied to the ‘Planaria full’ dataset. The cell-type attention network reflects the differentiation relation between different cell types.

We demonstrated the local and global equilibrium effects in CellDiffusion on selected cells from a mouse dataset (Tabula Muris, six cell types, single batch)(*39*). Cell embeddings after diffusion illustrated condensed intra-cell type structures and clearer boundaries compared to the input data, demonstrating the local equilibrium effect (**Fig. 5C**). Meanwhile, the biologically related cell types, such as immature T cell and T cell, as well as the keratinocyte and basal cell of epidermis (precursors to keratinocyte), remain closely connected after diffusion, reflecting the global equilibrium effect. Together, these local and global equilibrium effects enable CellDiffusion to accurately preserve both discrete cell types and continuous lineages in the diffused embeddings compared with Seurat and Scanpy (**Fig. 5C**).

### CellDiffusion implements enhanced clustering and trajectory inference

Precisely recapitulating cell-cell associations through local and global diffusion processes can directly improve other inferences, including clustering and trajectory analysis. CellDiffusion includes built-in attention-based clustering and trajectory inference modules, enabling informative post-diffusion interpretation of cellular organization.

First, to quantitatively evaluate the contribution of local equilibrium effect to unsupervised clustering, we employed cell type-based attention modularity(*40*), which increases along with the diffusion process in the Tabula Muris dataset as well as two additional benchmark datasets, Cortex (7 cell types)(*2*) and PBMC-UC (8 cell types)(*11*) (**Fig. 5D**). The increasing modularity indicates improved separation of distinct cell types during diffusion, thereby enhancing the performance of unsupervised clustering. The clustering performance of CellDiffusion was further compared with seven existing methods: Seurat(*20*), Scanpy(*31*), scMoMaT(*41*), RaceID(*42*), SHARP(*43*), Spectrum(*44*), and scCCESS(*45*). CellDiffusion achieved the highest AvgBio scores across all three datasets (**Fig. 5E**) and consistently outperformed other methods in ARI, NMI, and FMI metrics (**table S1**), demonstrating its superior ability in unsupervised clustering.

Second, we applied the trajectory inference module of CellDiffusion to a scRNA-seq dataset of the regenerative flatworm *Schmidtea mediterranea* (Planaria full)(*46*), which encompasses 33 cell types arising from a continuous and well-orchestrated differentiation of stem cells (neoblast 1) into all major lineages, making it an ideal model to assess the performance of trajectory inference (**Fig. 5F**). CellDiffusion successfully extracted a high-quality tree-shaped trajectory that closely recapitulates inherent lineages (**Fig. 5F**). Thick attention edges in the cell-type attention network indicate strong similarity between linked cell types, reflecting robust relationships in the cell fate transition. To further benchmark its performance, we compared CellDiffusion against six existing trajectory inference methods, including PAGA(*21*), Slingshot(*47*), Slingshot Root(*47*), Monocle3(*48*), MST(*49*), and RaceID(*42*), in Planaria full and three additional datasets, Planaria pair (14 cell types)(*46*), Oligodendrocyte (12 cell types)(*50*), and Hepatoblast (11 cell types)(*51*). The AvgTra score represents the average of three graph-based evaluation metrics: graph edit index(*52*), graph kernel index(*53*), and Jaccard similarity index(*54*) (**Supplementary Text 4**). CellDiffusion consistently achieved a higher AvgTra score than competing methods in each dataset (**table S2**), indicating improved trajectory inference.

## Discussion

In this study, we introduce a graph-based diffusion framework designed to address data integration challenges at the co-occurrence of imbalanced cell compositions, unmatched cell types, and batch effects, which frequently arise in real-world datasets but cannot be handled properly by existing tools. Our model includes three practical innovations in data integration. First, it enables the automatic distinguishing of biological variations and batch effects, removing technical noise from cell embeddings. Second, as a dynamic model, it refines both short- and long-range cell-cell associations by employing local and global diffusion processes, preserving the hierarchical organization of cell ontology populations effectively during integration. Third, CellDiffusion employs a trainable attention mechanism(*25, 26*) to learn biological similarities and differences between cell populations across batches. It embeds learned cell-cell associations into a batch-removed cell-cell attention network, which guides and is jointly optimized with the diffusion process, enabling attention-based downstream analysis. Collectively, CellDiffusion constructs a robust cell-cell association graph that emphasizes biologically meaningful relationships while suppressing batch-induced noise.

Compared with studies involving decade-long efforts to improve batch effect correction, CellDiffusion better preserves biological variation without requiring manual input or assumptions about sample groupings. A recently published method, CellANOVA, significantly eliminates the overcorrection issue seen in Harmony. However, as illustrated in the endometrium dataset, CellANOVA risked defining possible biological differences as batch effects inside the control pool and retaining residual batch effects as biological signals between different sample conditions. Moreover, its reliance on control pools limits its applicability to single-condition datasets, such as the BMMC dataset. In contrast, CellDiffusion is a fully unsupervised method, offering broad generalizability across diverse experimental designs and dataset structures. We also compared CellDiffusion with UniPort(*17*) and BBKNN(*55*), where CellDiffusion consistently achieved better data integration performance (**Supplementary Text 5** and **figs. S16-17**).

Although the fuzzy clustering function from Harmony was used to construct the auxiliary graph in the OMNN algorithm, CellDiffusion does not use Harmony’s integration results or Harmony-corrected embeddings during the diffusion process. To evaluate the dependence of CellDiffusion on the auxiliary graph, we performed graph-construction ablation analyses using five strategies: pure MNN, Harmony-MNN, scVI-MNN, Harmony-assisted OMNN, and scVI-assisted OMNN. CellDiffusion maintained stable integration performance across all graph-construction strategies and consistently outperformed GCN models using the same input graphs (**fig. S18A**). Furthermore, in the mesenchymal lineage of the endometrial dataset, the Harmony- and scVI-assisted graphs showed lower biological conservation than the pure-MNN graph (**fig. S18B**), indicating that auxiliary batch-corrected representations can occasionally introduce biologically inappropriate edges. Together, these results demonstrate that Harmony is useful for improving candidate edge coverage, but it does not determine the final CellDiffusion integration results, which are driven by the subsequent diffusion process.

The effectiveness of CellDiffusion in data integration stems from its mathematical foundation in diffusion processes and its physical explanation of how to model both local and global diffusion across multiple scales of graph structures to detect biological signals, such as inherent cell types and lineages. From the perspective of algorithms, CellDiffusion generally advances common graph neural networks (GNNs)(*56–58*) by breaking their depth limitation, allowing the interaction field of each node to grow exponentially as network layers increase (**fig. S2E**). It overcomes the defects of GNNs like over-smoothing(*59, 60*), where deep layers obscure differences between cells, and over-squashing(*61*), where long-range dependencies are lost in densely connected regions (**Supplementary Text 6**). These advantages enable CellDiffusion to model heterogeneous, hierarchical intercellular relationships with high fidelity, avoiding major integration pitfalls such as downalignment and misalignment. The model collapse diagnostics on the Endometrium-CD dataset show that CellDiffusion maintains high precision when the number of network layers increases (from 2 to 16 layers) and outperforms the graph convolutional network (GCN), which exhibits distortion of data structures at deep layers (**fig. S19A**). We evaluated model performance across layers using neighbor purity, intrinsic dimension, and variance explained by embeddings metrics (**Supplementary Text 7**). CellDiffusion consistently achieved high and stable scores across different evaluation metrics and network layer depths, while GCN performance declined with layers increasing in neighbor purity and intrinsic dimension metrics and showed unstable performance in the evaluation using variance explained by embeddings (**fig. S19B**). This exactly proved that CellDiffusion avoids over-smoothing and breaks depth limitation of GCNs.

The diffusion process in CellDiffusion depends on two cell graphs, the OMNN graph which propagates cell features, and the batch-KNN graph, which constitutes the diffusion loss function. Hyperparameter sensitivity analysis on the OMNN and batch-KNN graph edge densities revealed that CellDiffusion performance remains robust across varying edge densities (**fig. S20**).

As an unsupervised deep learning model, CellDiffusion does not rely on pre-trained data and allows users to train the model from scratch with their own data. This gets rid of the harsh requirement for large amounts of training data, computing resources, and training time compared with pre-trained or fine-tuned foundation models(*62, 63*). Using a single L40S GPU, CellDiffusion can handle data integration tasks of tens of thousands of cells with eight diffusion layers or hundreds of thousands of cells in the high-throughput mode, where diffusion processes can be completed within 30 minutes. The integration runtime benchmark showed that CellDiffusion was clearly slower than Harmony, but comparable to Seurat and scVI (**table S3**).

It is important to acknowledge that, despite the high accuracy and fast speed of data integration achieved by CellDiffusion, challenges remain when processing datasets containing millions of cells on a single GPU, as the memory requirements can exceed available capacity. Scaling to large-scale datasets will require further development in batch training strategies and improvements in parallel computing to support efficient multi-GPU training. However, CellDiffusion’s memory usage is comparable to Seurat for integrating tens of thousands of cells (**table S4**). In addition, CellDiffusion operates under a widely adopted assumption: each batch shares at least a subset of common cell types, and batch-induced variation within the same cell type is smaller than transcriptional differences between distinct cell types within a batch(*7, 10*). As such, CellDiffusion is not designed to handle datasets with extremely strong batch effects that violate this fundamental assumption.

In the future, we will extend CellDiffusion to time-resolved scRNA-seq data(*64–67*). Diffusion is inherently a time-related process, making it well-suited for analyzing time-resolved datasets and exploring dynamic processes of biological changes. Additionally, applying CellDiffusion to spatial transcriptomics(*68–70*) presents an exciting opportunity. By constructing single-cell graphs with spatial coordinates, the diffusion model can direct information flow across spatial areas, enabling the capture of spatially specific characteristics to deepen our understanding of spatial transcriptomics. We envision the CellDiffusion framework being applied across various fields, advancing research in single-cell transcriptomics and beyond.

## Materials and Methods

### Diffusion equation on single-cell graphs

We construct the diffusion equation for CellDiffusion by following the mathematical framework of physical diffusion(*38*), with the modification of a discrete structure on graphs. Using the heat diffusion equation as an example, we demonstrate how to discretize it onto single-cell graphs. The diffusion equation for a heat diffusion process is given by:

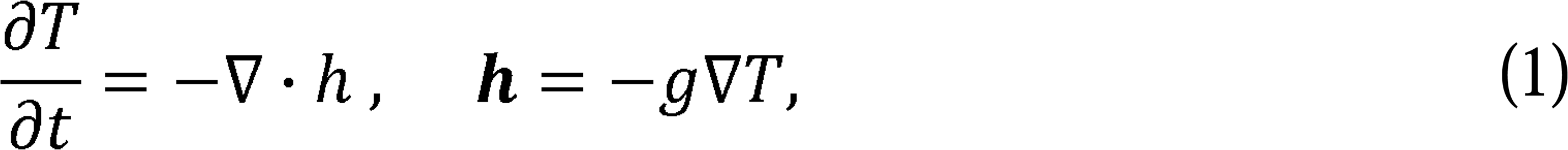

where the temperature *T* = *T*(***x***, *t*) is space coordinates ***x*** and time *t* dependent, and ***h*** is the heat flow in proportion to the gradient of temperature∇*T*. The coefficient *g* is called the thermal diffusivity(*71*), which is not only space and time but also temperature-dependent: *g* = *g*(***x***, *t, T*).

We project the diffusion equation (1) to discrete single-cell graphs by corresponding the temperature *T* with graph node features *X*. The gradient of the feature is defined on the edge between two linked nodes, which are denoted as node *α* and node *β*, in the following manner(*23*):

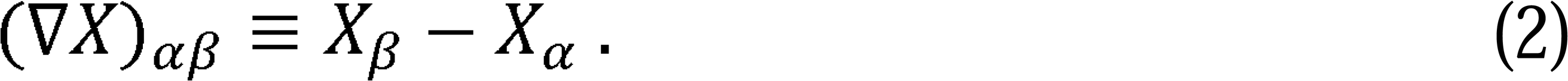

This definition is edge direction dependent, while (∇*X*)*_αβ_* is one type of edge features. For any edge feature *X*, we define its divergence as

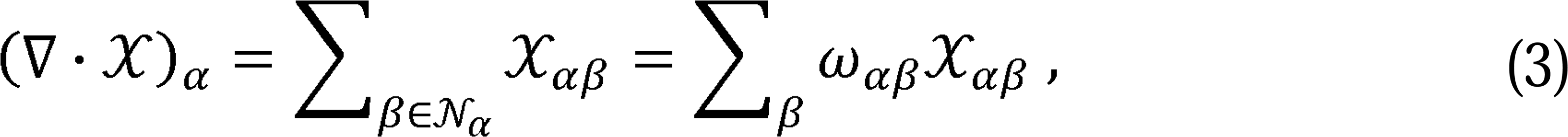

where *N_α_* represents the set of all linked neighbor nodes of the focal node *α*, and *ω* is the adjacency matrix of the single-cell graph. With the above definitions, the diffusion equation on the single graph reads

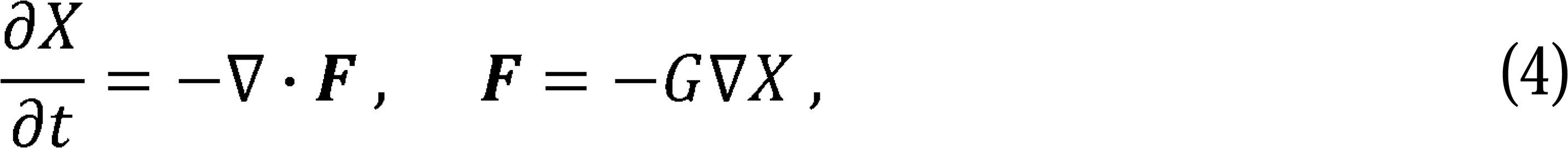

where ***F*** is the feature flow and *G* signifies the information diffusivity(*23*) on the cell graph. Since ***F*** and ∇*X* are located on graph edges, the information diffusivity *G* should be considered as another type of edge feature.

To simplify the calculations, we assume that the diffusivity on each edge is solely dependent on features of the two linked nodes, i.e., we have

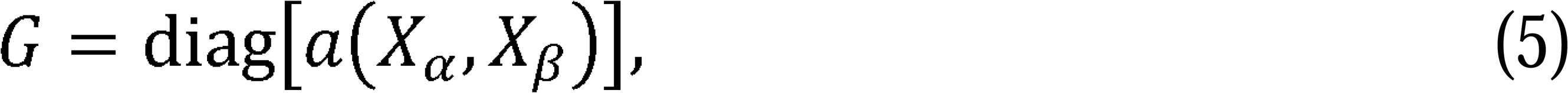

where *a*(·) is a scalar function. It is important to note that *G*, is an *e* ×*e* diagonal matrix, with *e* representing the total number of graph edges. It is useful to define a matrix *A* with the same structure as the adjacency matrix ω and the corresponding elements with *G*:

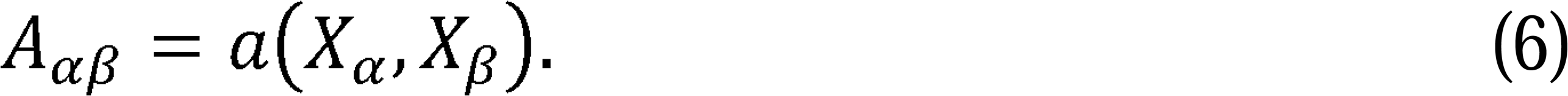

*A* is a *N* ×*N* matrix with its indices representing graph nodes, where *N* is the total number of graph nodes. We assume *A* to be right stochastic(*72*) for convenience, and then we can write Eq. (4) as

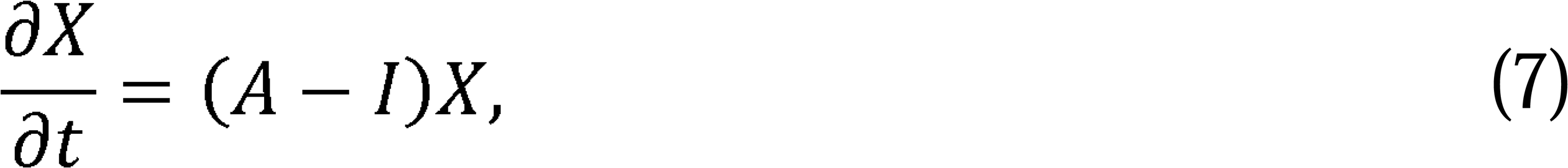

where *I* denotes the identity matrix.

To numerically solve the nonlinear diffusion equation (7), we further discretize it along the temporal direction, using the forward time difference:

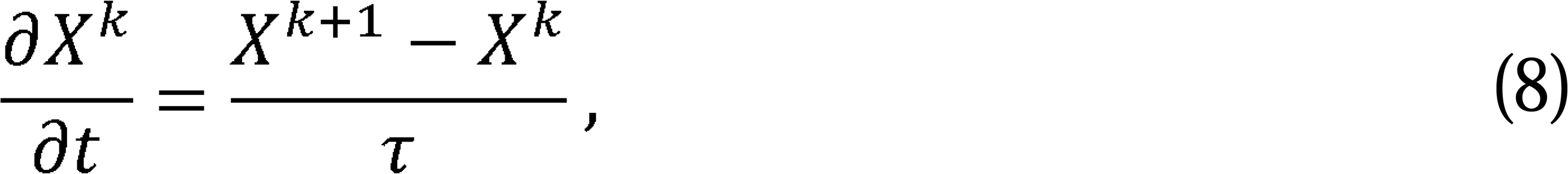

where *X^k^* and *X^k^*^+1^ denote the input and output features at *k* layer in the diffusion networks, and *τ* is the diffusion time interval in each layer. The total diffusion time is the total number of diffusion layers multiplied by *τ*. Now, we can write the diffusion equation (7) as follows:

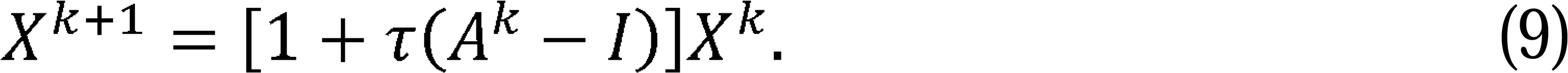

This scheme(*23*) is stable only when 0 < *τ <* 1. It is worth noting that the above equation shows the dimension of the node features *X* remains unchanged throughout the diffusion layers. Usually, the physical diffusivity(*71*) is carrier dependent. However, for computing convenience, we carry out 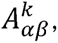 which represents the information diffusivity(*23*) on the edge between node *α* and node *β*, using the attention scheme(*25, 26*). The attention score between two linked nodes is learned based on node features on the single-cell graph. Thus, the diffusivity on the graph is indirectly dependent on the time *t* via node features.

### Attention scheme

In CellDiffusion, we use the following attention scheme to carry out the diffusivity(*23*) matrix *A* on a single-cell graph:

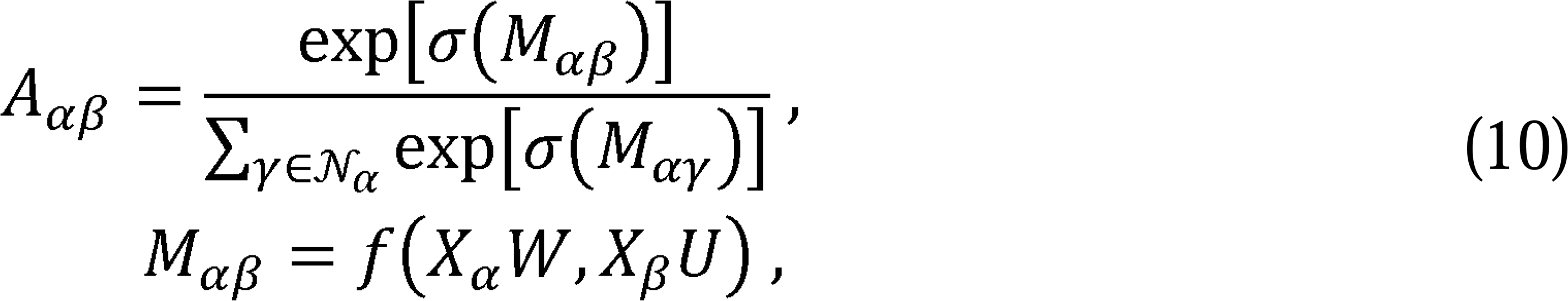

where *σ*(·) is the LeakyReLU function, *M_αβ_* is the attention coefficient, *f*(·) · is a scalar function, *W* and *U* are learned matrices. It is worth mentioning that the above attention matrix complies with all the necessary requirements for the diffusivity(*23*) on single-cell definitions of *f*(·) that correspond to the Bahdanau attention(*25, 57*) and scaled dot graphs. In CellDiffusion, we implement multi-head attentions and provide two optional product attention(*26*), respectively.

The Bahdanau attention(*25, 57*), also known as additive attention, assigns a different weight to each transformed feature and uses the sum of all weighted features for two linked nodes as the attention coefficient. The Bahdanau attention coefficient between node *α* and node *β* reads

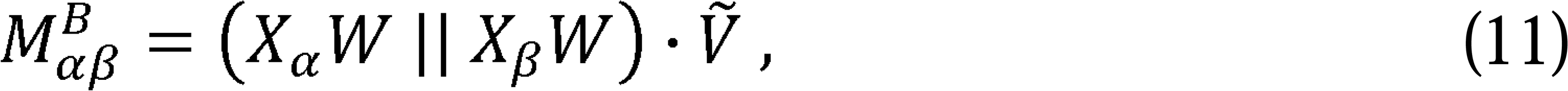

where the two linked nodes share the same weight matrix *W* and *Ṽ* is a learned vector. Without loss of generality, we remove the dependence of the weight matrix *W* in the above equation by defining a new weight vector *V*, and rewrite the equation as 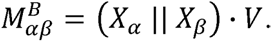 Unlike the graph attention network, where the weight matrix *W* is also during the diffusion process. Thus, the movement of the weight matrix *W* in Eq. (11) used in the dimension reduction process, CellDiffusion keeps the dimensionality invariant greatly reduces the total number of weights in the model, which accelerates the diffusion training process of CellDiffusion.

On the other hand, the scaled dot product attention(*26*) calculates the inner product of the weight matrix transformed features for two linked nodes. Following the work, we write the scaled dot product attention coefficient as

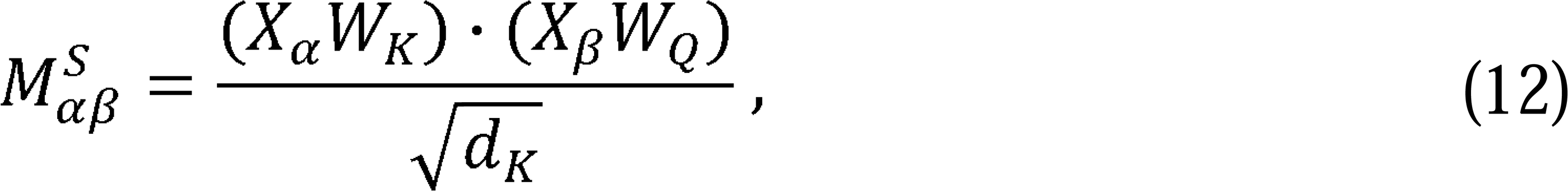

where the learned matrices *W_K_* and *W_Q_* are in the same shape, and *d_k_* is a hyperparameter determining the dimension of *W_K_*. We can also simplify the above equation by defining a new matrix 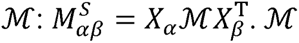 is a metric-like matrix that defines the inner product of two linked feature vectors *X_α_* and *X_β_*, which will be learned from the diffusion process.

In CellDiffusion, we share the same learned weights, i.e., the same weight vector *V* or matrix *M*, through all diffusion layers. The scaled dot product attention outperforms the Bahdanau attention in the language model(*26*). However, the number of learned weights for the scaled dot product attention increases fast as the number of feature dimensions increases in CellDiffusion. We verify that the Bahdanau attention performs much the same as the scaled dot product attention in scRNA-seq data analysis. To improve computational efficiency, our calculations in this work are done using the Bahdanau attention scheme. We reserve the option for users to use the scaled dot product attention scheme in CellDiffusion.

### From graph topology to cell-cell attention network

The graph topology is widely used to study intercellular relationships in various methods for scRNA-seq analysis, such as UMAP embeddings(*73*), graph-based clustering(*74, 75*), and trajectory inference(*21*). In CellDiffusion, we train a cell-cell attention network to present more accurate intercellular dependencies (**fig. S2B**). Attention plays a role of the information diffusivity on the graph. A high (or low) attention score indicates a strong (or weak) information flow along an edge (**fig. S2C**). Cells of the same inherent cell type are tightly connected and tend to become more similar with each other during diffusion, resulting in strong attention and high diffusion rates between these cells. Conversely, cells from different cell types are more distantly connected, leading to weak attention and slower diffusion between them. The cell-cell attention network directly improves the accuracy of graph-based cell-type clustering methods. By utilizing the cell type labels, we transform the cell-cell attention network into a cell-type attention network and use it to infer pseudo-time transitions(*4*).

### Building a mutual nearest neighbor graph for data integration

We use diffusion to balance data distribution across different batches and eliminate batch effects. A critical step in this process is constructing a single-cell graph that connects cells from different batches. We employ the mutual nearest neighbor (MNN) algorithm(*7*) to build a graph, *G*_1_, which typically connects cells of the same inherent cell types rather than cells from different types across batches. However, graph *G*_1_ can be flawed due to highly uneven connections caused by batch effects (**fig. S21A**). Cells in one batch tend to identify the same K-nearest neighbors (KNNs) in another batch due to proximity, which leads to only a few cells from the two batches being linked in the MNN graph, diminishing its effectiveness in removing batch effects.

To address the issue of uneven connections, we optimized the MNN method and built the OMNN algorithm. We perform a fuzzy clustering-based alignment of embeddings using Harmony(*10*) to build the second MNN graph *G*_2_ (**fig. S21B**). The fuzzy clustering-based alignment mixes cells across batches, making *G*_2_ a more evenly connected graph. After that we project *G*_2_ back onto the original embeddings, because the diffusion process for data integration still uses the original embeddings as input. We then combine the graph *G*_1_ and *G*_2_, taking the union of their edge sets to construct the OMNN graph:

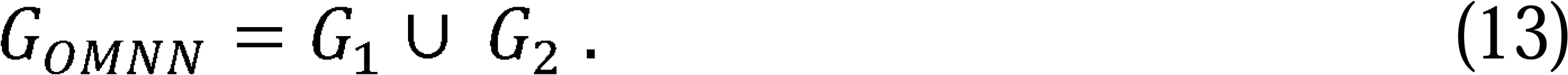

While *G_OMNN_* connects many cross-batch cell pairs, it may not include all cells, especially those that lack counterparts in other batches. To ensure complete connectivity, we build the supplementary graph *G_KNN_* by connecting unlinked and sparsely-linked cells to their nearest neighbors in the entire dataset to make sure that each cell has the same number of linked neighbors. The final diffusion graph for data integration reads:

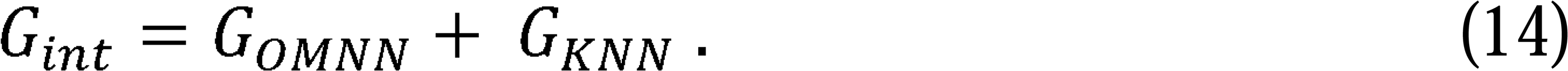

This final graph ensures that every cell is included in the diffusion process.

We emphasize that the OMNN graph in CellDiffusion is used exclusively for data integration but not for other tasks such as clustering or trajectory inference, as it is designed to deal with inter-batch cell correspondences. Due to the lack of intra-batch connections, the OMNN graph is not suitable for these tasks. For clustering and trajectory analysis, we build a standard KNN graph using the integrated embeddings to connect all cells without considering batches. We project the attention weights onto the KNN graph and obtain the final clustering or trajectory results.

### Label transfer algorithm

CellDiffusion transfers labels from the reference dataset to the aligned cells in the query dataset after integrating both datasets. Before label transfer, we identify unaligned cells in the query dataset with respect to the reference, following the data integration, to detect potential cell types in the query dataset that are not represented in the reference. To evaluate the alignment between query and reference cells, we construct a KNN graph on the integrated dataset, with a default K=100. Each query cell is flagged as poorly aligned if fewer than 5% of its K-nearest neighbors are reference cells. Further, if more than 80% of the neighbors of a poorly aligned cell are themselves poorly aligned, the cell is labeled as unaligned. We exclude these unaligned cells from label transfer. For the remaining query cells, we identify their KNNs in the reference dataset and transfer the label that appears most frequently among their neighbors.

### Graph modularity and unsupervised clustering

Graph modularity(*40*) is a measure used in network theory to quantify the strength of the division of a graph into clusters or communities. It evaluates how well the graph can be partitioned into distinct groups where nodes are densely connected within the same community but sparsely connected across different communities. The modularity value ranges from 0 to 1, where the value 1 represents all communities that are well separated from each other, and the value 0 indicates that communities are extremely mixed with each other.

Mathematically, modularity compares the actual density of edges within communities to the expected density of edges if they were randomly distributed. In this work, we used the attention-weighted edges to compute the graph modularity of the cell-cell attention network, with respect to cell types or data batches. The attention-based graph modularity (AG-Modularity) is calculated as follows(*76*):

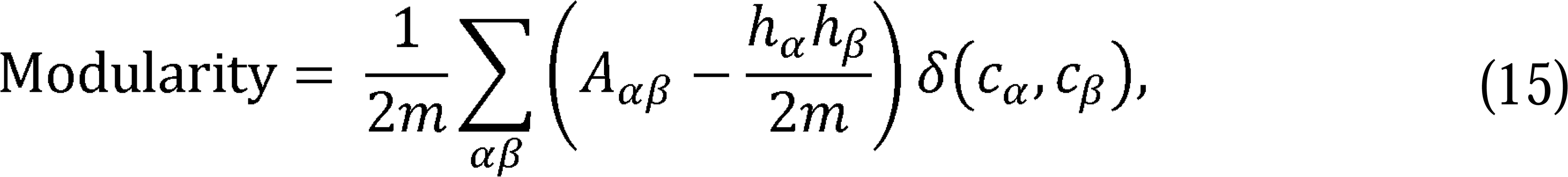

where *m* is the sum of all edge weights, *A* is the attention matrix, *h_α_* is the weighted degree of node *α*, and *δ*(*C_α_*, *C_β_*) is 1 if node *α* and node *β* are in the same community else 0.

Graph modularity helps in uncovering the underlying structure of complex networks. A higher cell-type-based modularity occurs with a clearer separation of different cell types, which usually indicates more accurate unsupervised clustering results to identify those cell types in the dataset. On the other hand, a lower batch-based modularity indicates that data from different batches are better aligned.

CellDiffusion contains an unsupervised graph-based clustering method to identify cell types in scRNA-seq data. We use Leiden algorithm to compute graph modularity and optimize cell clusters based on the cell-cell attention network (KNN graph), with attention scores serving as edge weights. The optimized clustering result is derived by maximizing the modularity for cell clusters.

### Trajectory algorithm

CellDiffusion builds the directed tree-shaped trajectory based on cell types using the cell-type attention network. We construct the cell-type attention network by aggregating cell-cell attention between different cell types. The attention between two different cell types *u* and *v* is defined as:

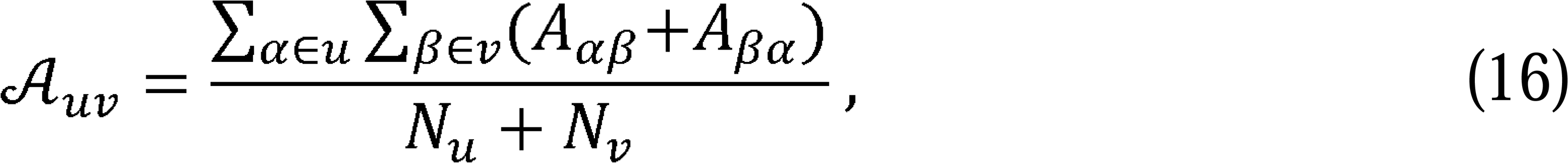

where *N_u_* is the total number of cells with cell type *u* and *N_v_* is the total number of cells with cell type *v*. The cell-type attention matrix *A* is a symmetric hollow matrix with zero diagonal elements, i.e., *A_uu_* = 0.

We normalize *A* by setting the average of its non-zero elements to 1. The normalized cell-type attention matrix reads:

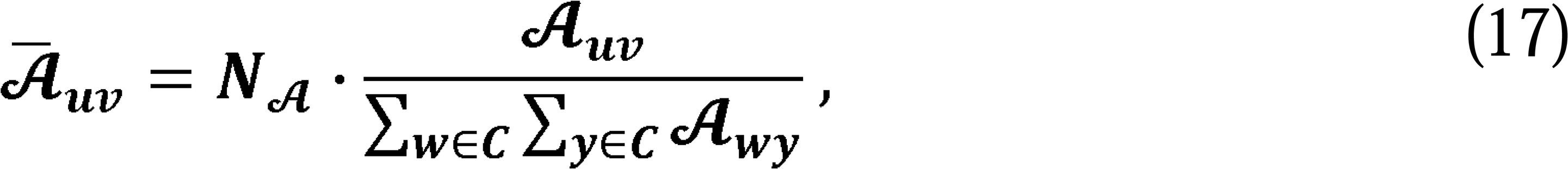

where *N_A_* is the number of non-zero elements of *A*, *C* is the set of all cell type labels. We define the distance between cell types *u* and *v* as the inverse of the log-changed normalized attention plus 1:

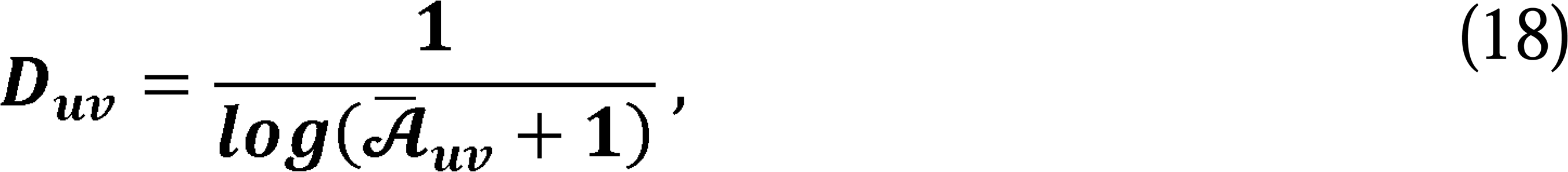

where two cell types with zero attention have the infinite distance between them. The greater the attention between two cell types, the smaller the distance between them. We calculate the length of a given path on the fully connected cell-type graph, by aggregating the edge distances on the path,

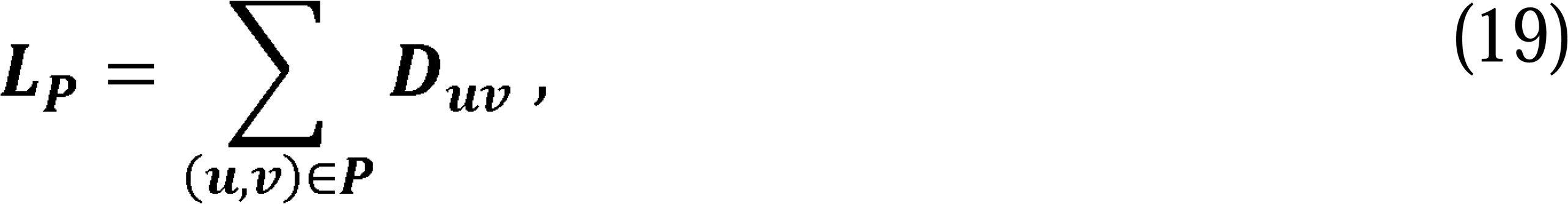

where *L_P_* represent the length of path *P* and (*u,v*) denotes the undirected edges. The tree-shaped trajectory is constructed by using the Shortest Path Tree approach, finding the shortest path from the root cell type to every other cell type.

### CellDiffusion framework

The CellDiffusion framework includes three main components: the autoencoder for dimension reduction, diffusion layers for conducting information flow on the single cell graph, and the decoder to reconstruct either the high-dimensional input data or the original adjacency matrix (**fig. S1A**). CellDiffusion takes normalized gene expression data as input. Preprocessing involves standardizing raw data from N cells by normalizing the total reads per cell to 10,000. By default, the top 2,000 variable features are selected as input, encoded into the latent space, and then diffused.

In data integration tasks, CellDiffusion accepts multi-batch datasets as input and generates the KNN adjacency matrix for each batch individually before diffusion. These batch-specific adjacency matrices are recovered in the decoding process after diffusion (**fig. S1B**). The output of the diffusion process for data integration includes batch-effect-corrected embeddings and the cell-cell attention network, which can be directly used for downstream tasks like annotation, clustering and trajectory inference.

### Data integration benchmarking

Using endometrium datasets, we compare the performance of CellDiffusion against Harmony (using Harmonypy 0.0.9, Python)(*10*) and CellANOVA (version 0.1.0, Python)(*19*), where CellANOVA was applied using secretory-stage samples as the control pool. Due to its reliance on manually defined control pools, CellANOVA could not be applied to other benchmark datasets in this study and was therefore excluded from other benchmarking tasks.

For benchmarking on additional integration datasets, we compare CellDiffusion with six existing data integration methods, Harmony(*10*), scVI (using scvi-tools 1.1.2, Python)(*12*), fastMNN (version 1.26.2, R)(*7*), Seurat (version 4.1.1, R)(*11*), ComBat (using Scanpy 1.9.8, Python)(*9*), and Scanorama(version 1.7.4, Python)(*8*). We use three evaluation metrics in the data integration benchmarking: BioCon, BatchCor, and Total(*27*). The BioCon score evaluates biological conservation and aggregates five cell type-based metrics, including isolated labels(*27*), Kmeans NMI(*33*), Kmeans ARI(*32*), silhouette label(*77*), and cLISI (cell-type local inverse Simpson’s index)(*10*). The BatchCor score assesses batch correction and aggregates four metrics, including silhouette batch(*77*), iLISI(*10*), KBET(*78*), and graph connectivity(*27*). The Total score represents the overall performance in both biological conservation and batch correction. More details on these evaluation metrics can be found in **Supplementary Text 1**.

### Reference-based annotation benchmarking

We benchmark CellDiffusion with three existing reference-based annotation packages, including CellTypist (version 1.6.3, Python)(*37*), scVI (using scvi-tools 1.1.6, Python)(*12*), and SingleR (version 2.6.0, R)(*3*). Annotation performance is evaluated in two cases, one where all cell types in the query dataset are included in the reference dataset (fully aligned case), and another where there are some cell types in the query dataset are not included in the reference dataset (partially aligned case). We compare the annotated labels with the benchmark labels for each dataset and calculate the percentage of correct annotations across all cells.

### Unsupervised clustering benchmarking

We benchmark CellDiffusion with seven existing clustering methods, Seurat (version 5.0.3, R)(*20*), Scanpy (version 1.10.1, Python)(*31*), scMoMaT (version 0.2.0, Python)(*41*), RaceID (version 0.3.5, R)(*42*), Spectrum (version 1.1, R)(*44*), SHARP (version 1.1.0, R)(*43*), and scCCESS (version 0.3.3, R)(*45*). To ensure a fair comparison, we fine-tune the clustering resolution parameter in each package to match the number of clusters with the number of cell types in the benchmark dataset. We use three clustering metrics to evaluate the performance of these packages: the adjusted Rand index (ARI)(*32*), normalized mutual information (NMI)(*33*), and the Fowlkes-Mallows index (FMI)(*34*) (**Supplementary Text 2**), along with an aggregated score, AvgBio, which represents the average of these three metrics.

### Trajectory inference benchmarking

We benchmark CellDiffusion against six existing trajectory inference methods: PAGA (using Scanpy 1.9.8, Python)(*21*), Slingshot/Slingshot Root (version 2.12.0, R)(*47*), Monocle3 (version 1.3.7, R)(*48*), MST (using dynverse from https://dynverse.org, R)(*49*), and RaceID (version 0.3.5, R)(*42*). CellDiffusion, PAGA, and Monocle3 require the root cell type of the trajectory as prior input, while in Slingshot, the root cell type is optional. Therefore, we refer to the version of Slingshot with the root cell type specified as ‘Slingshot Root’. MST and RaceID do not require a specified root cell type and output undirected trajectories, while other methods output the directed trajectories.

Trajectory formats vary across packages. To make the results comparable, we convert all outputs into directed tree-shaped trajectories with the same format as the benchmark trajectory. For MST and RaceID, the trajectory direction is achieved by applying the root cell type to the predicted trajectory. Monocle3 and MST are excluded from the benchmark on the Planaria full dataset because these two packages do not directly generate trajectories based on existing cell types, and their output is difficult to interpret for 33 cell types in the Planaria full dataset.

We introduce three graph-based metrics to evaluate the similarity between the predicted and benchmark trajectories, including graph edit index(*52*), graph kernel index(*53*), and Jaccard similarity index(*54*) (**Supplementary Text 4**). The aggregated score, AvgTra, represents the average of three evaluation metrics.

### Benchmarking datasets

#### Endometrium-314k

The Endometrium-314k dataset was collected from a flagship single-cell reference atlas of the human endometrium(*28*). It includes samples from 87 women with and without endometriosis, and covers all menstrual cycles including menstrual, proliferative, and secretory phases. This dataset contains 313,527 cells, 17,736 genes and 36 cell types.

#### Endometrium-35k

The Endometrium-35k dataset is a subset of the Endometrium-314k dataset(*28*). We randomly selected 4 samples, 2 from the proliferative phase and 2 from the secretory phase, corresponding to 4 batches (GSM4577308, GSM6605431, GSM6605434 and GSM6605439). In Endometrium-35k dataset, we removed those cell types with less than 50 cells, resulting in a final dataset of 34,658 cells and 23 cell types.

#### Endometrium-CD

The Endometrium-CD dataset is a subset of the Endometrium-314k dataset. It contains 2 samples, one from the proliferative phase and another from the secretory phase, corresponding to 2 batches (GSM6605435 and GSM6605434).

#### BMMC

The BMMC dataset is a subset of bone marrow mononuclear cells sequenced using the CITE-seq protocol(*35*). The original dataset came from 12 healthy human donors. We selected the data from 4 donors, comprising 30,669 cells, and representing 4 batches (s1d1, s2d1, s3d1, and s4d1). The BMMC dataset contains 14,087 genes and 40 detailed immune cell subtypes. We manually introduce cell type imbalances by removing CD4+ T activated cells from batches s1d1 and s2d1 and Erythroblast cells from batches s3d1 and s4d1 at different percentages (0%, 25%, 50%, 75%, and 100%), generating five cell type-imbalanced BMMC datasets with varying degrees of imbalance: BMMC (imbalance = 0%, 25%, 50%, 75%, and 100%).

#### PBMC-23k

The PBMC-23k dataset is a subset of the whole peripheral blood mononuclear cells sequenced using the CITE-seq(*11*). The whole PBMC dataset contains samples from 8 patients enrolled in an HIV vaccine trial measured at 3 time points, and was reprocessed by Gayoso et al.(*36*), resulting in 152,094 cells after quality control. The PBMC-23k dataset includes the data from one patient across 3 time points, which indicates 3 batches (P8_0, P8_3, and P8_7). It contains 23,589 cells and 20,729 genes. The annotations in the PBMC-23k dataset include 30 cell types. The cell type imbalance is introduced by manually removing CD4 Naive cells from batch P8_0, and NK cells from batch P8_3 at different percentages (0%, 25%, 50%, 75% and 100%), generating five cell type-imbalanced PBMC-23k datasets with varying degrees of imbalance: PBMC-23k (imbalance = 0%, 25%, 50%, 75%, and 100%).

#### PBMC-10k

The PBMC-10k dataset includes two scRNA-seq batches (batch_0 and batch_1) of human PBMCs from one healthy donor. The dataset was reprocessed by Gayoso et al.(*36*). Batch_0 contains 7,982 cells, while batch_1 contains 4,008 cells. The dataset contains 3,346 genes and nine cell types annotated using Seurat(*20*), namely, B cells, CD4+ T cells, CD8+ T cells, CD14+ Monocytes, Dendritic cells, NK cells, FCGR3A+ Monocytes, Megakaryocytes and Other. The cell type imbalance is introduced by manually removing CD4 T cells from batch_0 at different percentages (0%, 25%, 50%, 75%, and 100%), generating five cell type-imbalanced PBMC-10k datasets with varying degrees of imbalance: PBMC-10k (imbalance = 0%, 25%, 50%, 75%, and 100%).

#### PBMC-Gayoso, PBMC-Gayoso-ref1, PBMC-Gayoso-ref2

The PBMC-Gayoso and PBMC-Gayoso-ref1 datasets correspond to batch_0 and batch_1 of the PBMC-10k dataset(*36*), respectively, and both contain the nine cell types present in the PBMC-10k dataset. The PBMC-Gayoso-ref2 dataset is derived from PBMC-Gayoso-ref1 by removing CD4 T cells and FCGR3A+ Monocytes resulting in a reduced reference of 3,346 cells that do not fully cover all cell types. PBMC-Gayoso-ref1 and PBMC-Gayoso-ref2 are used as reference datasets to perform label transfer on the PBMC-Gayoso dataset individually, representing fully and partially aligned scenarios, respectively.

#### PBMC-Hao, PBMC-Hao-ref1, PBMC-Hao-ref2

These three datasets are subsets of the PBMC-23k dataset(*11*). The PBMC-Hao dataset is from one donor (P7_0), containing 8,843 cells, 20,729 genes, and 28 cell types. The PBMC-Hao-ref1 dataset is from another donor (P8_3), containing 8,289 cells, 20,729 genes, and 30 cell types, covering all cell types represented in the PBMC-Hao dataset. The PBMC-Hao-ref2 dataset is derived from PBMC-Hao-ref1 by removing B memory, B naïve, CD4 CTL, CD16 Mono, and cDC2 cells resulting in a reduced reference of 7,570 cells that do not fully cover all cell types. PBMC-Hao-ref1 and PBMC-Hao-ref2 are used as reference datasets to perform label transfer on the PBMC-Hao dataset individually, representing fully and partially aligned scenarios, respectively.

#### BMMC-LT, BMMC-LT-ref1, BMMC-LT-ref2

These three datasets are subsets of the BMMC dataset(*35*). The BMMC-LT dataset contains 5,574 cells from one donor (s2d4) with 29 cell types and 14,087 genes. The BMMC-LT-ref1 dataset contains 4978 cells from another donor (s1d2) with 41 cell types, covering all cell types represented in the BMMC-LT dataset. The BMMC-LT-ref2 dataset is derived from BMMC-LT-ref1 by removing CD4+ T naïve, CD14+ Mono, CD16+ Mono, Erythroblast, Naïve CD20+ B IGKC+, and Reticulocytes, resulting in a reduced reference of 3,320 cells that do not fully cover all cell types. BMMC-LT-ref1 and BMMC-LT-ref2 are used as reference datasets to perform label transfer on the BMMC-LT dataset individually, representing fully and partially matched scenarios, respectively.

#### Tabula Muris

The Tabula Muris dataset is a subset from the Tabula Muris Senis(*39*) dataset. We selected 11,470 cells from one female mouse to build the Tabula Muris dataset. This dataset contains 6 cell types from variant tissues and 23,433 genes in each cell.

#### Cortex

The Cortex dataset contains scRNA-seq data from two regions of the mouse cerebral cortex: the somatosensory cortex and hippocampus CA1(*2*). The dataset was reprocessed by Gayoso et al.(*36*) and can be accessed from scVI tools using the API scvi.data.cortex. It includes 3,005 cells, 19,972 genes, and 7 cell types (astrocytes_ependymal, endothelial-mural, interneurons, microglia, oligodendrocytes, pyramidal CA1, and pyramidal SS).

#### PBMC-UC

The PBMC-UC dataset is a subset of the PBMC-23k dataset(*11, 36*), containing the data from a single donor at one time point. It includes 5,686 cells and 20,729 genes. The annotations in the PBMC-UC dataset include 8 cell types (B, CD4 T, CD8 T, DC, Mono, NK, other T, and other).

#### Planaria full, Planaria pair

The Planaria full and pair datasets are scRNA-seq from adult Schmidtea mediterranea(*46*), and contain a large pool of pluripotent stem cells and continuously differentiating cell types. The data was reprocessed by Saelens et al.(*49*). The Planaria full dataset contains 18,837 cells, 4,210 genes and 33 cell types, while the Planaria pair dataset contains 14,636 cells, 4,210 genes and 14 cell types. Each dataset includes a clearly defined lineage tree.

#### Oligodendrocyte

The Oligodendrocyte dataset contains scRNA-seq data of oligodendrocyte cell lineage from ten regions of the mouse juvenile/adult central nervous system(*50*). The data was reprocessed by Saelens et al.(*49*). It includes 4,930 cells, 3,534 genes and 12 cell types (OPC, COP, NFOL1, NFOL2, MFOL1, MFOL2, MOL1, MOL2, MOL3, MOL4, MOL5, and MOL6).

#### Hepatoblast

The Hepatoblast dataset(*51*) contains scRNA-seq data of mouse embryo cell lineage from the bi-potential hepatoblasts to hepatocytes and cholangiocytes, collected between embryonic day 10.5 to 17.5. The data was reprocessed by Saelens et al.(*49*), and includes 504 cells, 6,138 genes and 11 cell groups.

## Acknowledgments

We thank Bingtian Ye and Saumya Pothukuchi at Boston University and Nidhi Siddharam Loni at Indiana University for their help in running some benchmark packages.

## Funding

National Key Research and Development Project of China 2023YFC2705901 (HP)

Key Clinical Projects of Peking University Third Hospital BYSYZD2023028 (HP)

Beijing Nova Program 20220484073 (HP).

AnalytiXIN initiative (JW)

National Institutes of Health grants R01HL159620 (VBK)

National Institutes of Health grants R01AG083735 (VBK)

National Institutes of Health grants R01AG062109 (VBK)

National Institute on Aging’s Artificial Intelligence and Technology Collaboratories P30AG073104 (VBK)

National Institute on Aging’s Artificial Intelligence and Technology Collaboratories P30AG073105 (VBK)

Wing Tat Lee award (CZ)

American Cancer Society TheoryLab Collaborative Grant (CZ)

American Cancer Society Institutional Research Grant Pilot Award (CZ)

## Author contributions

Conceptualization: CZ, HP, JW

Methodology: YCL, CZ, JW, DX

Investigation: ZH, CZ, HP

Data curation: YCL, ZH, ZL, RL

Formal analysis: ZH, YCL

Software: YCL, LJ

Validation: YCL, ZH, SLL, LJ, JHL

Visualization: ZH, YCL, CZ

Supervision: CZ, HP, JW, DX, RL

Writing-original draft: YCL, CZ, HP, ZH, JW

Writing-review & editing: CZ, ZH, YCL, HP, JW, DX, RL, VBK

## Data, Code, and Materials Availability

All data and code needed to evaluate and reproduce the results in the paper are present in the paper and the Supplementary Materials. This study did not generate new materials. The Endometrium dataset can be downloaded through the web portal: https://www.reproductivecellatlas.org/endometrium_reference.html. The BMMC dataset can be accessed from the GEO database via accession number GSE194122, and the PBMC-10k dataset can be accessed using API scvi.data.pbmc_dataset. The PBMC-23k datasets can be extracted from the scVI tools dataset with the API scvi.data.pbmc_seurat_v4_cite_seq. The Tabula Muris dataset is accessible from a public resource (http://tabula-muris.ds.czbiohub.org/). The Cortex can be accessed from scVI tools using API scvi.data.cortex, The Planaria full, Planaria pair, Oligodendrocyte and Hepatoblast datasets can be accessed from Zenodo (https://doi.org/10.5281/zenodo.1443566). The source code of CellDiffusion is freely available at https://github.com/CZCBLab/CellDiffusion.

