## Supplementary material for "CellDiffusion: Enhancing scRNA-seq data integration with accurate inference of cell-cell association"

Science Advances
AAAS

### Supplementary Materials for

#### CellDiffusion: Enhancing scRNA-seq data integration with accurate inference of cell-cell association

Yu-Chen Liu *et al.*

**This PDF file includes:**

Supplementary Text
Figs. S1 to S22
Tables S1 to S4
References (*80*)

#### Supplementary Text

##### 1. Evaluation metrics in data integration benchmarking

We used the evaluation metrics outlined by Luecken et al.(*27*) in their benchmark study. Each metric is described below.

***Graph LISI***

LISI (local inverse Simpson's index)(*10*) is a metric designed to assess the degree of mixing or separation in high-dimensional data, specifically proposed for evaluating batch mixing (iLISI) and cell-type separation (cLISI) in the context of batch effect correction or integration in scRNA-seq data analysis. LISI scores are computed from neighborhood lists for each node in integrated KNN graphs. Specifically, the inverse Simpson's index determines the number of cells that can be drawn from a neighbor list before a batch is observed twice. As a result, LISI scores range from 1 to $B$, where $B$ is the total number of batches in the dataset. Varying the number of nearest neighbors per node can lead to different sensitivities within neighborhoods, skewing comparisons with graph-based integration outputs. Therefore, the original LISI score is not directly applicable to graph-based outputs.

Graph LISI extends the original LISI to be applicable to graphs by using the integrated graph structure as an embedded space for distance calculation. The computed graph distances are then used to determine a consistent number of nearest neighbors per node. Since LISI scores range from 1 to $B$, indicating perfect separation at one end and perfect mixing at the other, they are rescaled to a 0 to 1 range. For cLISI, a value of 0 corresponds to low cell-type separation, while for iLISI, a value of 0 indicates low batch integration.

***Average silhouette width***

The silhouette width(*78*) measures how similar a cell is to its own cluster compared to other clusters, evaluating the relationship between the within-cluster distance of a cell and its distance to the nearest neighboring cluster. By averaging the silhouette widths of all cells, we obtain the average silhouette width (ASW), which ranges from -1 to 1. The ASW is commonly used to assess the separation of clusters, where a value of 1 indicates dense and well-separated clusters, while values close to 0 or -1 suggest overlapping clusters or significant misclassification, respectively.

To evaluate data integration outputs, we applied the classical definition of ASW to determine the silhouette for cell type labels (silhouette label), scaling it to a value between 0 and 1 using the following formula:

$$\textbf{Silhouette label} = \frac{\boldsymbol{ASW_{label}} + \mathbf{1}}{\mathbf{2}}.$$

For assessing batch effects, we used a modified approach, considering the absolute silhouette width, s(i), for batch labels of each cell i. The silhouette batch is defined as:

$$\textbf{Silhouette batch} = \frac{\mathbf{1}}{|\boldsymbol{L_B}|}\sum_{j\in L_B}\frac{\mathbf{1}}{|\boldsymbol{C_j}|}\sum_{i\in C_j}(\mathbf{1} - |\boldsymbol{s(i)}|),$$

where $L_B$ is the set of unique batch labels, $|L_B|$ is the number of elements in the set, and $C_j$ is the set of cells with the batch label j. A value of 1 represents ideal batch mixing and a value of 0 indicates strongly separated batches.

***Isolated labels***

The isolated labels metric(*27*) evaluates how well these isolated cell type labels separate from other cell identities, where the isolated cell type labels are identified as the labels present in the least number of batches in the integration task. This is achieved in two ways, the best clustering of the isolated label and the global ASW of the isolated label. In the clustering method, we optimize the cluster assignment of the isolated label across Louvain clustering resolutions ranging from 0.1 to 2 in resolution steps of 0.1. The best cluster assignment corresponds to the highest $I_1$ score defined using the precision and recall given by the equation:

$$\boldsymbol{I_1 = 2 \times \frac{\text{precison} \times \text{recall}}{\text{precison} + \text{recall}}}.$$

It returns a value between 0 and 1, where the value 1 shows that all the isolated label cells and no others are captured in the cluster. For the isolated label ASW score, we compute the ASW of isolated versus non-isolated labels and scale this score to be between 0 and 1. The final score of Isolated labels are the average of the above two scores.

***KBET***

KBET (K-nearest neighbor batch effect test)(*79*) is a statistical method used to assess the effectiveness of batch effect correction, evaluating whether cells from different batches are evenly mixed in the corrected data. Specifically, KBET operates on KNN graphs, which are computed based on the corrected feature outputs. It determines whether the label composition of a K-nearest neighborhood of a cell is similar to the expected label composition. If the cells within a neighborhood are mostly from the same batch, this indicates poor mixing, suggesting that the batch effect correction was inadequate. The test is repeated for a random subset of cells, and the results are summarized over all tested neighborhoods. The final KBET score ranges from 0 to 1, with the value 1 indicates the ideal batch mixing.

***Graph connectivity***

The graph connectivity metric(*27*) evaluates whether all cells with the same cell type label are directly connected through a KNN graph. For each cell type label c in the the set C, we created the subset KNN graph $G_c$ to contain only cells belonging to type c and compute the size of the largest connected component. The value is divided by the total number of cells with cell type label c. The graph connectivity score is calculated as the average across all cell types:

$$\textbf{Graph connectivity} = \frac{1}{|C|}\sum_{c \in C} \frac{|LCC(G_c)|}{N_c},$$

where LCC represents the largest connected component, and $N_c$ denotes the total number of cells with cell type label c.

***Aggregated Metrics***

We used three aggregated metrics to evaluate the data integration, where the aggregated metric BioCon computes the average of biological conservation metrics:

$$\textbf{BioCon} = (\textbf{isolate labels} + \textbf{KMeans NMI} + \textbf{KMeans ARI} + \textbf{silhouette label} + \textbf{cLISI})/5,$$

and the aggregate metric BatchCor calculates the average of batch correction metrics:

$$\textbf{BatchCor} = (\textbf{silhouette batch} + \textbf{iLISI} + \textbf{KBET} + \textbf{graph connectivity})/4.$$

The overall metric Total is derived as the weighted average:

$$\textbf{Total} = \mathbf{0.6} * \textbf{BioCon} + \mathbf{0.4} * \textbf{BatchCor}.$$

#### 2. Evaluation metrics in clustering benchmarking

We adopted the evaluation metric calculations in scikit-learn(*80*). Each metric is described below.

***Adjusted Rand index***

The adjusted Rand index (ARI)(*32*) is a metric used to evaluate the similarity between two data clustering results. It improves upon the Rand index (RI), which is a basic measure of agreement between two partitions. The RI considers all pairs of elements and counts the pairs that are either assigned to the same cluster or different clusters in both the predicted and true clustering results. The raw RI score is defined as follows:

$$RI = \frac{\textbf{number of agreeing pairs}}{\textbf{number of pairs}},$$

Scikit-learn adjusted the raw RI score into the ARI score using the following scheme:

$$ARI = \frac{RI - RI_{Expected}}{max(RI) - RI_{Expected}},$$

where $RI_{Expected}$ is the expected value of the RI for random clustering results. The ARI is designed to be close to 0 for random labeling, regardless of the number of clusters or samples, and exactly 1 when the clustering results are identical (up to a permutation).

***Normalized mutual information***

Normalized mutual information (NMI)(*33*) is a normalized version of the mutual information (MI) score, scaling the results between 0 (no mutual information) and 1 (perfect correlation). MI measures the similarity between two sets of labels for the same data, denoted as W and Y, and is defined as follows:

$$MI = \sum_{i=1}^{|W|}\sum_{j=1}^{|Y|}\frac{|W_i \cap Y_i|}{N} log \frac{N|W_i \cap Y_i|}{|W_i||Y_i|},$$

where $|W_i|$ represents the number of the data points in cluster $W_i$. The MI is normalized using an entropy-based method implemented in scikit-learn to produce the NMI score. More details can be found in the 'sklearn.metrics.normalized_mutual_info_score' function. NMI is independent of the absolute label values, meaning that permuting the class or cluster labels does not affect the score.

***Fowlkes-Mallows index***

The Fowlkes-Mallows index (FMI)(*34*) is a metric used to evaluate the similarity between two clustering results by comparing pairs of data points. FMI is based on precision and recall-like metrics, focusing on true positives, false positives, and false negatives, derived from pairwise comparisons of data points in two clustering results, W and Y, as follows:

- True positives (TP): The number of pairs of points that are in the same cluster in both W and Y.
- False positives (FP): The number of pairs of points that are in the same cluster in W but in different clusters in Y.
- False negatives (FN): The number of pairs of points that are in different clusters in W but in the same cluster in Y.

The definition of FMI reads

$$FMI = \frac{TP}{\sqrt{(TP + FP)(TP + FN)}},$$

The FMI score ranges from 0 to 1, where a high value indicates a good similarity between two clustering results.

#### 3. Brief introduction of diffusion in physics

This section is not essential for understanding the graph neural diffusion model but serves as supplementary reading for those interested in deepening their understanding of diffusion theory. Diffusion, a widely recognized physical phenomenon, is intuitively grasped by most people, as seen in examples like heat spreading from a hot area to a cooler one or ink dispersing in water.

The conventional view is that diffusion results from an uneven distribution, such as a temperature gradient, and ends once uniformity is achieved. While this perspective is accurate, it presents an overly simplistic picture.

Using heat diffusion(*38*) as an example, we can explore the complete description of the diffusion process and the concepts of local and global equilibrium states in physics. Heat diffusion begins from a non-equilibrium state, which requires a comprehensive description from both classical and quantum perspectives. In the classical perspective, heat is unevenly distributed throughout the system, causing macroscopic heat flows between different regions. On the other hand, each small element of the heat system, composed of numerous micro-particles, forms a local quantum system (**fig. S22A**). These particles are highly disordered in momentum (or energy) space and have strong local interactions with each other. The non-equilibrium state of the heat system is highly chaotic, making it impossible to define an overall temperature for the entire system and even challenging to determine a localized temperature at a specific point within the system due to quantum physics principles. The disordered micro-particles prevent the establishment of a consistent temperature within the local small quantum system at any specific point.

This situation, where temperature is undefinable, improves during the diffusion process, as it moves the chaotic non-equilibrium system toward equilibrium. The strong local interactions among micro-particles within each quantum element and the macroscopic heat flows between different macro-regions represent the local and global diffusion processes in heat diffusion, respectively. It is important to note that both the local and global diffusion processes begin simultaneously from the outset. However, due to the significant difference in diffusion rates, local and global diffusion lead to distinct equilibrium states.

Local equilibrium within each small element occurs first because of the strong local quantum interactions. In this state, heat (or energy) is sufficiently diffused among the micro-particles, causing them to follow a distribution function in momentum space within each localized small element, allowing the definition of a localized temperature (**fig. S22B**). However, a uniform temperature across the entire system has not yet been achieved, so heat continues to transfer between various macro-regions even in the local equilibrium state. The global equilibrium state occurs after the local equilibrium state, characterized by a consistent temperature throughout the system and the cessation of macroscopic heat flows.

#### 4. Evaluation metrics in trajectory benchmarking

We introduced three trajectory benchmark evaluation metrics, graph edit index(*52*), graph kernel index(*53*) and Jaccard similarity index(*54*), all of which range from 0 to 1.

##### ***Graph edit index***

The graph edit index is derived from the graph edit distance (GED)(*52*), a measure used to quantify the similarity or difference between two graphs. GED is defined as the minimum total cost of the graph edit operations required to transform one graph into another. These edit operations include:

- Node insertion: Adding a new node to the graph.
- Node deletion: Removing an existing node from the graph.
- Node substitution: Changing the label or attributes of a node.

- Edge insertion: Adding a new edge between two nodes.
- Edge deletion: Removing an existing edge between two nodes.
- Edge substitution: Changing the label or attributes of an edge.

The cost of each operation is defined as 1. In our case, all trajectories are tree-shaped graphs, where nodes represent the cell types and edges indicate cell fate transitions. Node and edge substitutions are not required in the graph editing process. It is important to note that the GED from an inferred trajectory to the benchmark trajectory, both of which are tree-shaped, will not exceed the sum of the total number of nodes ($n_{bt}$) and the total number of edges ($e_{bt}$) in the benchmark trajectory. We define the graph edit index as follows:

$$\textbf{Graph edit index} = \frac{n_{bt} + e_{bt} - GED}{n_{bt} + e_{bt}},$$

The graph edit index ranges from 0 to 1, with a value 1 indicating the inferred trajectory exactly matches the benchmark trajectory.

***Graph kernel index***

The graph kernel index relates to graph kernels(*53*), which are functions used to measure the similarity between graphs. This index is applied in various machine learning contexts, particularly with graph-structured data. Graph kernels are designed to capture structural similarities between graphs by comparing their substructures (e.g., walks, paths, subtrees). These kernels map graphs into a high-dimensional space where similarity can be measured using a dot product. In this work, we compute the graph kernel index using the Python package GraKeL with the shortest path kernel type.

***Jaccard similarity index***

The Jaccard similarity index(*54*), also known as the Jaccard index or Jaccard coefficient, is a statistical measure used to compare the similarity and diversity of sample sets. It is a widely used metric in various fields, including data science, ecology, and information retrieval. We use the Jaccard similarity index to compare the similarity of the edges between the inferred and benchmark trajectory graphs. The Jaccard similarity index is defined as follows:

$$\textbf{Jaccard similarity index} = \frac{|E_{it} \cap E_{bt}|}{|E_{it} \cup E_{bt}|},$$

where $E_{it}$ represent the set of edges for the inferred trajectory and $E_{bt}$ represent the set of edges for the benchmark trajectory. $|E_{it} \cap E_{bt}|$ is the number of elements common to both sets (the intersection), and $|E_{it} \cup E_{bt}|$ is the number of elements present in either set (the union). A higher Jaccard similarity index indicates a greater degree of similarity between the two sets.

#### 5. Comparing CellDiffusion with UniPort and BBKNN

We compared CellDiffusion with UniPort and BBKNN. We noticed that UniPort was primarily developed for aligning unpaired single cell datasets across modality settings and heterogeneous domains, which does not aim to construct a unified scRNA-seq latent space that supports standard downstream analyses such as clustering or marker identification. However, we have successfully run UniPort integration pipeline using only scRNA-seq data. We found UniPort did not integrate different batches for the Endometrium-CD dataset (including samples GSM6605435 and GSM6605434 from Endometrium-314k dataset) (**fig. S16A**). We verified it using another human bone marrow mononuclear cell dataset, where the two batches are also clearly separated in UniPort's umap (**fig. S16B**), though the two batches contain most common cell types. These results show UniPort does not outperform the competitive methods for pure scRNA-seq data integration in this specific scenario.

For BBKNN, we emphasize that it does not change cell embeddings in the integration process and cannot be evaluated using the batch correction and biological conservation metrics in (*27*). However, we compared the UMAPs generated from CellDiffusion embeddings with those from the BBKNN graph using BMMC dataset (**fig. S9 and S17**). The UMAPs show CellDiffusion outperformed BBKNN in distinguishing IGKC+/- B cells.

#### 6. From graph neural networks to CellDiffusion

Graph neural networks (GNNs)(*56, 57*) are capable of directly handling graph structures, enabling feature exchange between nodes and dimensionality reduction in each layer. However, a typical GNN, such as graph convolutional network(*56*) and graph attention network(*57*), generally contains only two feature aggregation layers (**fig. S2E**). When more diffusion layers are added, these networks often face challenges like over-smoothing(*59, 60*) and information over-squashing(*61*). This occurs because GNNs usually perform simple and coarse feature exchanges based solely on adjacency or attention matrices, which fail to effectively capture important signals from large interaction fields for each node.

In graph learning method, the number of indirectly linked nodes increases exponentially with each additional layer (**fig. S2E**). This leads to rough information exchanges between too many nodes, making their features increasingly similar and eventually indistinguishable, resulting in the over-smoothing problem(*59, 60*). Over-smoothing also occurs in densely connected graphs, where excessive direct feature exchanges further compound the issue.

Additionally, GNNs often struggle to effectively extract relevant information from a source signal about a target signal when aggregating information over long paths, leading to the over-squashing of exponentially growing information into the fixed-size features of individual nodes(*61*). This makes it difficult for GNNs to propagate messages from distant nodes, resulting in poor performance on tasks that depend on long-range interactions. These issues of over-smoothing and information over-squashing result in the deterioration of GNN performance in deeper networks.

CellDiffusion facilitates long-range interactions by incorporating more diffusion layers (see Chamberlain et al.(*23*) for the effectiveness of graph neural diffusion with 1-32 layers), making it much deeper than typical GNNs. This significantly increases the diffusion depth, as the interaction field of a node expands exponentially with each additional layer (**fig. S2E**). Despite the depth, CellDiffusion maintains a detailed and accurate diffusion process. This is achieved through the core partial differential equation governing the model, along with the discretization choices of temporal and spatial operators. With this, the feature flow spreads on the single-cell graph like the heat diffusion process in physics (**fig. S2A**). Furthermore, a graph rewiring process is introduced at each layer to capture and refine the key structures of the single-cell graph. CellDiffusion also maintains consistent data dimensionality throughout the diffusion process, enabling all layers to share the same attention matrix. This not only reduces computational overhead but also enhances the cell-cell attention network, accurately identifying cell-cell interactions and effectively extracting relevant information from a source signal about a target signal even across long paths.

CellDiffusion not only spreads node features farther across the single-cell graph but also enables the handling of densely connected graphs, thanks to its stable and precise diffusion process, allowing for more direct feature exchanges. These features meet the demands of scRNA-seq analysis, where single-cell graphs often exhibit diverse structures at different scales (**fig. S2D**). Within any given scale, finer structures can be observed at sub-scales, capturing rich biological information such as coarse and fine cell types, as well as broad and detailed lineage relationships.

CellDiffusion enhances the detection of information at both small and large scales, enabling better capture of graph structures across varying scales.

**7. Model collapse diagnostics at deep layers**

We carried out model collapse diagnostics and compared CellDiffusion with GCN at different network layers (2-16 layers). For the GCN model, we just use GCN to replace the diffusion module in CellDiffusion. The GCN model uses the same preprocessed scRNA-seq data, auto-encoded cell embeddings, single-cell graph for integration and loss function, as that used in CellDiffusion. We used three model collapse diagnostic metrics: neighbor purity, intrinsic dimension and variance explained by embeddings. Neighbor purity evaluates the percentage of the cells with the same cell type as each estimated cell in its KNN neighborhoods. Intrinsic dimension evaluates the effective dimensions of cell embeddings that explaining 95% variance of all cell embeddings. Variance explained by embeddings evaluates the percentage of variance of the preprocessed scRNA-seq data (includes 2000 variable genes) that has been explained by cell embeddings.

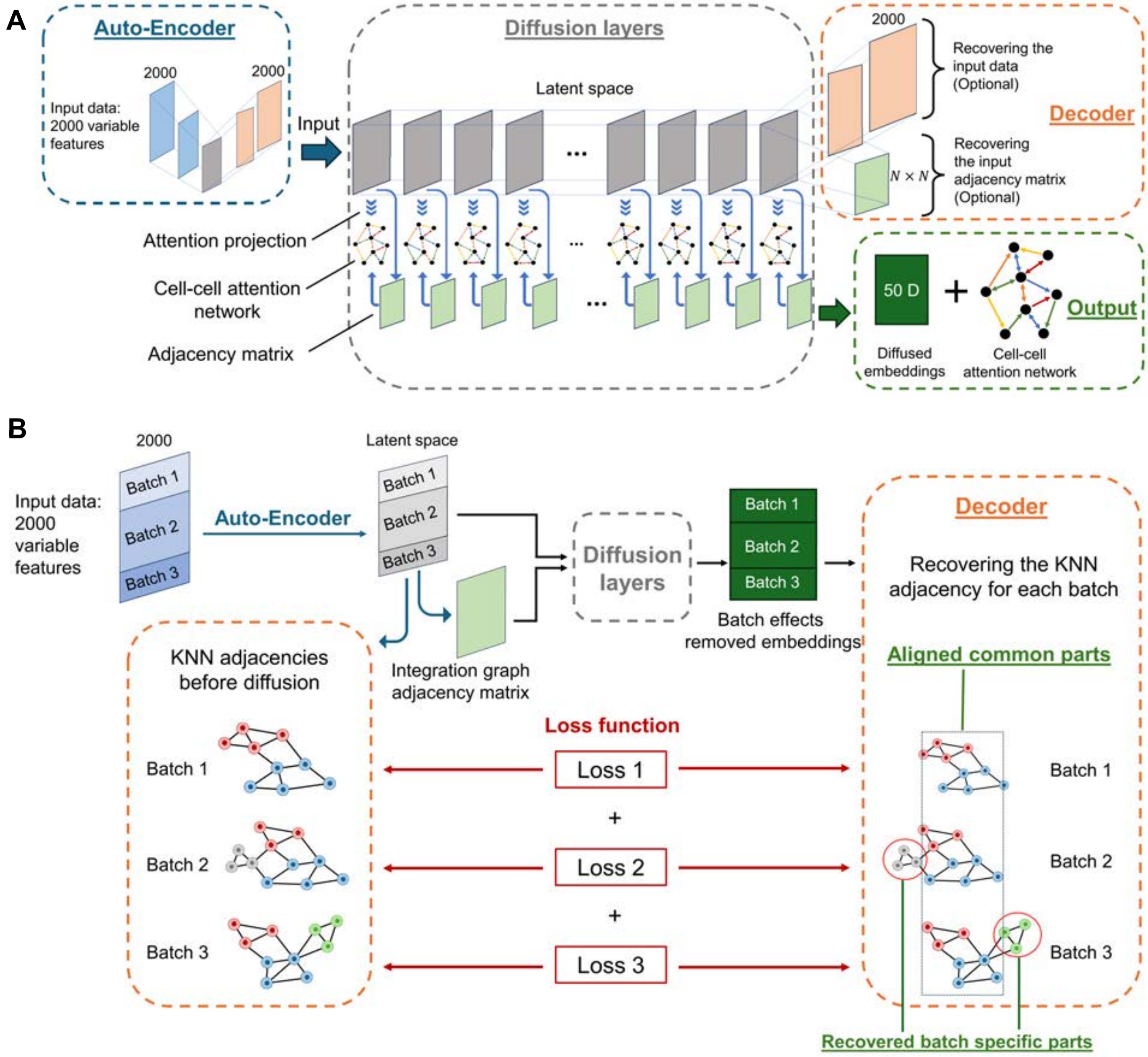


**Fig. S1. CellDiffusion network architecture and data integration workflow.** (**A**) The architecture of CellDiffusion network. (**B**) Data integration workflow. CellDiffusion reconstructs a KNN graph within each batch to ensure that intra-batch structures are preserved after batch correction, especially for those batch specific biological signals.

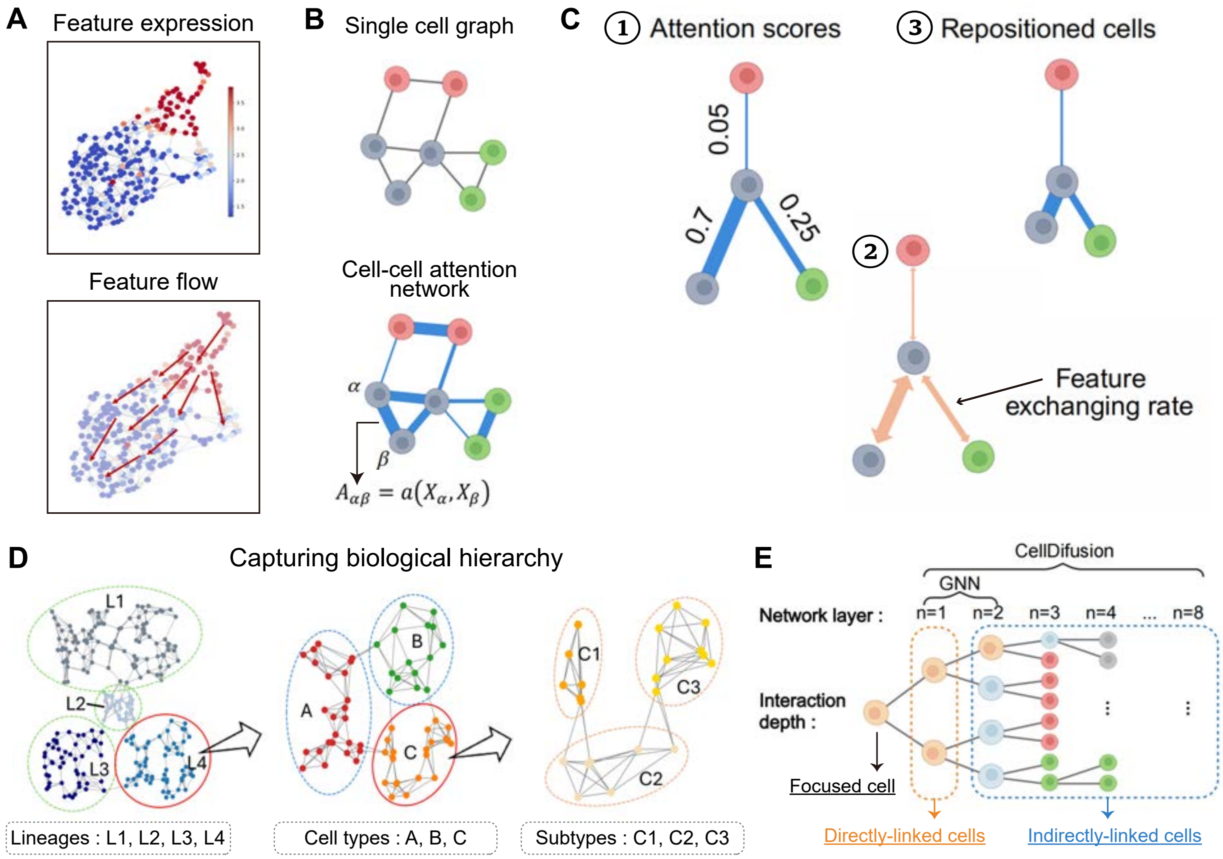


**Fig. S2. Schematic illustration of feature diffusion steps in CellDiffusion.** (**A**) Feature flow spreads from cells with high expression to cells with low expression through linked edges. (**B**) Training a cell-cell attention network on a single cell graph. (**C**) Attentions guide feature diffusion rates and cell alignments. (**D**) CellDiffusion captures the biological hierarchy from large lineages to fine-grained subtypes. (**E**) Feature diffusion depth in CellDiffusion.

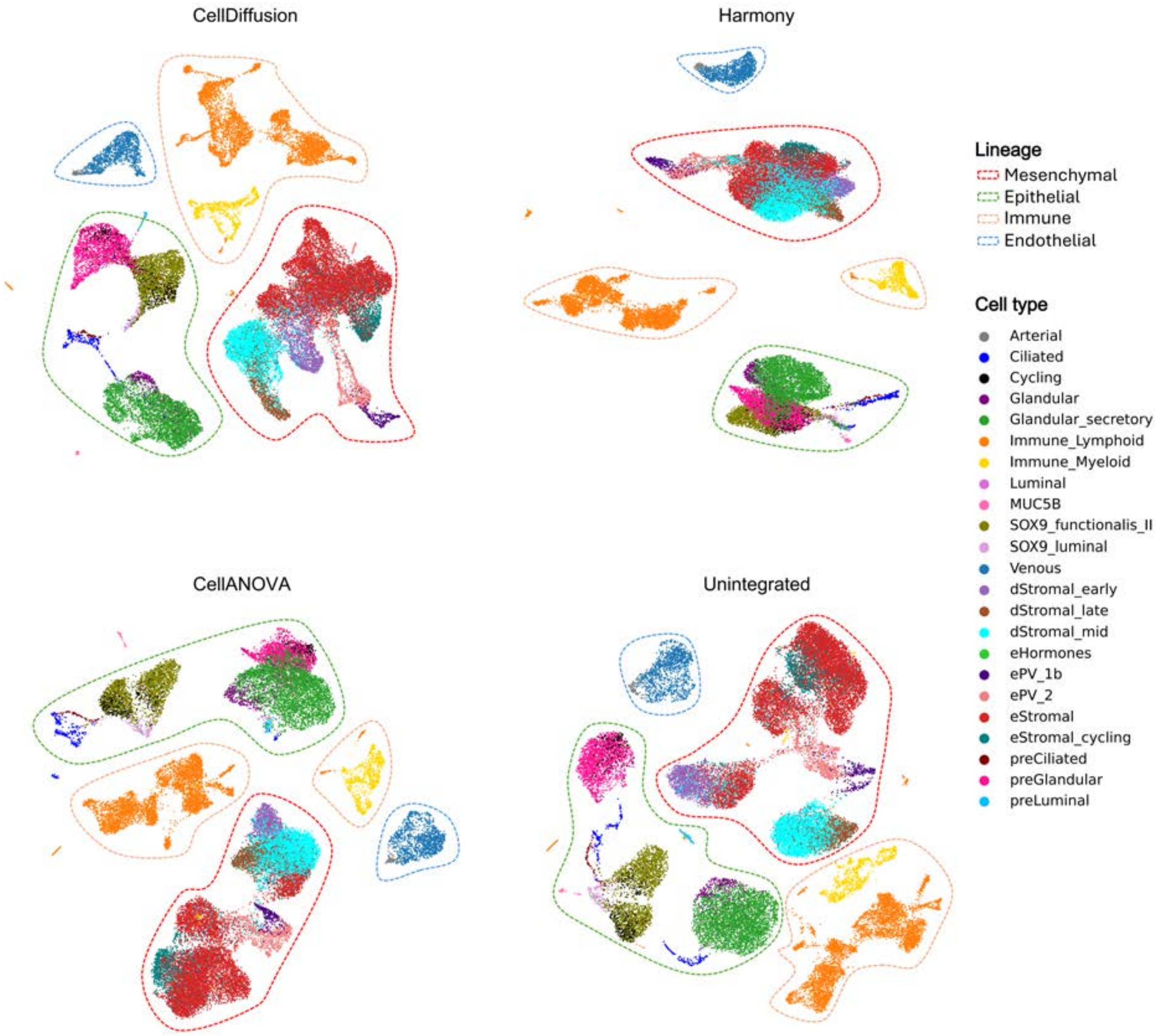


**Fig. S3. UMAP plots showing integration results from CellDiffusion, Harmony, and CellANOVA compared to the unintegrated Endometrium-35K dataset.** Different lineages were circled and highlighted to facilitate cross-method comparison.

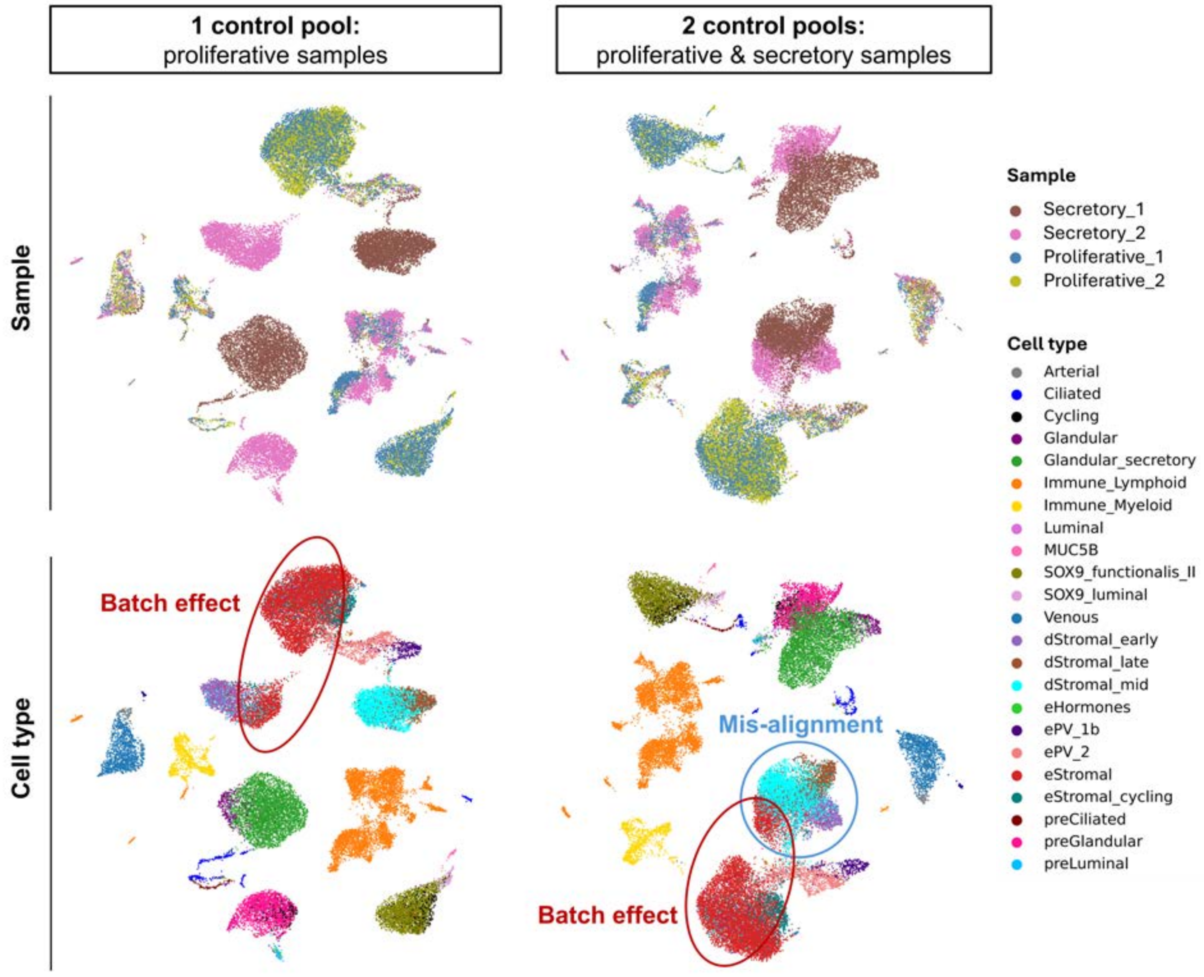


**Fig. S4. UMAP plots showing CellANOVA integration results with alternative control pool settings on the Endometrium-35K dataset.** UMAP plots at the top are colored by samples, and those at the bottom are colored by cell types. UMAP plots on the left were generated using proliferative samples as the only control pool, while those on the right used both proliferative and secretory samples as control pools.

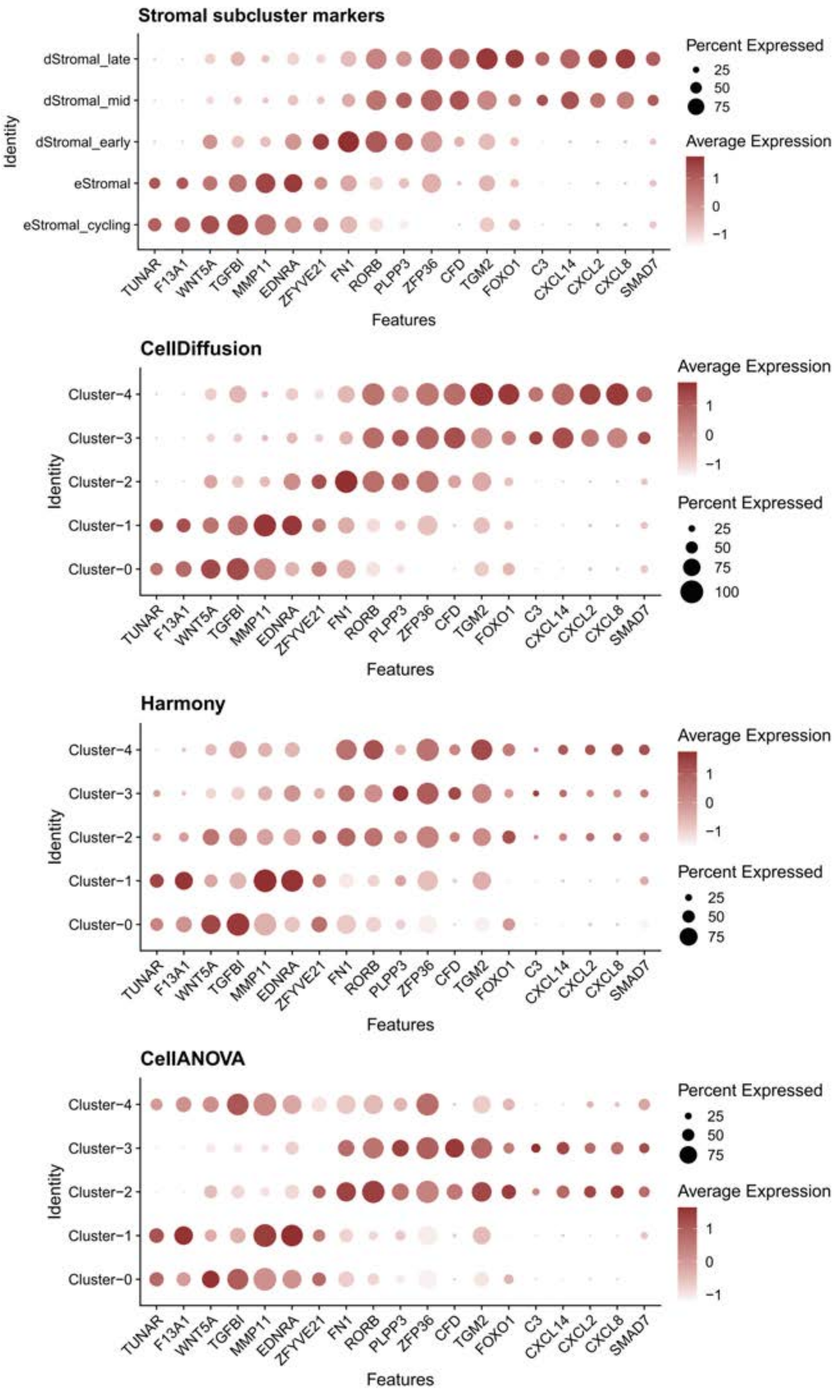


**Fig. S5. Stromal subtype marker expression across clusters based on the integration results of different methods applied to the Endometrium-35k dataset.** From top to bottom: manually curated cell types from the original study (*28*), and clustering results obtained using Scanpy applied to the integration outputs of CellDiffusion, Harmony, and CellANOVA.

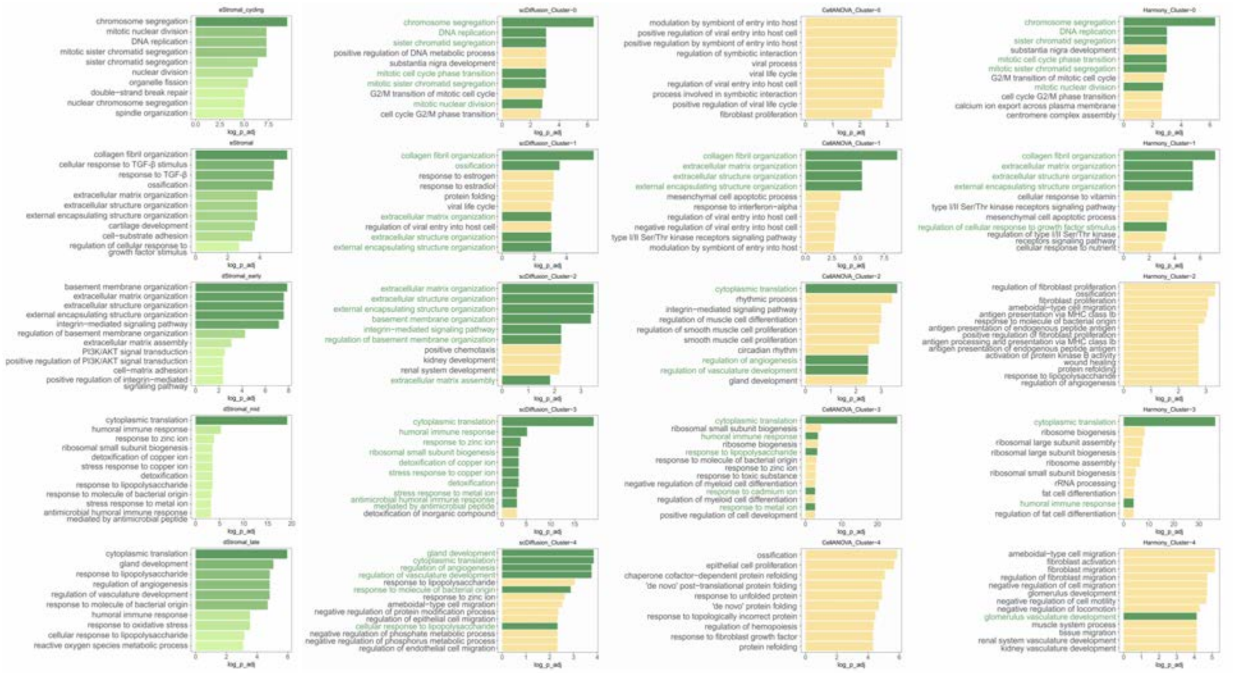

**Fig. S6. Comparison of functional enrichment analysis results from different data integration methods applied to the Endometrium-35K dataset.** Differential expression analysis was performed on clusters defined by manually curated cell types from the original study, as well as clustering results from CellDiffusion, Harmony, and CellANOVA (from left to right). Only the top 100 cluster-specific genes were used for the analysis.

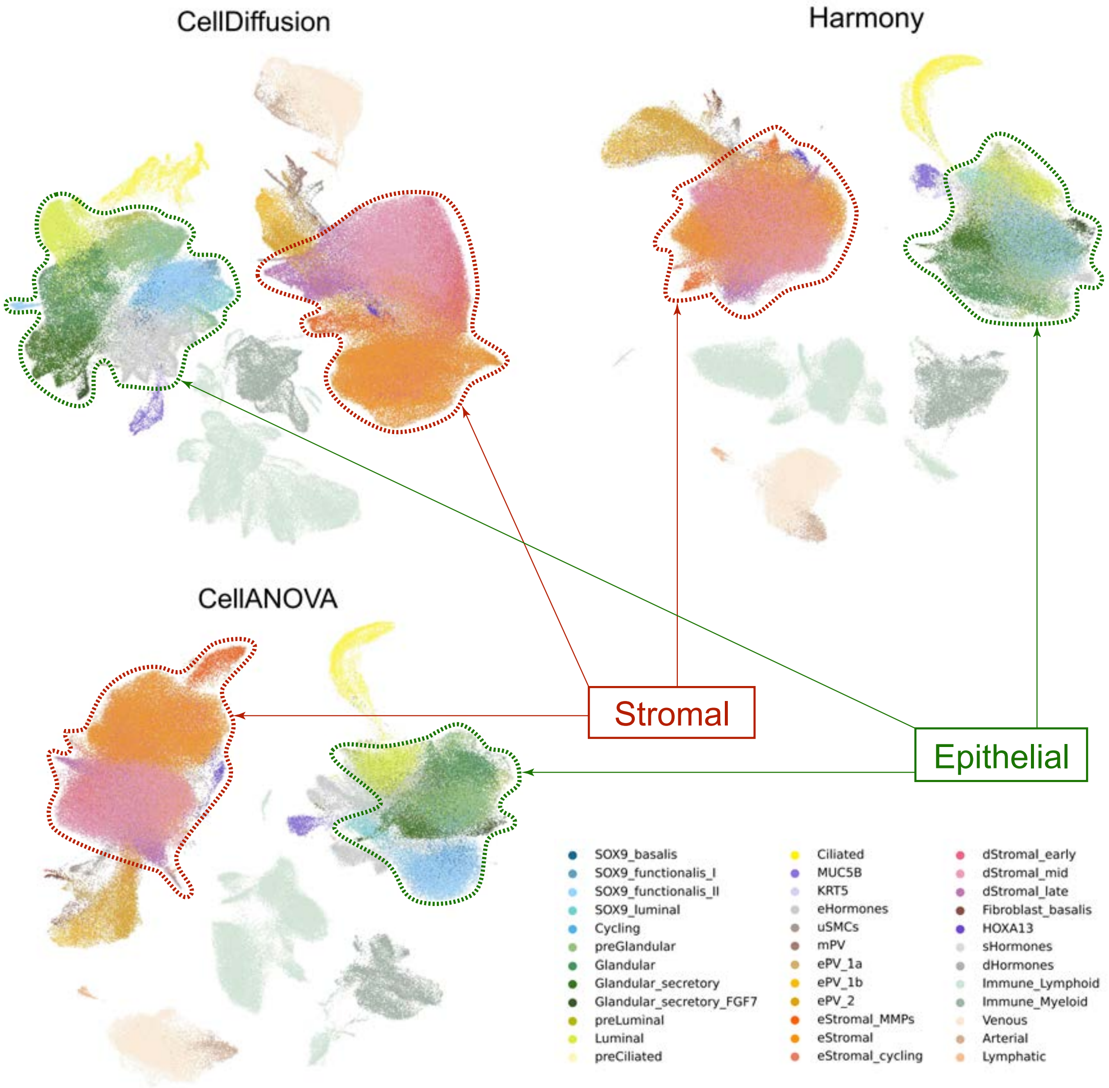


**Fig. S7. UMAP plots showing integration results of CellDiffusion, Harmony, and CellANOVA on the Endometrium-314K dataset.** 'Stromal' and 'glandular-luminal epithelial' cell populations were highlighted for comparison.

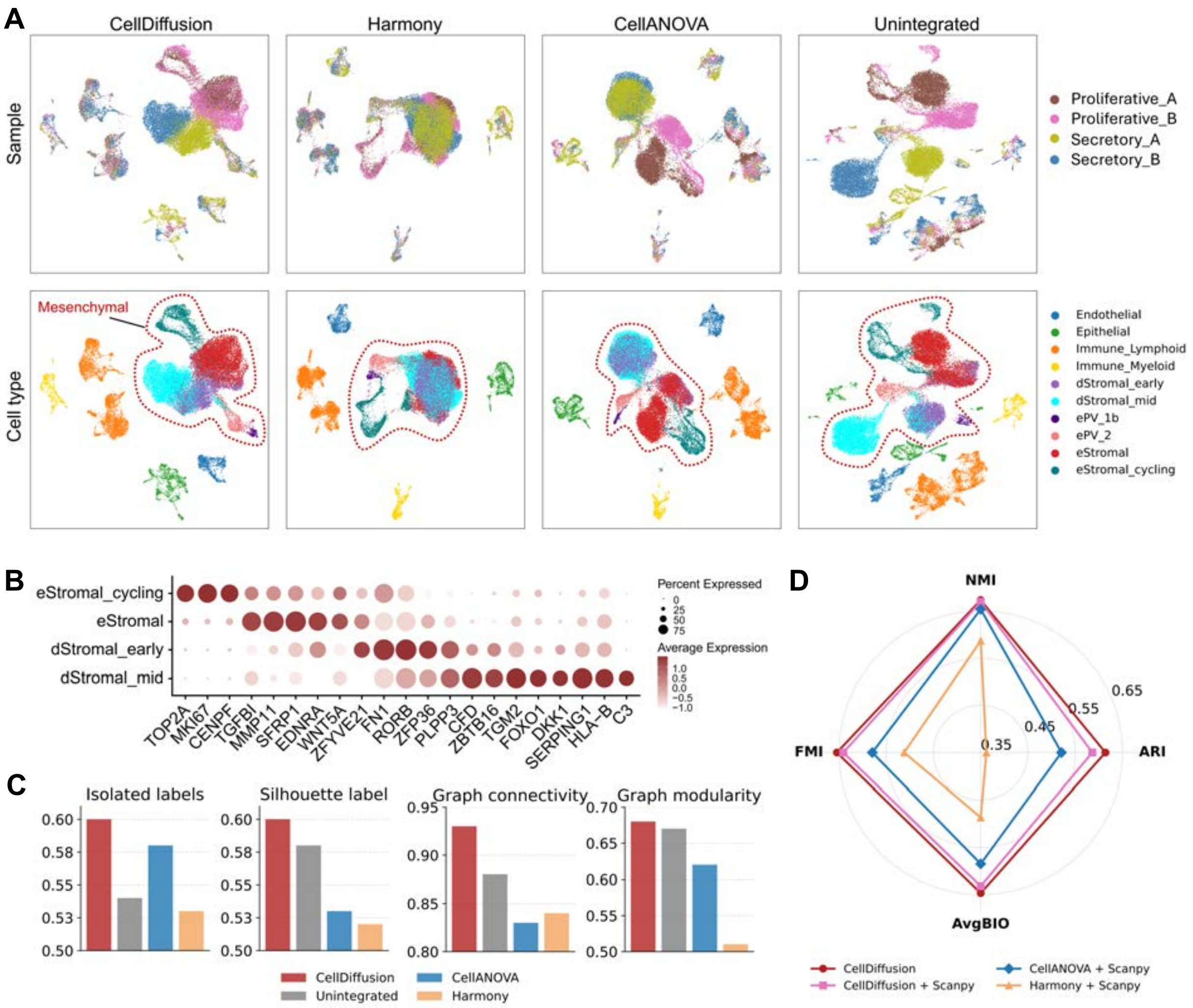


**Fig. S8. Data integration on the Endometrium-EV dataset.** (**A**) UMAP plots of samples and cell types for different tools. (**B**) Dot-plot of stromal cell subtype markers. (**C**) Data integration evaluation. (**D**) Clustering evaluation.

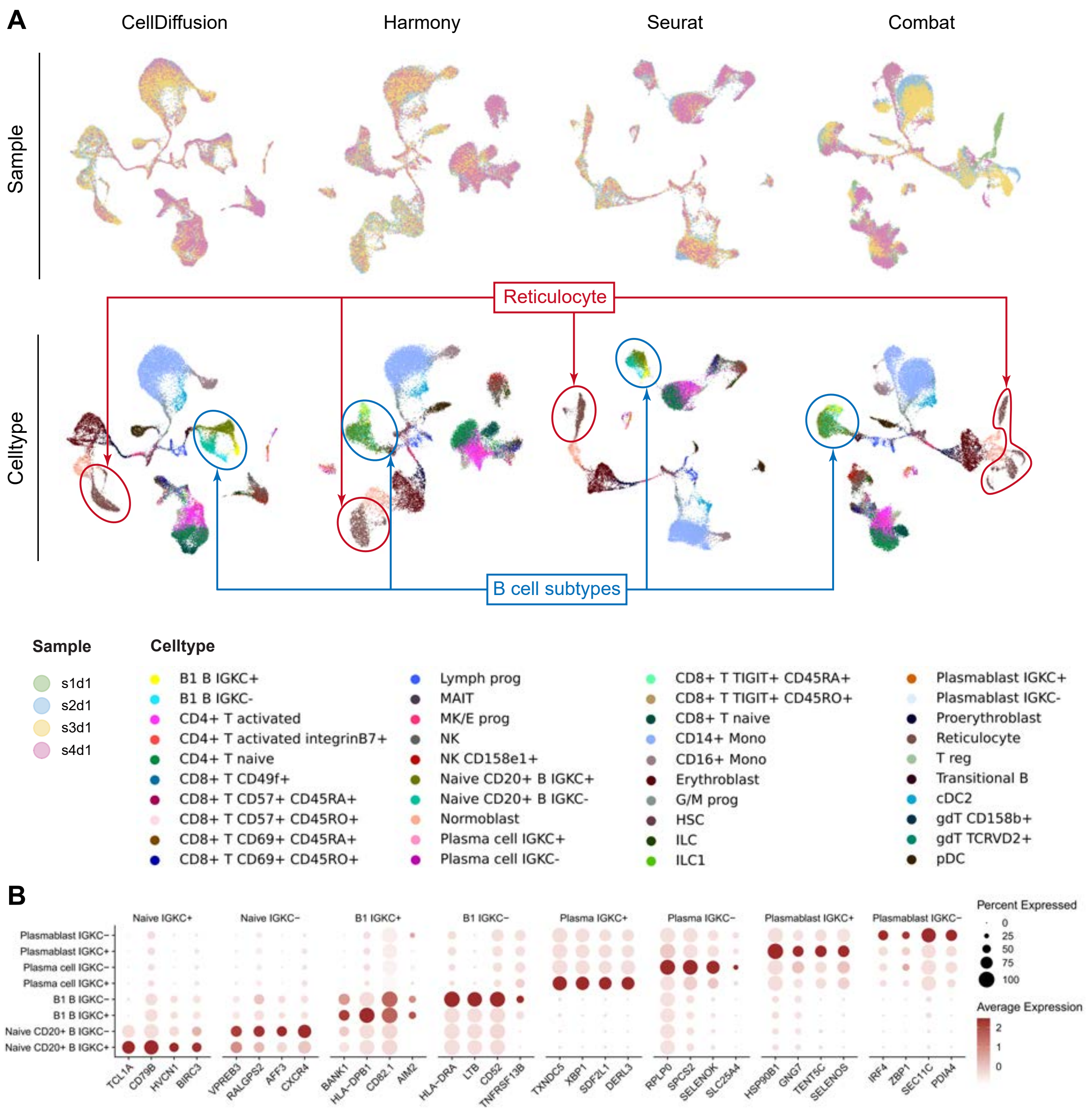


**Fig. S9. Compare different tools using the BMMC dataset.** (**A**) UMAP plots of cell types for different tools. (**B**) Distinct marker gene expression profiles for the eight IGKC+ subpopulations.

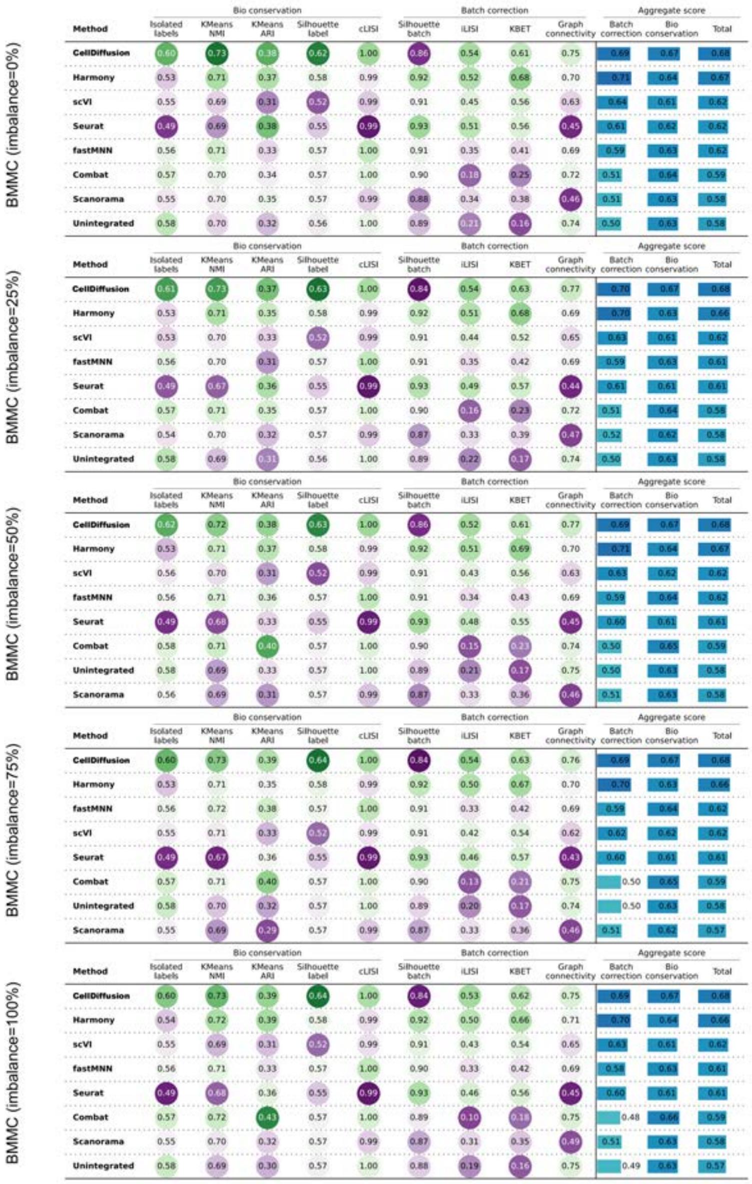


**Fig. S10. scRNA-seq integration benchmark results on the BMMC dataset (table generated using scib-metrics).** CellDiffusion was benchmarked with Harmony, scVI, Seurat, fastMNN, Combat, Scanorama and Unintegrated embeddings for data integration performance. Unintegrated embeddings are obtained from Scanpy PCA process after the normalization. BMMC contains 40 cell types and 4 batches (s1d1, s2d1, s3d1, s4d1). The cell type imbalance is introduced by manually removing CD4+ T activated cells from batches s1d1 and s2d1and Erythroblast cells from batches s3d1 and s4d1 at different percentages (0%, 25%, 50%, 75% and 100%). We assessed the performance of all methods in terms of biological conservation and batch correction using five detailed metrics for biological conservation and four for batch correction.

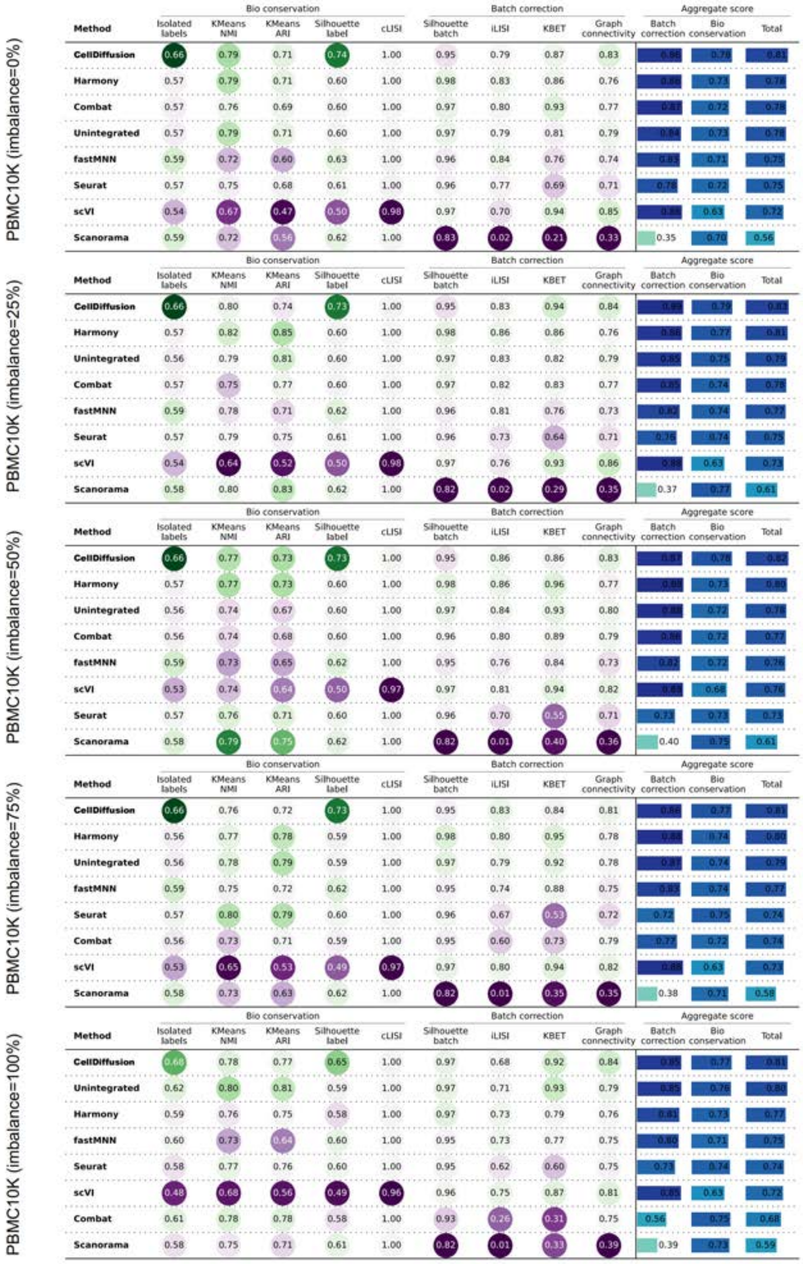


**Fig. S11. scRNA-seq integration benchmark results on the PBMC-10k dataset (table generated using scib-metrics).** CellDiffusion was benchmarked with Harmony, scVI, Seurat, fastMNN, Combat, Scanorama and Unintegrated embeddings for data integration performance. Unintegrated embeddings are obtained from Scanpy PCA process after the normalization. PBMC-10k contains 9 cell types and 2 batches. The cell type imbalance is introduced by manually removing CD4 T cells from one batch at different percentages (0%, 25%, 50%, 75% and 100%). We assessed the performance of all methods in terms of biological conservation and batch correction using five detailed metrics for biological conservation and four for batch correction.

**PBMC23K (imbalance=0%)**

| Method | Bio conservation: Isolated labels | KMeans NMI | KMeans ARI | Silhouette label | cLISI | Batch correction: Silhouette batch | iLISI | KBET | Graph connectivity | Aggregate score: Batch correction | Bio conservation | Total |
|---|---|---|---|---|---|---|---|---|---|---|---|---|
| CellDiffusion | 0.65 | 0.74 | 0.45 | 0.66 | 1.00 | 0.87 | 0.77 | 0.76 | 0.84 | 0.81 | 0.70 | 0.74 |
| Harmony | 0.53 | 0.72 | 0.42 | 0.60 | 1.00 | 0.91 | 0.79 | 0.89 | 0.85 | 0.86 | 0.65 | 0.73 |
| scVI | 0.64 | 0.69 | 0.33 | 0.57 | 1.00 | 0.92 | 0.76 | 0.85 | 0.91 | 0.86 | 0.65 | 0.73 |
| Seurat | 0.46 | 0.73 | 0.46 | 0.61 | 1.00 | 0.88 | 0.70 | 0.82 | 0.80 | 0.80 | 0.65 | 0.71 |
| Unintegrated | 0.54 | 0.71 | 0.40 | 0.59 | 1.00 | 0.90 | 0.66 | 0.77 | 0.85 | 0.79 | 0.65 | 0.71 |
| fastMNN | 0.57 | 0.69 | 0.34 | 0.59 | 1.00 | 0.92 | 0.53 | 0.74 | 0.82 | 0.75 | 0.64 | 0.68 |
| Combat | 0.54 | 0.72 | 0.45 | 0.59 | 1.00 | 0.90 | 0.41 | 0.61 | 0.85 | 0.69 | 0.66 | 0.67 |
| Scanorama | 0.51 | 0.71 | 0.40 | 0.62 | 1.00 | 0.81 | 0.55 | 0.68 | 0.76 | 0.70 | 0.65 | 0.67 |

**PBMC23K (imbalance=25%)**

| Method | Bio conservation: Isolated labels | KMeans NMI | KMeans ARI | Silhouette label | cLISI | Batch correction: Silhouette batch | iLISI | KBET | Graph connectivity | Aggregate score: Batch correction | Bio conservation | Total |
|---|---|---|---|---|---|---|---|---|---|---|---|---|
| CellDiffusion | 0.69 | 0.74 | 0.47 | 0.66 | 1.00 | 0.88 | 0.77 | 0.75 | 0.86 | 0.82 | 0.71 | 0.75 |
| Harmony | 0.53 | 0.72 | 0.42 | 0.60 | 1.00 | 0.91 | 0.79 | 0.89 | 0.85 | 0.86 | 0.65 | 0.73 |
| scVI | 0.56 | 0.70 | 0.36 | 0.56 | 1.00 | 0.91 | 0.76 | 0.81 | 0.90 | 0.84 | 0.64 | 0.72 |
| Unintegrated | 0.54 | 0.71 | 0.40 | 0.59 | 1.00 | 0.90 | 0.67 | 0.78 | 0.85 | 0.80 | 0.65 | 0.71 |
| Seurat | 0.49 | 0.72 | 0.42 | 0.60 | 1.00 | 0.88 | 0.66 | 0.75 | 0.80 | 0.77 | 0.65 | 0.70 |
| fastMNN | 0.57 | 0.68 | 0.33 | 0.59 | 1.00 | 0.92 | 0.52 | 0.75 | 0.82 | 0.75 | 0.63 | 0.68 |
| Combat | 0.53 | 0.71 | 0.41 | 0.59 | 1.00 | 0.90 | 0.39 | 0.62 | 0.84 | 0.69 | 0.65 | 0.66 |
| Scanorama | 0.37 | 0.71 | 0.41 | 0.60 | 1.00 | 0.81 | 0.28 | 0.66 | 0.64 | 0.60 | 0.62 | 0.61 |

**PBMC23K (imbalance=50%)**

| Method | Bio conservation: Isolated labels | KMeans NMI | KMeans ARI | Silhouette label | cLISI | Batch correction: Silhouette batch | iLISI | KBET | Graph connectivity | Aggregate score: Batch correction | Bio conservation | Total |
|---|---|---|---|---|---|---|---|---|---|---|---|---|
| CellDiffusion | 0.70 | 0.72 | 0.40 | 0.65 | 1.00 | 0.87 | 0.76 | 0.76 | 0.84 | 0.81 | 0.69 | 0.74 |
| Harmony | 0.53 | 0.70 | 0.37 | 0.60 | 1.00 | 0.91 | 0.78 | 0.88 | 0.85 | 0.85 | 0.64 | 0.73 |
| scVI | 0.59 | 0.68 | 0.33 | 0.56 | 1.00 | 0.92 | 0.75 | 0.85 | 0.90 | 0.86 | 0.63 | 0.72 |
| Unintegrated | 0.54 | 0.71 | 0.40 | 0.59 | 1.00 | 0.90 | 0.65 | 0.78 | 0.86 | 0.82 | 0.65 | 0.71 |
| Seurat | 0.49 | 0.72 | 0.41 | 0.60 | 1.00 | 0.88 | 0.67 | 0.77 | 0.81 | 0.78 | 0.64 | 0.70 |
| fastMNN | 0.57 | 0.69 | 0.33 | 0.59 | 1.00 | 0.92 | 0.51 | 0.75 | 0.82 | 0.75 | 0.64 | 0.68 |
| Combat | 0.53 | 0.71 | 0.41 | 0.59 | 1.00 | 0.90 | 0.36 | 0.60 | 0.85 | 0.67 | 0.65 | 0.66 |
| Scanorama | 0.38 | 0.71 | 0.39 | 0.60 | 1.00 | 0.82 | 0.28 | 0.60 | 0.62 | 0.58 | 0.62 | 0.60 |

**PBMC23K (imbalance=75%)**

| Method | Bio conservation: Isolated labels | KMeans NMI | KMeans ARI | Silhouette label | cLISI | Batch correction: Silhouette batch | iLISI | KBET | Graph connectivity | Aggregate score: Batch correction | Bio conservation | Total |
|---|---|---|---|---|---|---|---|---|---|---|---|---|
| CellDiffusion | 0.66 | 0.72 | 0.39 | 0.66 | 1.00 | 0.84 | 0.77 | 0.71 | 0.83 | 0.79 | 0.68 | 0.73 |
| Harmony | 0.53 | 0.71 | 0.40 | 0.60 | 1.00 | 0.91 | 0.75 | 0.86 | 0.83 | 0.84 | 0.65 | 0.72 |
| scVI | 0.58 | 0.70 | 0.36 | 0.56 | 1.00 | 0.92 | 0.73 | 0.85 | 0.88 | 0.85 | 0.64 | 0.72 |
| Unintegrated | 0.55 | 0.71 | 0.40 | 0.59 | 1.00 | 0.90 | 0.61 | 0.79 | 0.83 | 0.78 | 0.65 | 0.70 |
| Seurat | 0.49 | 0.70 | 0.36 | 0.60 | 1.00 | 0.88 | 0.69 | 0.74 | 0.82 | 0.78 | 0.63 | 0.69 |
| fastMNN | 0.57 | 0.69 | 0.35 | 0.59 | 1.00 | 0.92 | 0.50 | 0.75 | 0.82 | 0.75 | 0.64 | 0.68 |
| Combat | 0.53 | 0.68 | 0.34 | 0.59 | 1.00 | 0.90 | 0.30 | 0.55 | 0.85 | 0.65 | 0.63 | 0.64 |
| Scanorama | 0.38 | 0.72 | 0.41 | 0.60 | 1.00 | 0.82 | 0.27 | 0.64 | 0.64 | 0.59 | 0.62 | 0.61 |

**PBMC23K (imbalance=100%)**

| Method | Bio conservation: Isolated labels | KMeans NMI | KMeans ARI | Silhouette label | cLISI | Batch correction: Silhouette batch | iLISI | KBET | Graph connectivity | Aggregate score: Batch correction | Bio conservation | Total |
|---|---|---|---|---|---|---|---|---|---|---|---|---|
| CellDiffusion | 0.57 | 0.75 | 0.51 | 0.64 | 1.00 | 0.87 | 0.70 | 0.74 | 0.87 | 0.80 | 0.69 | 0.73 |
| scVI | 0.61 | 0.69 | 0.33 | 0.56 | 1.00 | 0.91 | 0.69 | 0.81 | 0.92 | 0.83 | 0.64 | 0.72 |
| Harmony | 0.55 | 0.70 | 0.39 | 0.60 | 1.00 | 0.90 | 0.71 | 0.82 | 0.84 | 0.82 | 0.65 | 0.72 |
| Seurat | 0.54 | 0.69 | 0.34 | 0.60 | 1.00 | 0.88 | 0.65 | 0.77 | 0.80 | 0.78 | 0.63 | 0.69 |
| Unintegrated | 0.56 | 0.68 | 0.31 | 0.59 | 1.00 | 0.90 | 0.51 | 0.79 | 0.84 | 0.76 | 0.63 | 0.68 |
| fastMNN | 0.57 | 0.68 | 0.31 | 0.58 | 1.00 | 0.91 | 0.48 | 0.76 | 0.81 | 0.74 | 0.63 | 0.67 |
| Combat | 0.55 | 0.70 | 0.36 | 0.59 | 1.00 | 0.89 | 0.23 | 0.47 | 0.86 | 0.61 | 0.64 | 0.63 |
| Scanorama | 0.48 | 0.70 | 0.35 | 0.61 | 1.00 | 0.77 | 0.32 | 0.65 | 0.63 | 0.59 | 0.63 | 0.61 |

**Fig. S12. scRNA-seq integration benchmark results on the PBMC-23k dataset (table generated using scib-metrics).** CellDiffusion was benchmarked with Harmony, scVI, Seurat, fastMNN, Combat, Scanorama and Unintegrated embeddings for data integration performance. Unintegrated embeddings are obtained from Scanpy PCA process after the normalization. PBMC-23k contains 30 cell types and 3 batches (P8_0, P8_3, P8_7). The cell type imbalance is introduced by manually removing CD4 Naive cells from batch P8_0, and NK cells from batch P8_3 at different percentages (0%, 25%, 50%, 75% and 100%). We assessed the performance of all methods in terms of biological conservation and batch correction using five detailed metrics for biological conservation and four for batch correction.

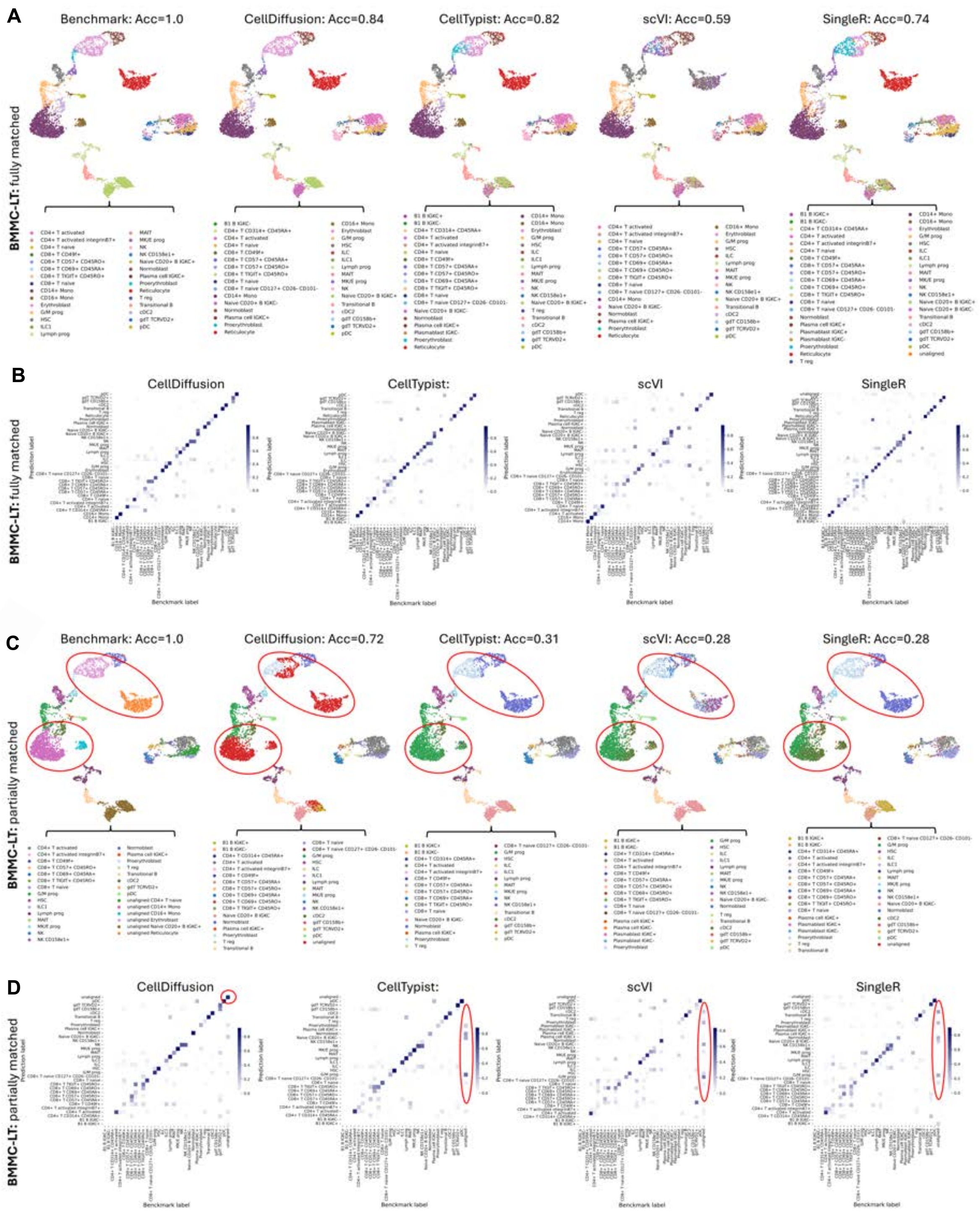


**Fig. S13. Results of cell type annotation on the BMMC-LT dataset.** (**A**) Annotation result using a fully matched reference. (**B**) Confusion matrix for the fully matched reference. (**C**) Annotation result using a partially matched reference. (**D**) Confusion matrix the partially matched reference.

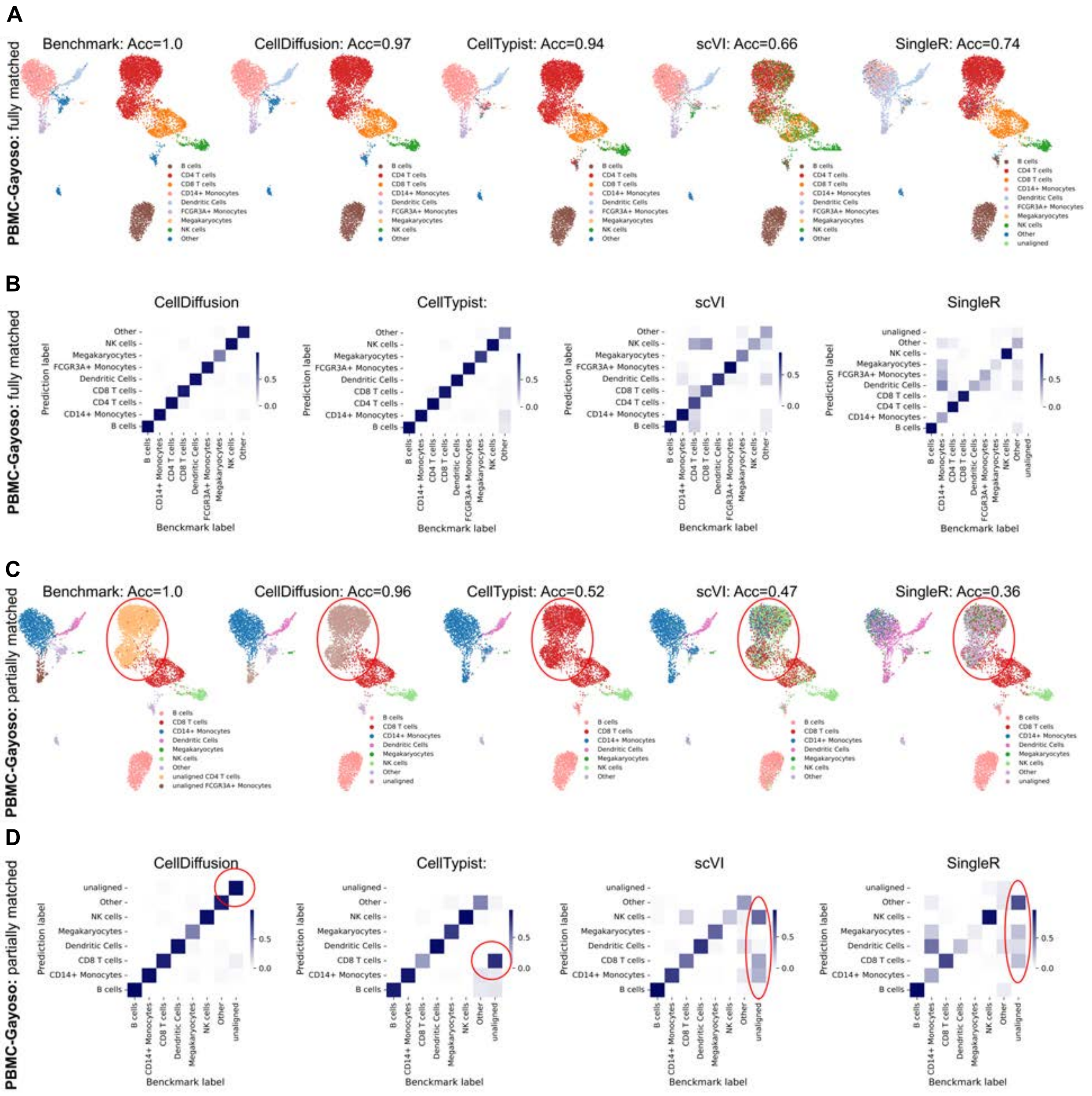


**Fig. S14. Results of cell type annotation on the PBMC-Gayoso.** (**A**) Annotation result using a fully matched reference. (**B**) Confusion matrix for the fully matched reference. (**C**) Annotation result using a partially matched reference. (**D**) Confusion matrix for the partially matched reference.

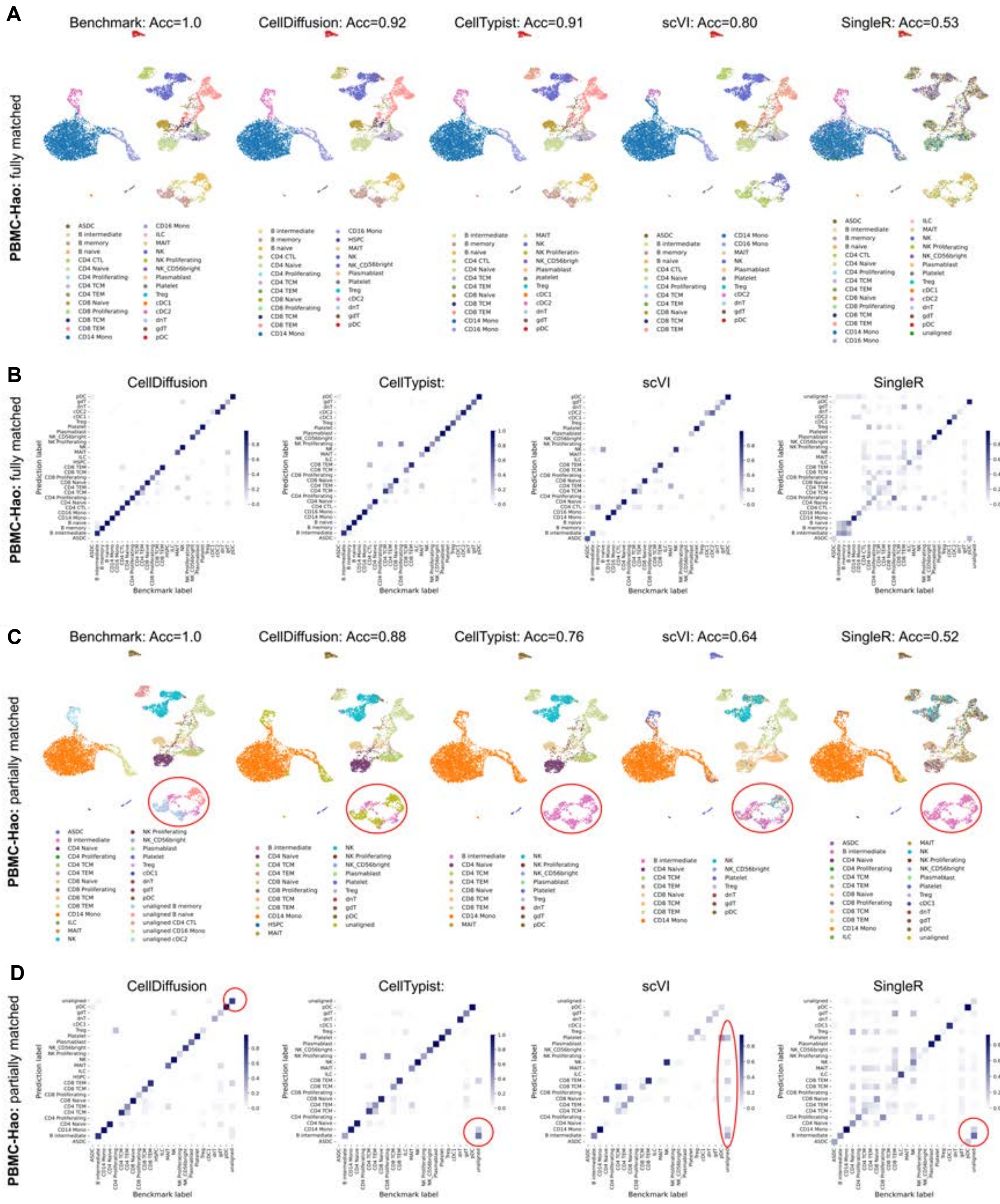


**Fig. S15. Results of cell type annotation on the PBMC-Hao.** (**A**) Annotation result using a fully matched reference. (**B**) Confusion matrix for the fully matched reference. (**C**) Annotation result using a partially matched reference. (**D**) Confusion matrix for the partially matched reference.

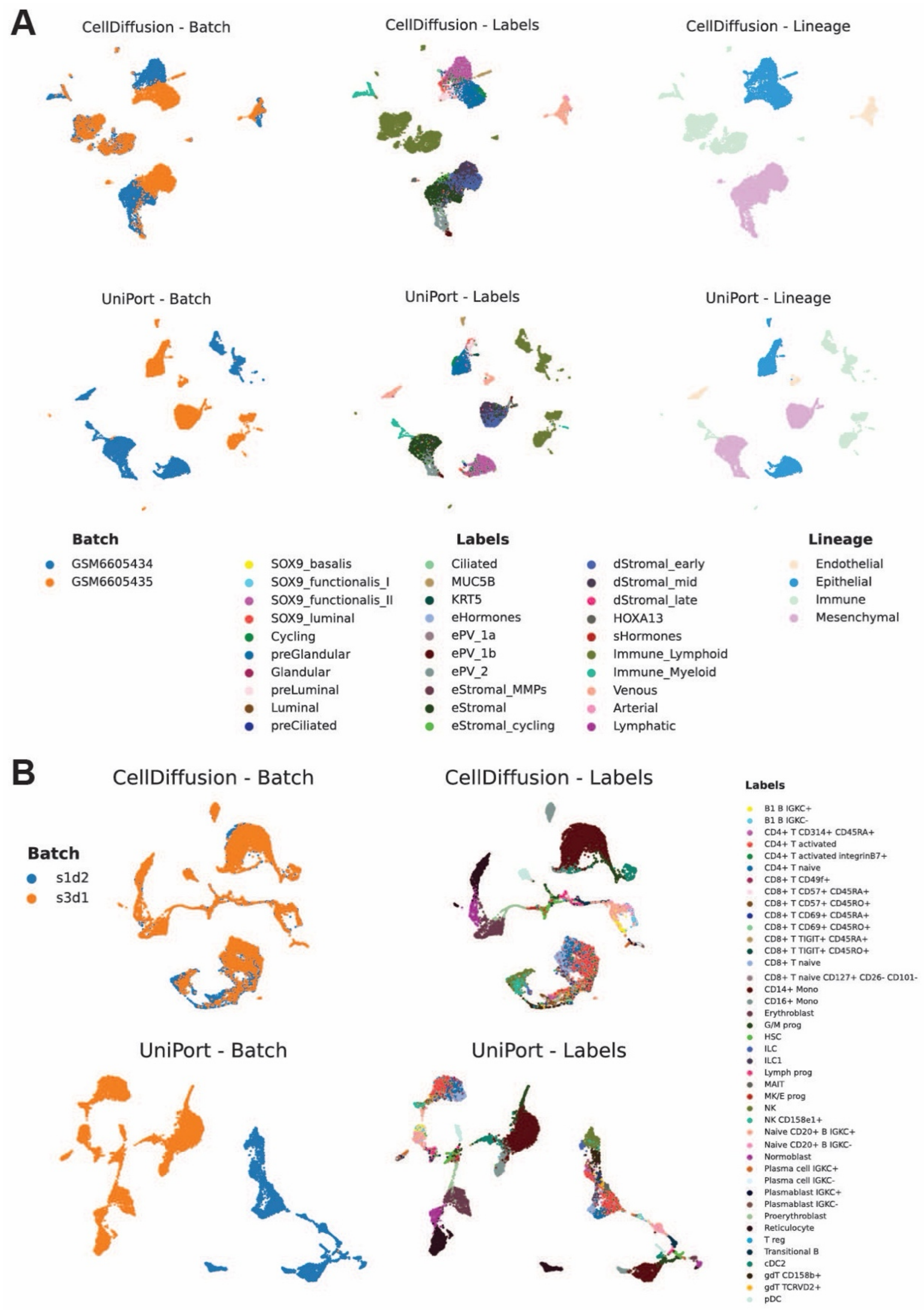


**Fig. S16. UMAP comparison between CellDiffusion and UniPort.** (**A**) UMAP visualizations on the Endometrium-CD dataset. (**B**) UMAP comparison on two samples from human bone marrow mononuclear cell dataset with batch effects.

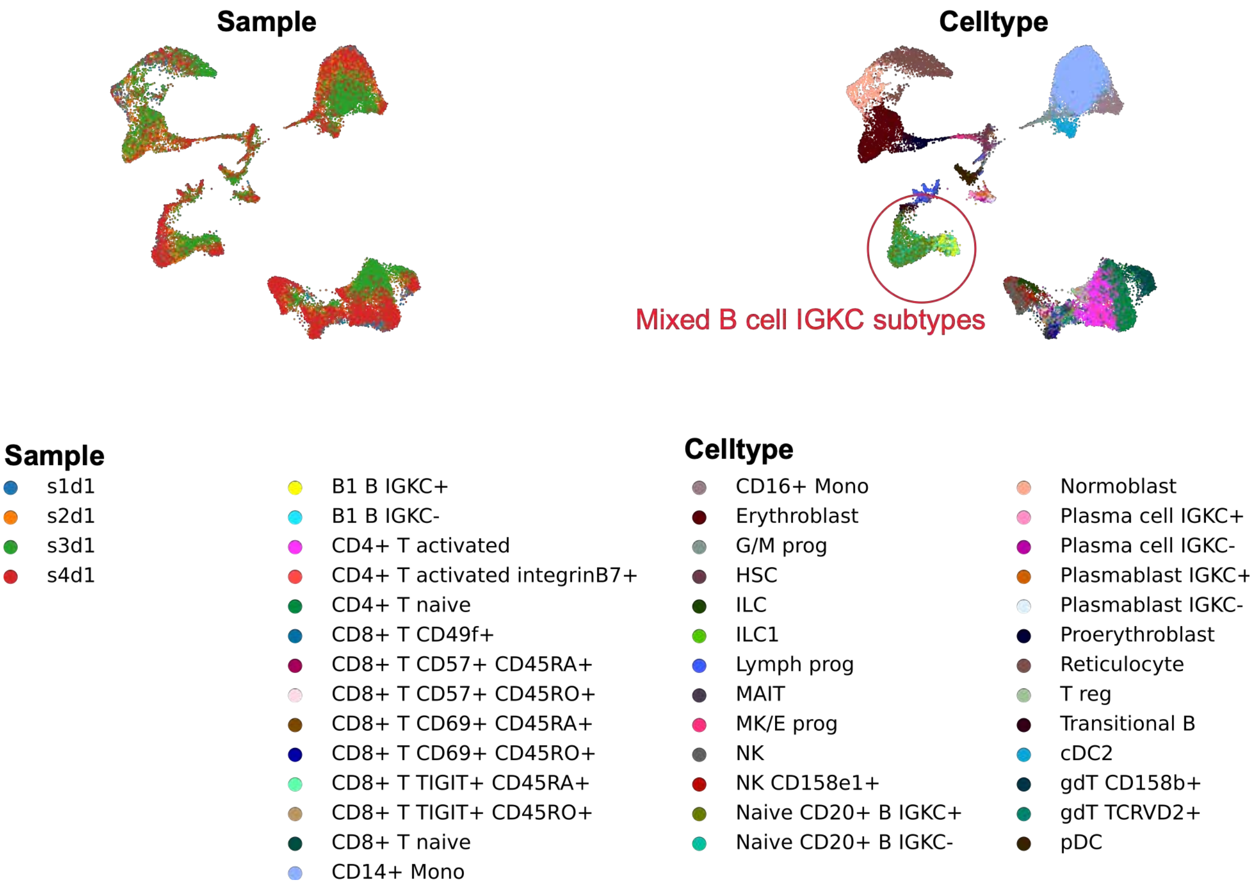


**Fig. S17. UMAP visualizations of integration results on the BMMC dataset generated using BBKNN.**

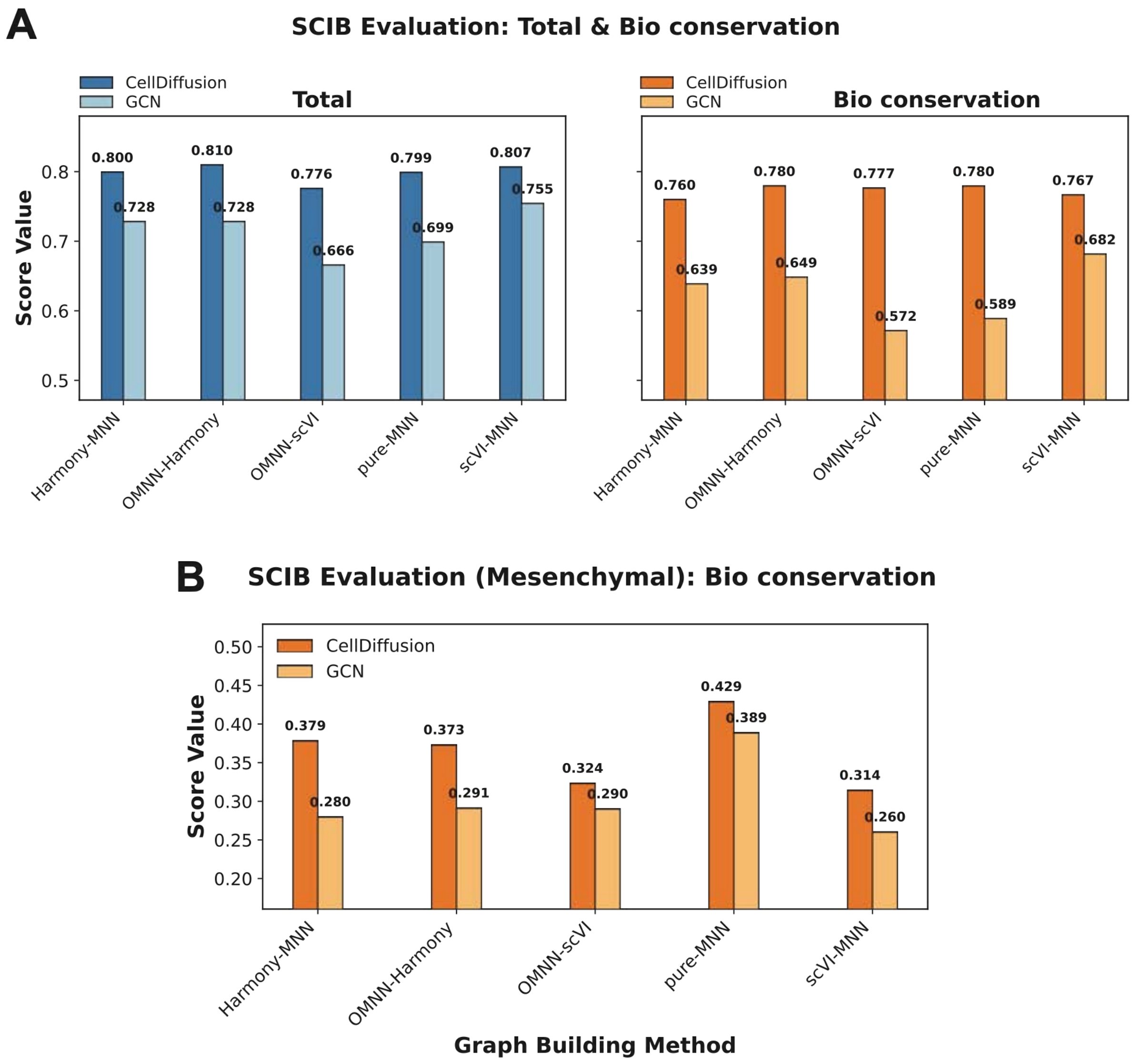


**Fig. S18. Benchmark of CellDiffusion and GCN across different cell graphs.** (**A**) The total integration and biological conservation scores of CellDiffusion and GCN on PBMC-10k dataset across different cell graphs. (**B**) The biological conservation scores of CellDiffusion and GCN on the mesenchymal lineage of the endometrial dataset across different cell graphs.

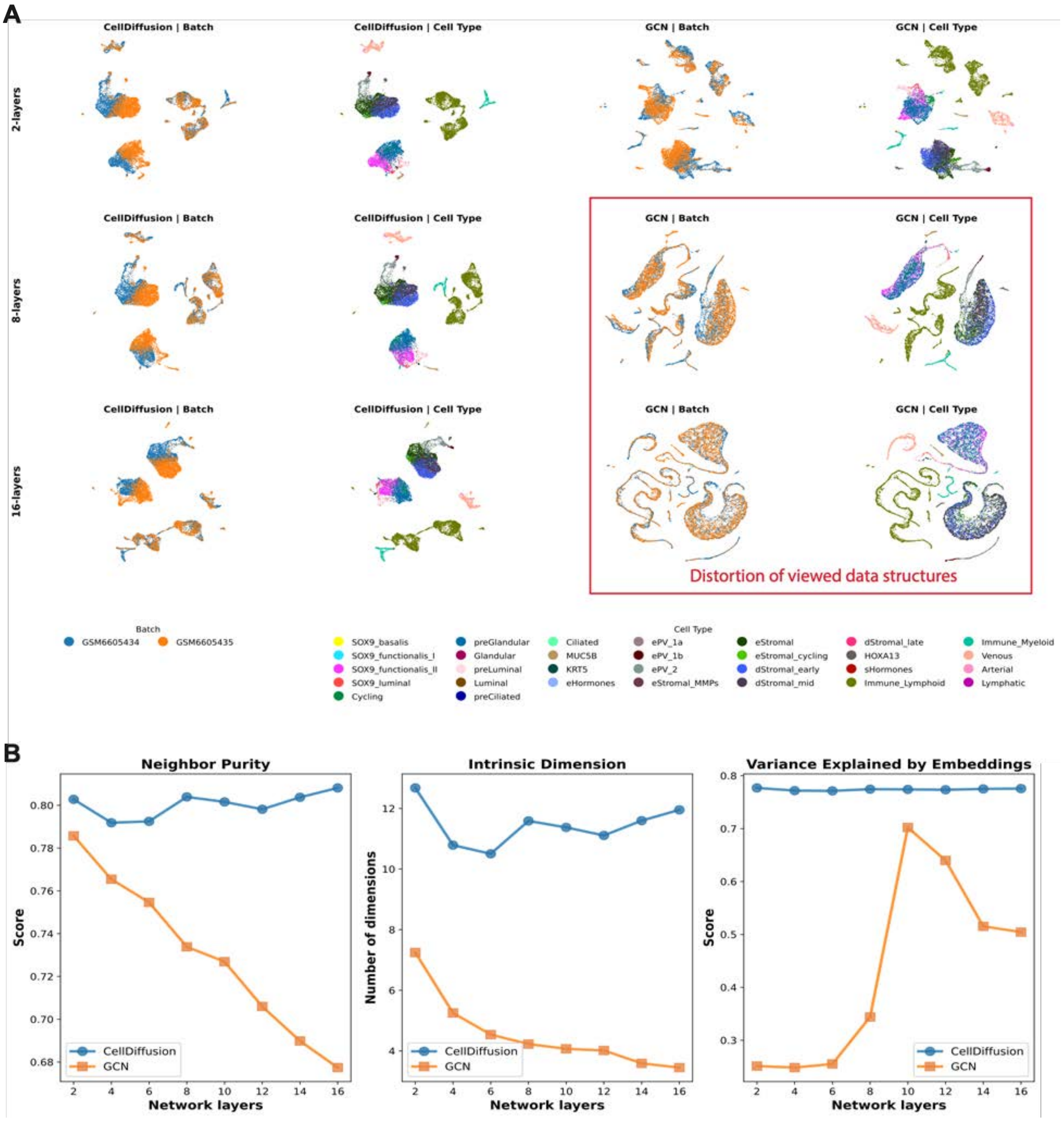


**Fig. S19. Model collapse diagnostics on the Endometrium-CD dataset.** (**A**) UMAP plots of CellDiffusion and GCN with 2, 8 and 16 network layers. (**B**) Evaluation results based on three metrics: neighbor purity, intrinsic dimension and variance explained by embeddings.

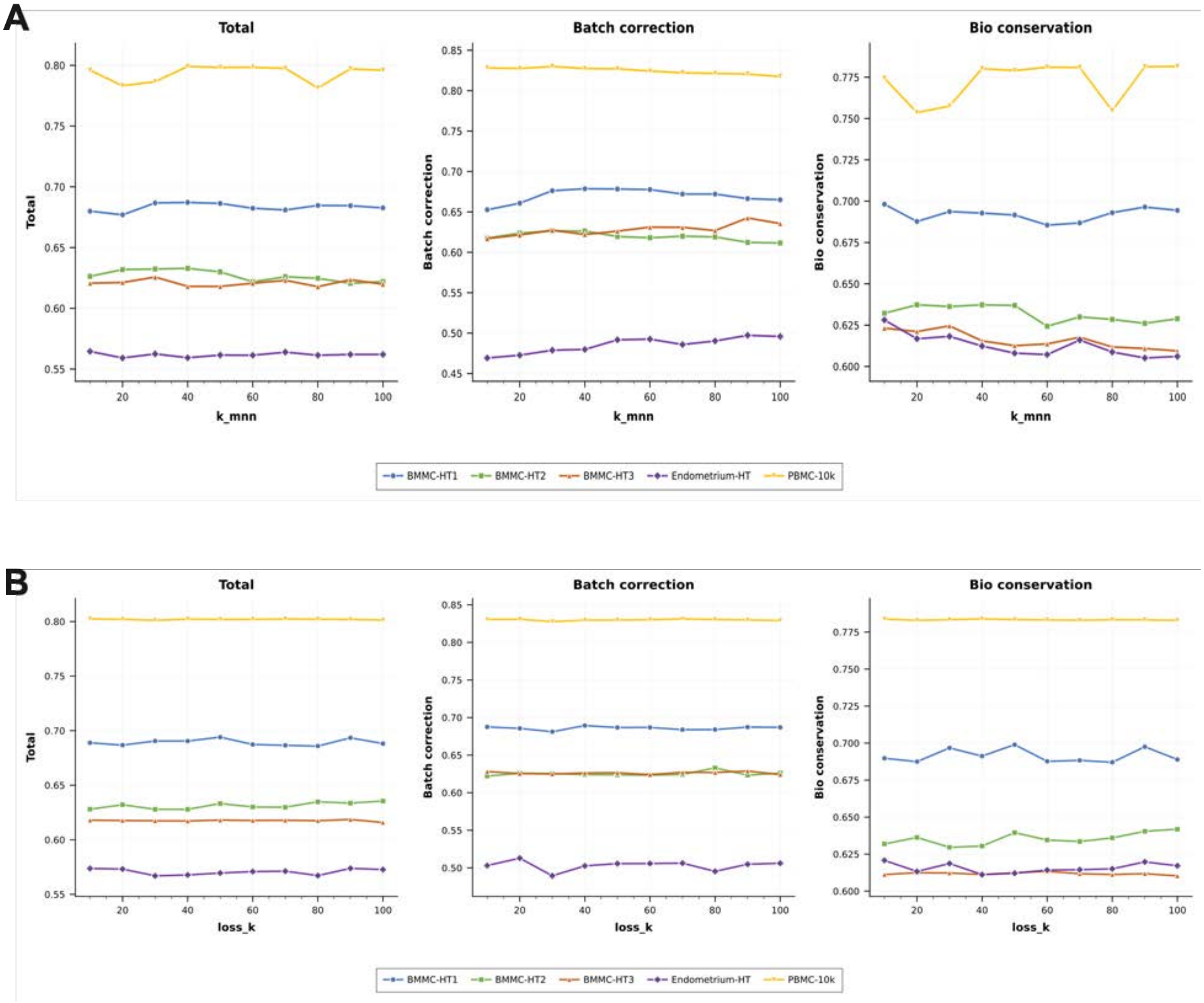


**Fig. S20. Hyperparameter sensitivity analysis of CellDiffusion.** (**A**) Sensitivity analysis at different values of K for KNN to build MNN graphs (k_mnn). The parameter k_mnn determines the edge density of the OMNN graph that used to conduct cell feature diffusion. The data integration results were evaluated using scib-metrics at different k_mnn values. (**B**) Sensitivity analysis at different values of K for KNN to build the batch-KNN graphs that used in the loss function (loss_k).

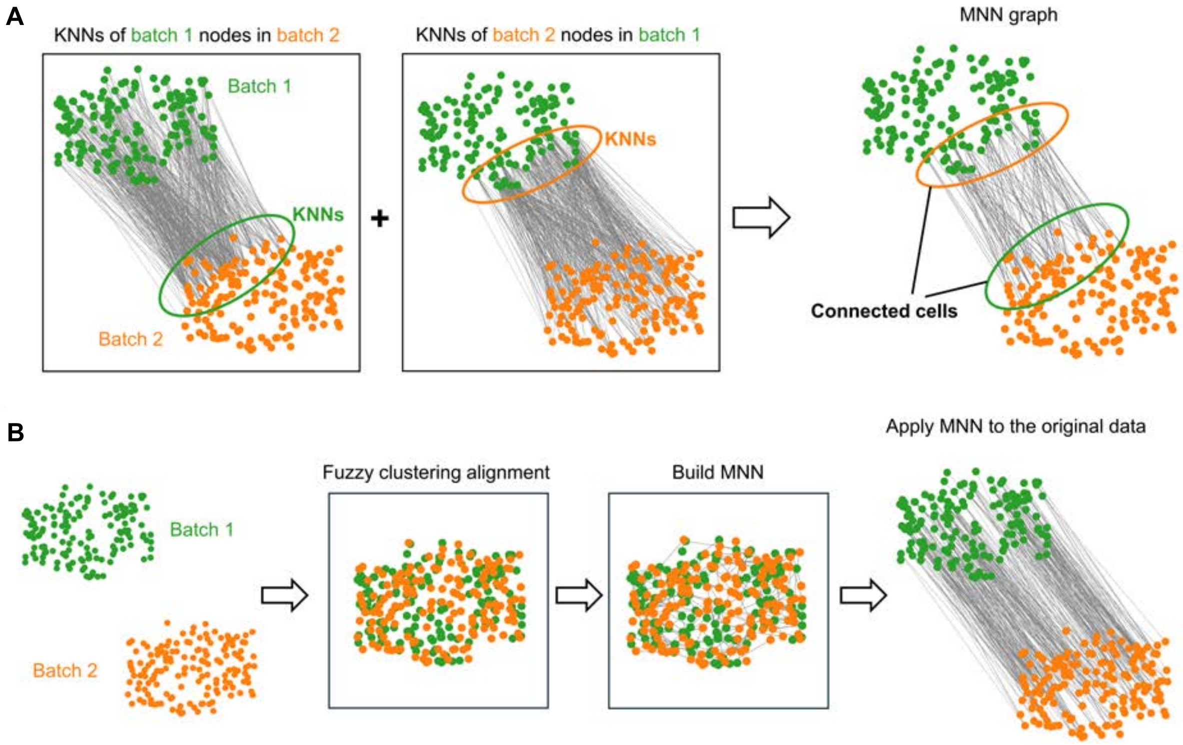


**Fig. S21. MNN and fuzzy clustering alignment.** (**A**) MNN between two batches. We find that nodes in one batch tend to find the same KNNs in other batches because of the short distance. Thus, only a few nodes in each batch closer to another batch can be linked by MNN edges. (**B**) The modified MNN method in CellDiffusion. We use the fuzzy clustering method to preprocess the data before building the MNN graph. The node features are modified by the fuzzy clustering method, and different batches overlap with each other. We build an MNN graph based on the modified data and then apply the MNN graph to the original data.

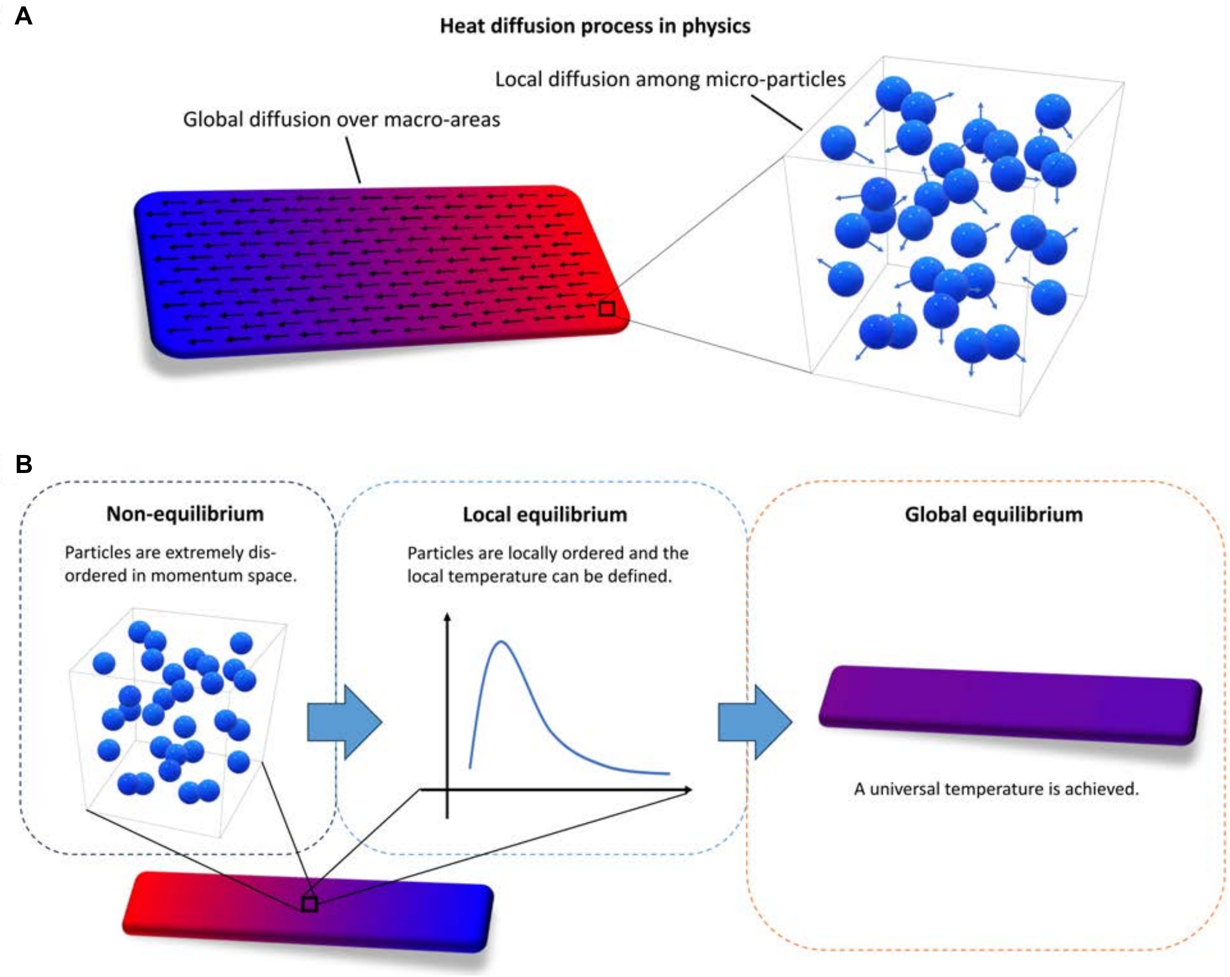


**Fig. S22. The heat diffusion process in physics.** (**A**) The local and global diffusion processes. The local diffusion exchanges energy and momentum between micro-particles inside each localized small quantum element, while the global diffusion transits heat flows over macro-areas. (**B**) Diffusion transforms the non-equilibrium states to the local equilibrium and then the global equilibrium state. In the non-equilibrium state, not only is heat unevenly distributed throughout the system, but micro-particles within each small element are also highly disordered. The localized temperature cannot be defined for the disordered small quantum system. The local diffusion quickly orders micro-particles and leads the system into the local equilibrium, where the local temperature can be well-defined. The global diffusion continues over macro-areas until the universal temperature is achieved, giving rise to the global equilibrium state.

| Dataset | Method | Evaluation Metrics | | | AvgBio |
|---|---|---|---|---|---|
| | | ARI | NMI | FMI | |
| Tabula Muris | CellDiffusion | 0.95 | 0.93 | 0.97 | 0.95 |
| | RaceID | 0.84 | 0.82 | 0.88 | 0.85 |
| | Scanpy | 0.81 | 0.84 | 0.86 | 0.83 |
| | scMoMaT | 0.77 | 0.78 | 0.83 | 0.79 |
| | Seurat | 0.76 | 0.79 | 0.82 | 0.79 |
| | Spectrum | 0.42 | 0.62 | 0.57 | 0.53 |
| | SHARP | 0.34 | 0.63 | 0.54 | 0.50 |
| | scCCESS | 0.20 | 0.30 | 0.50 | 0.33 |
| Cortex | CellDiffusion | 0.87 | 0.82 | 0.90 | 0.86 |
| | Seurat | 0.84 | 0.79 | 0.88 | 0.84 |
| | scMoMaT | 0.84 | 0.78 | 0.88 | 0.83 |
| | Scanpy | 0.72 | 0.70 | 0.78 | 0.73 |
| | SHARP | 0.64 | 0.73 | 0.74 | 0.71 |
| | RaceID | 0.66 | 0.65 | 0.73 | 0.68 |
| | Spectrum | 0.34 | 0.49 | 0.48 | 0.44 |
| | scCCESS | 0.29 | 0.49 | 0.46 | 0.41 |
| PBMC-UC | CellDiffusion | 0.83 | 0.84 | 0.87 | 0.85 |
| | Seurat | 0.79 | 0.82 | 0.84 | 0.82 |
| | Scanpy | 0.58 | 0.74 | 0.72 | 0.68 |
| | SHARP | 0.45 | 0.68 | 0.66 | 0.60 |
| | scMoMaT | 0.51 | 0.62 | 0.62 | 0.58 |
| | RaceID | 0.37 | 0.58 | 0.50 | 0.48 |
| | Spectrum | 0.25 | 0.46 | 0.42 | 0.38 |
| | scCCESS | 0.27 | 0.41 | 0.44 | 0.38 |

**Table S1. scRNA-seq clustering benchmark results.** CellDiffusion was benchmarked with Seurat, Scanpy, scMoMaT, RaceID, Spectrum, SHARP and scCCESS on the Tabula Muris (6 cell types), Cortex (7 cell types), PBMC-UC (8 cell types) datasets for cell type clustering performance. We presented three evaluation metrics: ARI, NMI and FMI, and the average score AvgBio.

| Dataset | Method | Evaluation Metrics | | | AvgTra |
|---|---|---|---|---|---|
| | | Graph edit index | Graph kernel index | Jaccard similarity index | |
| Planaria Full | CellDiffusion | 0.72 | 0.53 | 0.56 | 0.60 |
| | PAGA | 0.66 | 0.47 | 0.49 | 0.54 |
| | Slingshot Root | 0.57 | 0.37 | 0.39 | 0.44 |
| | Slingshot | 0.51 | 0.37 | 0.33 | 0.40 |
| | RaceID | 0.45 | 0.29 | 0.28 | 0.34 |
| Planaria pair | CellDiffusion | 0.85 | 0.76 | 0.73 | 0.78 |
| | Slingshot Root | 0.78 | 0.54 | 0.63 | 0.65 |
| | PAGA | 0.70 | 0.39 | 0.53 | 0.54 |
| | Slingshot | 0.63 | 0.53 | 0.44 | 0.53 |
| | Monocle3 | 0.63 | 0.38 | 0.50 | 0.50 |
| | MST | 0.56 | 0.44 | 0.37 | 0.45 |
| | RaceID | 0.41 | 0.16 | 0.24 | 0.27 |
| Oligodendrocyte | CellDiffusion | 0.83 | 0.76 | 0.69 | 0.76 |
| | Monocle3 | 0.65 | 0.60 | 0.47 | 0.57 |
| | MST | 0.65 | 0.52 | 0.47 | 0.55 |
| | PAGA | 0.57 | 0.47 | 0.38 | 0.47 |
| | Slingshot | 0.57 | 0.47 | 0.38 | 0.47 |
| | Slingshot Root | 0.57 | 0.47 | 0.38 | 0.47 |
| | RaceID | 0.52 | 0.43 | 0.31 | 0.42 |
| Hepatoblast | CellDiffusion | 0.90 | 0.76 | 0.82 | 0.83 |
| | PAGA | 0.81 | 0.74 | 0.67 | 0.74 |
| | MST | 0.71 | 0.69 | 0.58 | 0.66 |
| | Slingshot Root | 0.81 | 0.42 | 0.67 | 0.63 |
| | RaceID | 0.71 | 0.47 | 0.55 | 0.58 |
| | Monocle3 | 0.62 | 0.50 | 0.46 | 0.53 |
| | Slingshot | 0.43 | 0.36 | 0.25 | 0.35 |

**Table S2. scRNA-seq trajectory inference benchmark results.** CellDiffusion was benchmarked with PAGA, Slingshot, Slingshot Root, RaceID, Monocle 3, and MST on the Planaria full (33 cell types), Planaria pair (14 cell types), Oligodendrocyte (12 cell types) and Hepatoblast (11 cell types) datasets for trajectory inference. We presented three evaluation metrics: graph edit index, graph kernel index and Jaccard similarity index, and the average score AvgTra.

| Dataset | N_Cells | N_Batches | CellDiffusion_Time (min) | Harmony_Time (min) | scVI_Time (min) | Seurat_Time (min) |
|---|---|---|---|---|---|---|
| BMMC | 30,669 | 4 | 14.35 | 0.99 | 5.91 | 10.12 |
| PBMC-10K | 11,990 | 2 | 5.25 | 0.57 | 1.34 | 2.1 |
| PBMC-23K | 23,589 | 3 | 10.57 | 0.83 | 4.63 | 6.59 |

**Table S3. Timing benchmark for CellDiffusion, Harmony, scVI and Seurat integrations on three datasets.**

| Dataset | N_Cells | N_Batches | CellDiffusion_GPU (GB) | Harmony_CPU (GB) | scVI_CPU (GB) | Seurat_CPU (GB) |
|---|---|---|---|---|---|---|
| BMMC | 30,669 | 4 | 14.299 | 2.12 | 2.291 | 11.353 |
| PBMC-10K | 11,990 | 2 | 5.387 | 1.271 | 2.069 | 3.38 |
| PBMC-23K | 23,589 | 3 | 10.977 | 1.805 | 2.18 | 13.635 |

**Table S4. Memory benchmark for CellDiffusion, Harmony, scVI and Seurat integrations on three datasets.**
